# Alternative splicing broadens antiviral diversity at the human *OAS2* locus

**DOI:** 10.1101/2025.02.24.639105

**Authors:** Emma L. Davies, Hanna Sowar, Arda Balci, Elliot Moorhouse, Arthur Wickenhagen, Matthew L. Turnbull, Massimo Palmarini, Sam J. Wilson, Adam J. Fletcher

**Author notes:** These authors contributed equally (Hanna Sowar, Arda Balci).

## Abstract

Interferons (IFN) are cytokines that regulate the expression of hundreds of genes during viral infections to generate a broadly antiviral environment in the stimulated cell. Antiviral breadth is provided by the concurrent expression of many individual IFN-stimulated genes (ISG), each encoding a protein with often exquisite antiviral specificity. Here, we show that mechanistic plasticity at a single genetic locus is a novel mechanism to diversify the antiviral profile of human cells. Through alternative splicing, the *OAS2* gene encodes two antiviral molecules with distinct target specificities. The shorter OAS2 p69 isoform blocks the replication of seasonal human coronavirus OC43 (HCoV-OC43), while the longer p71 isoform restricts the replication of picornavirus Cardiovirus A (EMCV). The restriction profile is determined by the variable length OAS2 C-terminal tail. Remarkably, the antiviral mechanisms underlying these distinct antiviral profiles are either RNase L dependent or independent, suggesting that splicing divides ‘classic restriction’ versus ‘virus sensing’ systems across two distinct OAS2 polypeptides. Together, our data reveal that the human *OAS2* locus uses alternative splicing and mechanistic plasticity to diversify antiviral profiles.

## Introduction

Across the tree of life, antiviral defence systems are often based on the recognition of viral RNA (vRNA). In eukaryotes, from cartilaginous fish to birds and mammals, vRNA detection leads to the secretion of cytokines called interferons (IFN) (Dalskov *et al*, 2023; Sa Ribero *et al*, 2020). IFNs regulate the transcription of hundreds of interferon-stimulated or interferon-repressed genes (ISG/IRG), many of which encode proteins with antiviral function (Schoggins, 2019; Shaw *et al*, 2017). The importance of a broad antiviral defence is evidenced by patients with severe viral disease, including herpes simplex encephalitis (HSE) and COVID-19, who harbour defects in components of the IFN pathway, including self-reactive antibodies against IFN or genetic errors in key IFN signalling components like vRNA sensors TLR3 and MDA5 (Bastard *et al*, 2020; Hambleton *et al*, 2013; Lamborn *et al*, 2017; Zhang *et al*, 2007).

The causative agent of COVID-19, SARS-CoV-2, is the third coronavirus to emerge into the human population since the turn of the century, along with SARS-CoV-1 and MERS-CoV (Drosten *et al*, 2003; Zaki *et al*, 2012; Zhu *et al*, 2020). Owing to the recent spillover, these emerging human coronaviruses can be highly pathogenic and cause more severe, lower respiratory tract infections than their seasonal counterparts (HCoV-229E, HCoV-NL63, HCoV-HKU1 and HCoV-OC43), which mainly cause common cold-like symptoms (Gaunt *et al*, 2010). HCoV-OC43 is thought to have diverged from bovine coronavirus (BCoV) in the late 19th century, when it jumped into humans via an intermediate livestock species (Shaw & Gatherer, 2023; Vijgen *et al*, 2005). Based on contemporary events, it is likely that HCoV-OC43 caused an epidemic upon its emergence but later evolved into an endemic respiratory virus (Gaunt *et al*., 2010). Understanding the cellular factors that regulate HCoV-OC43 replication in humans could contribute to our understanding of viral adaptation in new hosts.

To search for these factors, we employed large-scale cDNA screens and uncovered antiviral activity of 2’-5’ oligoadenylate (2-5A) synthetase 2 (OAS2) against the seasonal coronavirus HCoV-OC43. Of the two major OAS2 isoforms sharing the same enzymatic core, only the shorter p69 isoform is antiviral toward HCoV-OC43. Conversely, we find that the longer p71 isoform restricts picornavirus Cardiovirus A (encephalomyocarditis virus (EMCV)). While p71 restriction of EMCV occurs via the classic OAS/RNase L pathway, p69 restriction of HCoV-OC43 is entirely RNase L-independent. Our data demonstrates that alterations in the OAS2 C-terminal tail fine-tunes target specificity, providing a novel mechanism to achieve antiviral diversity from a single genetic locus.

## Results

### Developing a high throughput assay for dsRNA formed during HCoV-OC43 replication

Unlike HCoV-229E and SARS-CoV-2, which originated in bats (Latinne *et al*, 2024), rodents are thought to be the ancestral host of HCoV-OC43 (Corman *et al*, 2018), and there has been little research on genes with antiviral activity against it. Like other positive-sense single-stranded RNA viruses, coronaviruses replicate their genomes via dsRNA replication intermediates (V’kovski *et al*, 2021). Since dsRNA is a potent immune agonist, endogenous dsRNA is typically expressed at low levels (Chen & Hur, 2022). A recent coronavirus screen monitored virus-derived dsRNA to score viral replication (Schneider *et al*, 2021), which we adapted here for HCoV-OC43 and plate-based image cytometry (**Supplementary Fig. 1a,c**). Benefits of this system include its high-throughput nature and utility where neither a recombinant viral clone nor viral antigen-specific antibodies are available. We titrated HCoV-OC43 on A549 lung adenocarcinoma cells, fixing cells at intervals post infection. An increase in signal over time confirmed that detected dsRNA was a robust correlate of viral infection (**Fig. 1a**). Pre-treatment of A549 cells with type I IFN substantially reduced the dsRNA signal, suggesting that these cells provide a suitable context in which to study anti-HCoV-OC43 machinery (**Supplementary Fig. 1b**).

**Figure 1.**
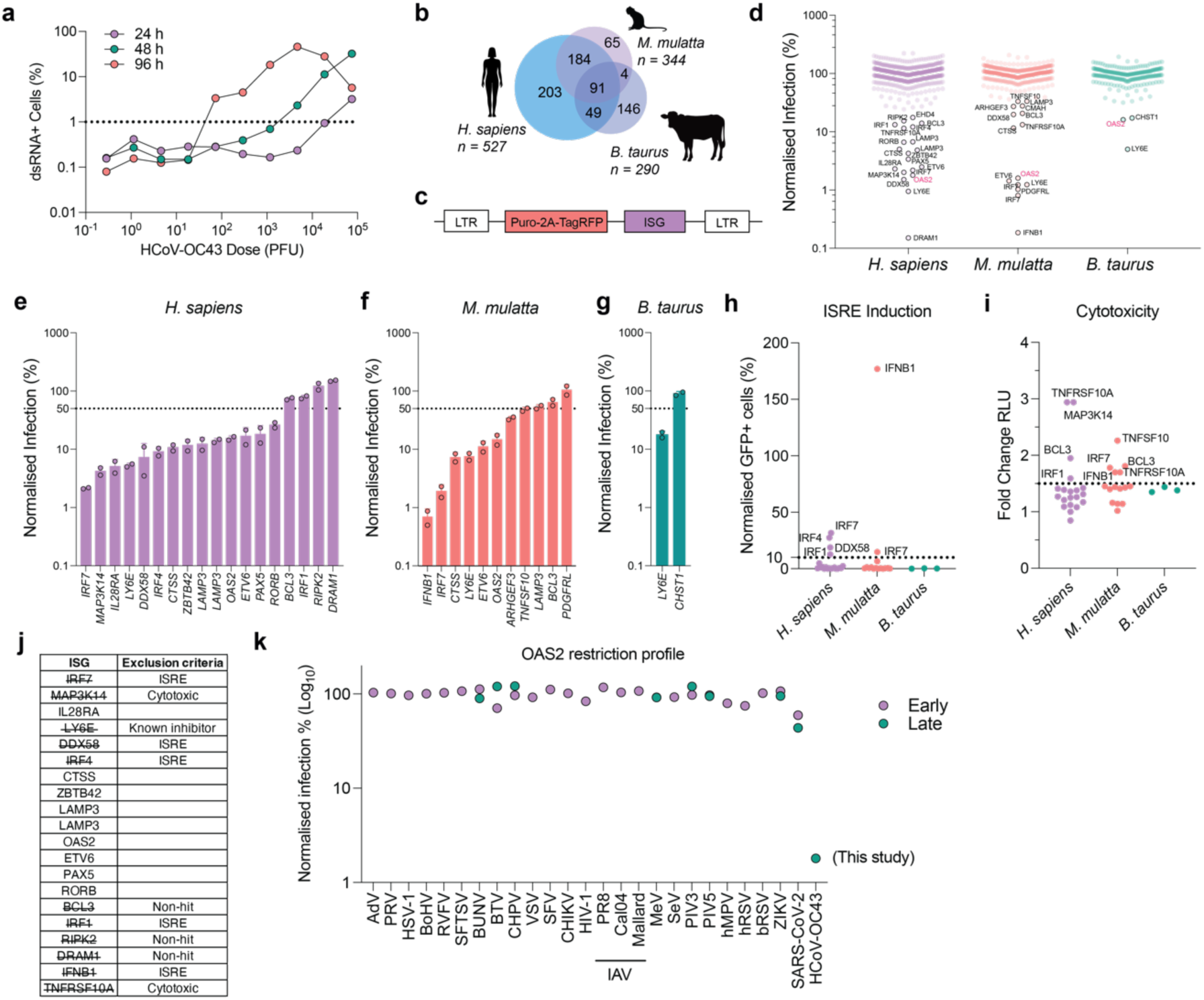
– Identification of genes with antiviral activity against HCoV-OC43. **a)** A549 cells were infected with serially diluted HCoV-OC43. The proportion of dsRNA+ cells at multiple time points was quantified in fixed cells by immunostaining for dsRNA followed by image cytometry. **b)** Schematic of the human (*H. sapiens*), macaque (*M. mulatta*) and bovine (*B. taurus*) ISG libraries used in this study. **c)** Schematic of the SCRPSY lentiviral vector, encoding an individual ISG and TagRFP. **d)** A549 cells were transduced with hundreds of ISGs, listed in **Supplementary Table 1** (Fig. 1b), infected with HCoV-OC43 for 72 h, immunostained for dsRNA, and the level of infection quantified using image cytometry. Infection was normalised to the mean of the species library. **e-g)** Miniscreens were performed to validate the ability of (**e**) human, (**f**) macaque or (**g**) bovine ISGs identified in **d)** to restrict HCoV-OC43 at 72 hpi. The LAMP3 gene is duplicated in the human ISG library. Shown is the mean of duplicate experiments. **h)** The ability of candidate ISGs to stimulate the ISRE in A549-ISRE-GFP cells, assessing GFP expression by flow cytometry 120 h post transduction. **i)** The ability of candidate ISGs to cause cytotoxicity, determined using the CytoTox-Glo™ assay using supernatant harvested from the A549-ISRE-GFP cells in **h)**. **j)** Shortlist of human ISGs that are candidate restriction factors of HCoV-OC43. **k)** Normalised infection levels in cells expressing the p69 isoform for a variety of viruses, from 34 arrayed ISG expression screening datasets.

### Identification of genes with antiviral activity against HCoV-OC43

To search for genes with anti-HCoV-OC43 activity, we undertook a multi-species cDNA screen by transducing A549 cells with ISG libraries encoding human (*Homo sapiens*), rhesus macaque (*Macaca mulatta*) and bovine (*Bos taurus*) ISGs, encoded in lentiviral vectors arrayed in 96-well format, with one ISG expressed per well (**Supplementary Table 1**). Combined, these libraries encoded >500 human genes, >300 macaque genes and >250 cow genes (**Fig. 1b**, **Supplementary Table 1**). In addition to an ISG, the lentiviral vector expresses the fluorescent protein TagRFP as an independent ORF (**Fig. 1c**), to allow quantification of transduction efficiency (**Supplementary Fig. 1a**).

After library transduction, we infected cells in parallel with a single dose of HCoV-OC43, determined previously to infect 30-50% of cells by 96 h post infection (hpi) (**Fig. 1a**). Infection was scored in each well and presented here as relative to the mean of the species library (**Fig. 1d**). The antiviral activity of candidate human, macaque and cow ISGs were re-tested in secondary screens, using independently prepared ISG lentiviruses, newly transduced cells, and repeat HCoV-OC43 infections (**Fig. 1e-g**). The efficacy of our approach was reinforced by the identification of the known antiviral protein LY6E across all three libraries. LY6E inhibits the replication of multiple coronaviruses, including HCoV-OC43, by interfering with membrane fusion at virus entry (Pfaender *et al*, 2020).

Other antiviral candidates from our screen included the endosomal protease cathepsin S (CTSS), the vRNA sensors OAS2 and DDX58/RIG-I, and the transcription factors ETV6 and IRF7. CTSS was previously identified in a CRISPR-activation screen against SARS-CoV-2, suggesting broad anti-CoV behaviour (Danziger *et al*, 2022). Cathepsins B (CTSB) and L (CTSL) proteolyse SARS-CoV-2 Spike protein, promoting virus entry in endosomes (Jackson *et al*, 2022), suggesting that CTSS overexpression may modulate HCoV-OC43 entry pathways. OAS2 has appeared in several genetic screens against CoVs (Mac Kain *et al*, 2022; Pfaender *et al*., 2020; Wickenhagen *et al*, 2021), however its anti-CoV activity has not been well characterised. RIG-I is a key cellular sensor of vRNA and several reports also describe a role for RIG-I in sensing SARS-CoV-2 RNA (Marx *et al*, 2022; Thorne *et al*, 2021). ETV6 is poorly studied but the related protein ETV7 has been shown to regulate expression of ISGs that inhibit influenza A virus (Froggatt *et al*, 2021). IRF7 is a key transcription factor driving IFN signalling; IRF7 overexpression might be expected to inhibit viral infections via non-specific ISG induction.

To distinguish effectors acting directly vs indirectly on HCoV-OC43 infection, we next screened our candidate ISGs for their ability to induce an IFN response independently of viral infection. To do this, we expressed each ISG in A549 cells containing an interferon-stimulated response element (ISRE)-GFP reporter construct, whereby GFP expression is a proxy for IFN pathway activity. In doing so, we found that IRF1, IRF4, IRF7, IFNβ1 and RIG-I all caused >10-fold ISRE activation (**Fig. 1h**). We also used a cytotoxicity assay, which pinpointed TNFRSF10A and MAP3K14 as reducing cell viability, likely explaining the antiviral effects detected (**Fig. 1i**). After removing off-target genes, we could shortlist our putative anti-HCoV-OC43 ISGs (**Fig. 1j**).

Of these, we were most intrigued by OAS2. OAS enzymes synthesise short 2’-5’ phosphodiester-linked polyA (oligoadenylate) RNAs (2-5A) upon binding dsRNA (Marié *et al*, 1997). 2-5A behaves as a second messenger, binding the dormant endoribonuclease RNase L, inducing its dimerisation and activation (Dong & Silverman, 1995). Once unleashed, RNase L degrades host and viral RNAs, stemming viral replication whilst inhibiting cell proliferation (Ghosh *et al*, 2000; Maitra *et al*, 1995). The human OAS gene family (OAS1, OAS2, OAS3 and OASL) undergoes alternative splicing, generating multiple isoforms for *OAS1* (p42, p44, p46, p48 and p52), *OAS2* (p69 and p71) and *OASL* (p30 and p56-59) (Koul *et al*, 2024). Interestingly, HCoV-OC43 encodes a 2’-5’-phosphodiesterase (PDE) in its *ns2* gene, a described antagonist of the RNase-L pathway via 2-5A hydrolysis (Goldstein *et al*, 2017). This viral countermeasure was thought to explain the resistance of HCoV-OC43 to OAS1, which potently inhibits SARS-CoV-2 (Wickenhagen *et al*., 2021). We were also surprised to identify

OAS2 because in 33 additional ISG screens at the Centre for Virus Research, against diverse RNA viral families including *Lentiviridae*, *Togaviridae*, *Peribunyaviridae*, *Orthomyxoviridae* and *Coronaviridae*, OAS2 is notable in having only been identified as a putative antiviral in one (SARS-CoV-2), and in that example, with only very modest antiviral activity (1.7-2.1-fold inhibition) (**Fig. 1k**) (Wickenhagen *et al*., 2021). Combined, this suggested that by studying this specific combination of virus and host enzymes, we might uncover novel antiviral biology.

### The p69 isoform of OAS2 shows antiviral activity against HCoV-OC43

Our human ISG library only contained the p69 isoform, whereas the literature generally references two OAS2 transcripts, namely p69 and p71 (Koul *et al*., 2024; Kristiansen *et al*, 2010; Marié *et al*, 1999; Schwartz & Conn, 2019; Silverman, 2007). To investigate whether other OAS2 isoforms are also restrictive toward HCoV-OC43, we sought a panel of OAS2 transcripts to test. NCBI lists three OAS2 transcripts, while Ensembl lists fifteen, seven of which are predicted to encode protein. To corroborate this larger Ensembl dataset, we first referred to the Genotype-Tissue Expression project (GTEx), which quantifies Ensembl transcript abundance across 54 tissues (Consortium, 2015). Despite Ensembl’s large collection of transcripts, only three transcripts were significantly expressed across a range of tissues (although notably these sequencing data are from donors rather than IFN-treated cell cultures, so potentially underestimate the transcript repertoire found during an immune response). These corresponded with the NCBI annotated transcripts – ENST00000392583.7, p69 (NM_002535.3), ENST00000342315.8, p71 (NM_016817.3) and ENST00000449768.2, transcript 3, isoform 3 (NM_001032731.2) (**Fig. 2a**). The p69 and p71 transcripts encode the same core 683 residues but owing to an alternate splicing event near to the end of exon 10, an additional exon (exon 11) is formed, which includes a short in-frame coding sequence and stop codon, to give p71. Consequently, while p69 has a 4-residue C-terminal tail, p71 has a longer, 36-residue tail (**Fig. 2b**). Conversely, isoform 3 shares the first 150 residues with p69 and p71 but by missing a splice site, has an extended exon 2 resulting in a unique 22-residue tail. Isoform 3 lacks key residues required for RNA binding and 2-5A synthesis, suggesting its role might be regulatory rather than the detection of viral RNA. Moreover, while the PeptideAtlas project (Desiere *et al*, 2006) provided empirical evidence for the expression of both p69 and p71 isoforms in diverse cell types, there are no recorded peptides unique for isoform 3, suggesting this isoform might not even be expressed (**Supplementary Fig. 2a**). Thus, we decided to exclude OAS2 isoform 3 from further investigation.

**Figure 2.**
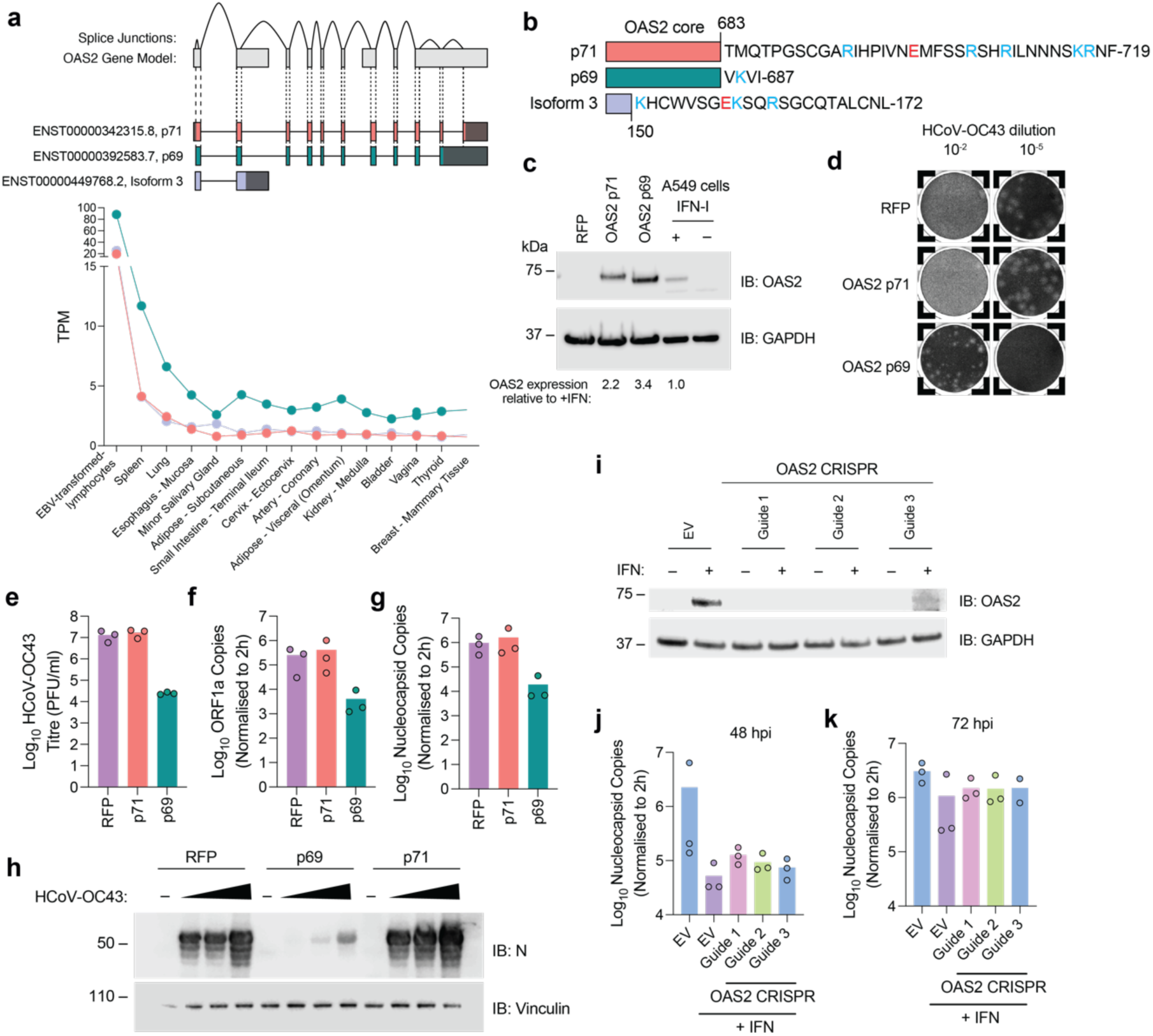
– The p69 isoform of human OAS2 restricts HCoV-OC43 replication. **a)** Representation of the OAS2 gene model including exon structures and splice junctions. The three major OAS2 transcripts are indicated, shaded boxes are non-coding, bright colouring indicates coding sequences. Transcript expression across tissues analysed using the GTEx database. **b)** Schematic of the OAS2 isoforms with the distinct C-termini sequences of p69 (NM_002535.5), p71 (NM_016817.3), and isoform 3 (NM_001032731.2) indicated. Basic residues are in blue, and acidic residues are in red. **c)** A549 cells were modified to express RFP, OAS2-p71 or OAS2-p69, confirmed by Western blotting. A549 cells treated with 1000 U/mL IFNβ are additionally shown. **d)** Representative images of HCoV-OC43 plaques formed in cell lines characterised in (**c**), 120 hpi. **e)** Infectious titre of HCoV-OC43 in cell lines generated in (**c**) were quantified by plaque assay at 120 hpi. **f)** Quantification of HCoV-OC43 *ORF1a* transcripts in OAS2-expressing cells infected for 72 h (MOI 0.01) by RT-qPCR. **g)** Quantification of HCoV-OC43 *nucleocapsid* (*N*) transcripts in OAS2-expressing cells infected for 72 h (MOI 0.01) by RT-qPCR. **h)** N protein levels in A549 cells expressing OAS2 p69 or p71, or an RFP control, were compared by Western blotting, following infection with HCoV-OC43 (MOI of 0.005, 0.01 and 0.05) for 72 h (n=1). **i)** Endogenous OAS2 levels in A549 cells transduced with lentiviral vector-derived OAS2 CRISPR guides, with or without pre-treatment of 100 U/mL IFNβ for 24 h. **j,k)** HCoV-OC43 *nucleocapsid* levels were measured in OAS2-depleted A549 cells stimulated with 100 U/mL IFNβ for 24 h and then infected with HCoV-OC43 (MOI 0.01) for 48 h (**j**) and 72 h (**k**), by RT-qPCR. **(e-g, j-k)** Data shown are the mean of 3 biological replicates.

We next stably expressed the p69 or p71 isoforms, or an RFP control, in A549 cells – Western blots confirmed expression, and the isoforms exhibited modest differential electrophoretic mobility (**Fig. 2c**). Parallel electrophoresis of lysates from A549 cells stimulated with IFN, suggested that p69 was the predominant endogenous isoform expressed in these cells (**Fig. 2c**). We next infected our OAS2 overexpression cells with HCoV-OC43 and monitored viral replication 120 hpi by plaque assay. Plaques in p69-expressing cells were smaller and more heterogenous than those formed in RFP-expressing cells (**Fig. 2d**). Strikingly, the p69 isoform displayed ∼500-fold inhibition of HCoV-OC43 replication across three biological replicates, whereas p71 showed no restriction of viral replication (**Fig. 2e**). To confirm antiviral activity, we developed a qPCR assay to quantify HCoV-OC43 replication using probes targeting either the 5’ end of the viral genome (within ORF1a/b) or the 3’ subgenomic region (within the ORF encoding N). Levels of these transcripts were examined over 72 h; ORF1a transcripts peaked at 48 hpi at all MOIs tested (**Supplementary Fig. 2b**), whereas nucleocapsid transcripts increased continuously at all MOIs tested (**Supplementary Fig. 2c**). At 72 hpi, we observed a 60-fold and 50-fold decrease in genomic (ORF1a/b) and genomic+subgenomic (N) transcripts, respectively, in p69-expressing cells compared to control cells (**Fig. 2f,g**). We also monitored HCoV-OC43 replication by immunoblotting for nucleocapsid (N) protein 72 hpi, which we determined was an optimal timepoint to detect abundant N protein in soluble lysates at multiple input doses of virus. N expression in p69-expressing cells was substantially reduced at all doses of virus, when compared to RFP-expressing or p71-expressing cells (**Fig. 2h**). Together, these data reveal a potent antiviral activity of OAS2 p69, but not p71, against HCoV-OC43 at the transcript, protein, and infectious particle levels.

### Endogenous OAS2 contributes to IFN-mediated suppression of HCoV-OC43 replication

The antiviral effect of IFNs against any single virus can be the cumulative activity of a small subset of ISGs (McDougal *et al*, 2023). A549 cells express the restrictive p69 isoform upon IFN stimulation (**Fig. 2b**), so we hypothesised that OAS2 might contribute to IFN’s inhibition of HCoV-OC43 in this setting (**Supplementary Fig. 1b**). We used CRISPR/Cas9 to reduce OAS2 expression in A549 cells, confirming depletions by Western blot (**Fig. 2i**). We then pre-treated control or OAS2-depleted cells with IFN before infecting with HCoV-OC43 and quantifying viral nucleocapsid transcripts at 48 hpi. We noted that IFN reduced HCoV-OC43 plaque sizes significantly, so we used qPCR to accurately quantify viral replication. In control cells, IFN induced ∼40-fold inhibition in N transcripts, whereas in OAS2-depleted cells, this inhibition was reduced by 1.4-to 2.4-fold (**Fig. 2j**). At 72 hpi, HCoV-OC43 exhibited ∼1.4-fold less sensitivity to IFN in all three OAS2 depleted cell lines, compared to the controls (**Fig. 2k**). The modest effect size was unsurprising, because removing one ISG often has minimal effects on IFN-mediated virus restriction, whereas synchronously removing 2-3 ISGs can restore viral replication completely (McDougal *et al*., 2023). Moreover, we note that depletion of related enzyme OAS1, in some contexts, has no effect on the replication of SARS-CoV-2 *in vitro* but has a substantial effect on SARS-CoV-2 replication *in vivo* (Lee *et al*, 2023; Wickenhagen *et al*., 2021). Nonetheless, these experiments support a contributory role for endogenous OAS2 in IFN-mediated restriction of HCoV-OC43 in A549 cells.

### N-myristoylation is necessary for OAS2 p69 antiviral activity

The p46 isoform of OAS2-related enzyme OAS1 requires anchoring to intracellular membranes to sense coronavirus and picornavirus replication, achieved via post-translational prenylation at its C-terminal CaaX motif (Soveg *et al*, 2021; Wickenhagen *et al*., 2021). Early characterisation of OAS2 p69 showed the protein was N-myristoylated *in vitro*, and this was hypothesised to facilitate its association with membranes (Marié *et al*, 1990). N-myristoylation is a distinct, co-translational lipid modification catalysed by the enzyme N-myristoyltransferase (NMT) and can direct protein membrane targeting (**Fig. 3a**) (Tate *et al*, 2024). Certain features of the OAS2 N-terminal peptide, including a glycine immediately after the initiator methionine, and a serine at position 5 (**Fig. 3a**), make OAS2 an ideal NMT substrate (Johnson *et al*, 1994). We first investigated the subcellular appearance of endogenous OAS2 using A549 cells treated with or without type I IFN for 24 h, which induces expression of the p69 isoform (**Fig. 2b**). OAS2 formed a reticular pattern that did not localise with ER marker calnexin (**Fig. 3b, Supplementary Fig. 3a**) but did with the Golgi marker formiminotransferase cyclodeaminase FTCD/58K (Bloom & Brashear, 1989) (**Fig. 3b,c**). This distribution was remarkably similar to that of the ectopic OAS2 isoforms (**Fig. 3b, Supplementary Fig. 3a**). To investigate a role for lipidation in OAS2 function, we substituted glycine at position 2 with alanine, which inhibits lipidation by NMT (Borgese *et al*, 1996; Deichaite *et al*, 1988), and stably expressed these proteins in A549 cells (**Fig. 3d**). In contrast to their WT counterparts, G2A variant OAS2 proteins were diffusely expressed and lost specific Golgi localisation (**Fig. 3b**), supporting a role for N-myristoylation in OAS2 membrane association.

**Figure 3.**
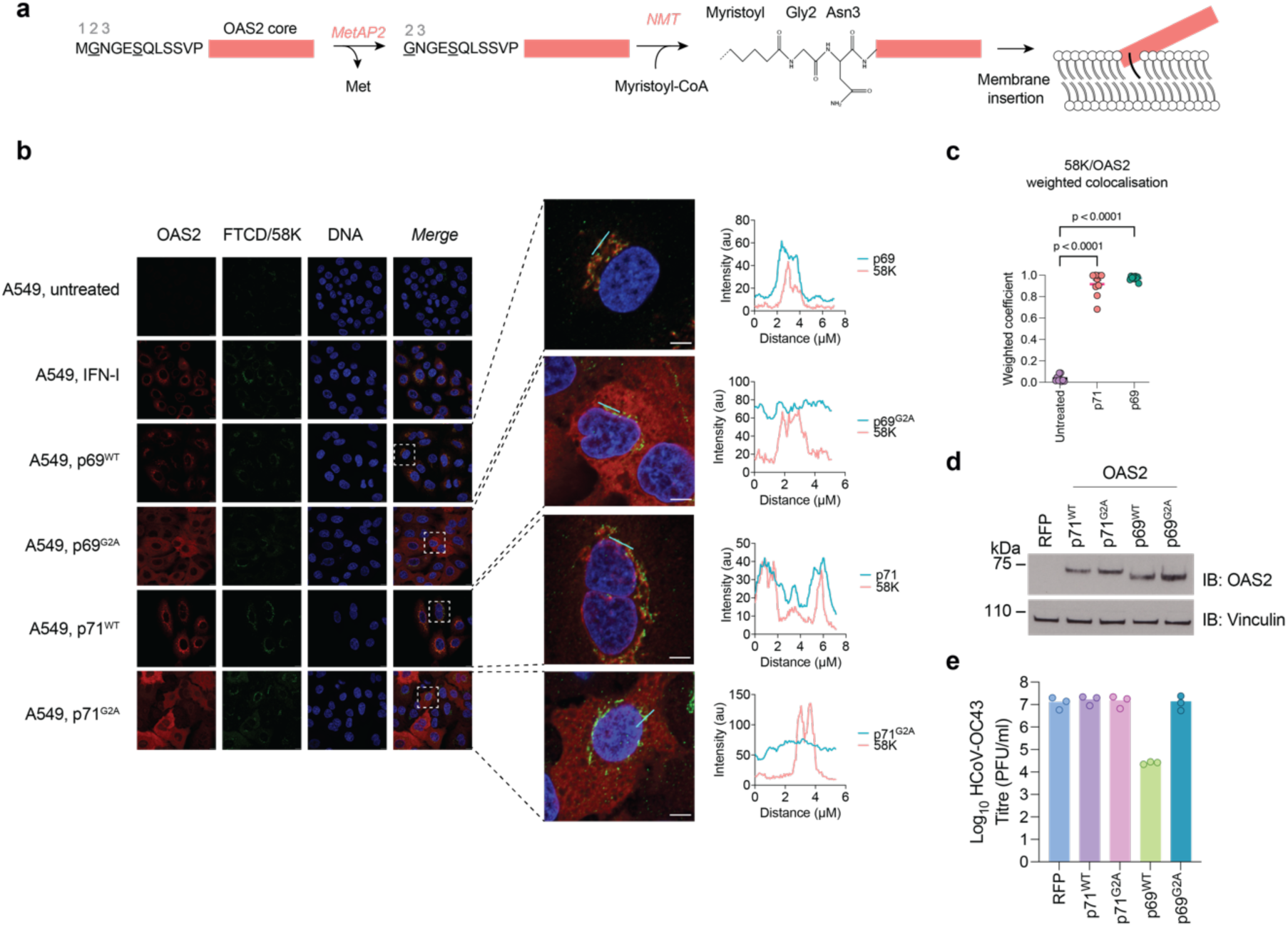
– Myristoylation is required for the antiviral activity of OAS2 p69. **a)** N-myristoylation of OAS2 proteins. The initiator methionine is removed by methionine aminopeptidase 2 (MetAP2) prior to N-myristoyltransferase (NMT) catalysing the addition of the myristoyl group to the glycine residue at the N-terminus. **b)** A549 cells stimulated with 1000 U/mL IFNβ, or A549 cells expressing p71, p69, p71^G2A^ or p69^G2A^, were immunostained for OAS2 (red), anti-FTCD/58K (green) and Hoechst (blue). Signal intensity profiles along the lines of magnified boxes. **c)** Weighted coefficients between p71 and p69 isoforms and Golgi marker 58K; a one-way ANOVA test was used to assess significance, threshold p = 0.05. Each data point represents a separate region of interest from a representative experiment. **d)** A549 cells were modified to express the OAS2 isoforms indicated, confirmed by Western blotting. **e)** The infectious titre of HCoV-OC43 in cell lines generated in **d)** were determined by plaque assay at 120 hpi; data shown are the mean of 3 biological replicates.

To look for OAS2-dsRNA interaction in cells, we infected A549 expressing p69 or p71 with HCoV-OC43 for 24 h and stained cells with the same dsRNA-specific antibody used in our initial screen. In this way, we observed that both p69 and p71 colocalised with dsRNA, although only the correlation for p71 achieved statistical significance (p < 0.05), perhaps indicative of aborted viral replication in the presence of p69 (**Supplementary Fig. 3b,c**). To test whether OAS2 N-myristoylation contributes to viral restriction, we infected A549 cells expressing the WT or G2A-substituted p69 or p71 variants with HCoV-OC43 and measured viral replication by plaque assay. As before, p69 robustly inhibited HCoV-OC43 replication, but this was completely reversed by G2A substitution (**Fig. 3e**). Importantly, *in vitro* studies demonstrate that p69 myristoylation is not necessary for enzymatic activity *per se* (Sarkar *et al*, 1999a). Thus, these data support a model where myristoylation facilitates OAS2 recruitment to membranes proximal to HCoV-OC43 replication compartments, where it can bind viral RNA and inhibit viral replication.

### Residues involved in RNA binding are necessary for antiviral activity of OAS2 p69

To test that interaction with RNA is necessary for the observed restriction activity, we sought to reduce p69 binding to viral RNA. In the absence of an experimentally determined OAS2-dsRNA structure, we used the AlphaFold3 (AF3) Server to generate a series of structural predictions (Abramson *et al*, 2024). As RNA ligands, we used two experimentally optimised ligands: one copy of a palindromic 43mer that activated OAS2 *in vitro* (Koul *et al*, 2020a), or two copies of an 18mer that co-crystallised with OAS1 (GGCUUUUGACCUUUAUGC), repeated once (2×36mer) (Donovan *et al*, 2013). We reasoned that comparing complexes with these two RNAs could help us dissect OAS2-RNA binding. OAS2 forms a stable homodimer in solution (Koul *et al*, 2020b; Sarkar *et al*., 1999a) so we included two copies of OAS2 in all predictions. Confidence metrics of the individual OAS2 protomers were relatively high (pLDDT >70) (**Fig. 4a,b**), although the relative arrangement of opposing molecules was variable, as revealed by superposition of all 5 AF3 structures, as well as predicted aligned error (PAE) plots of the top-ranked structures (**Supplementary Fig. 4a,b**). In contrast, expected position error values between OAS2 monomers and RNA were between 0-5 Å, suggesting higher confidence, particularly for the complex with 2×36mer RNA (**Supplementary Fig. 4a**). In both structures, RNA was sandwiched between antiparallel OAS2 protomers. The singlet palindromic 43mer appeared as a hairpin, contacting only the catalytic OAS domains (DII), while the helical 2×36mer appeared as a linear duplex running lengthways along the surface of both OAS domains (DI and DII) (**Fig. 4a,b**). Despite OAS2 protomers arranged in different relative conformations in the two complexes, lysine and arginine residues within 5 Å of nucleic acid were highly consistent between the two complexes (**Supplementary Fig. 4c**), forming an extended basic channel through which the RNA lay (**Supplementary Fig. 4d**).

**Figure 4.**
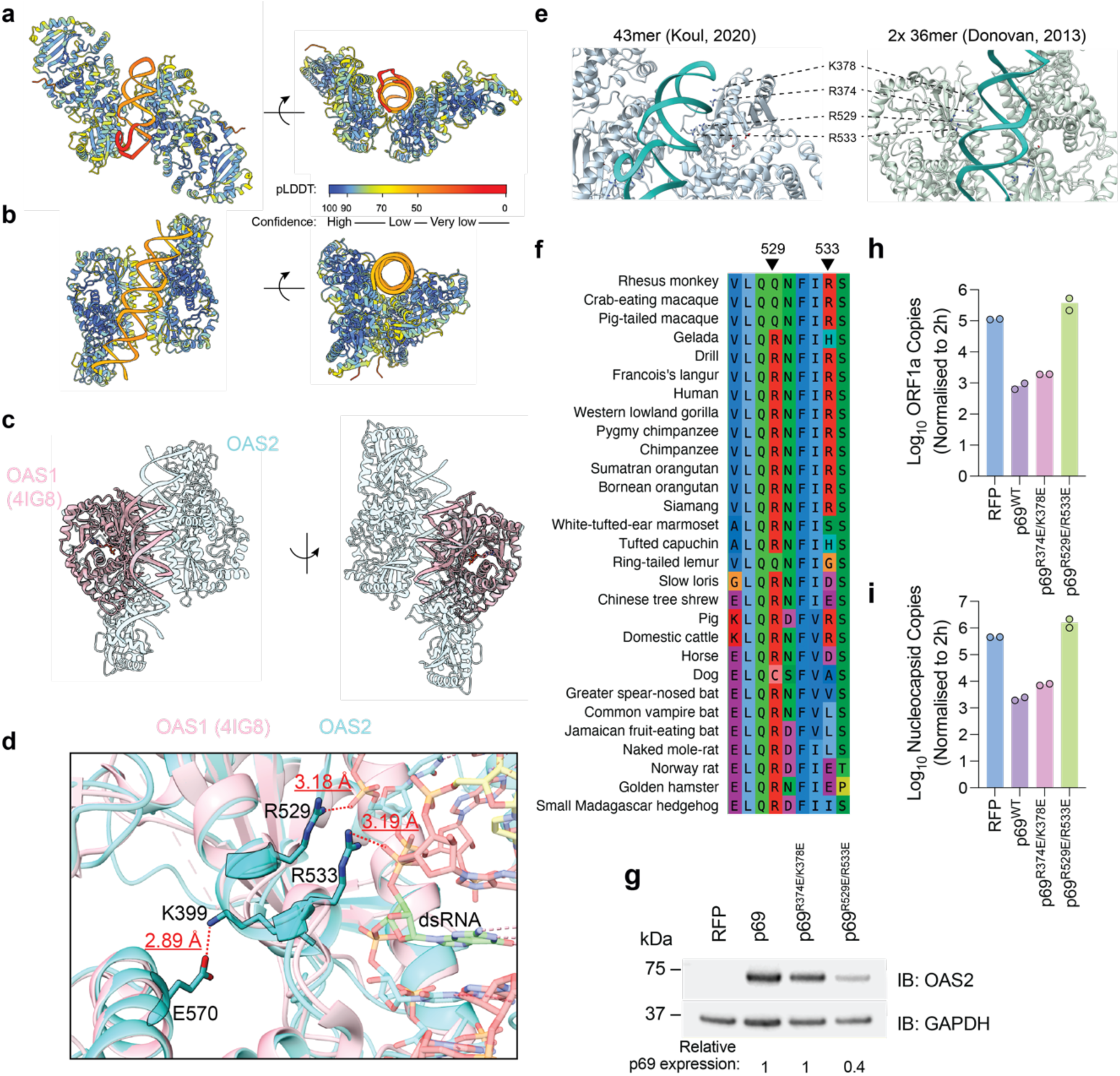
– Predicted RNA binding residues are necessary for OAS2 antiviral activity. **a-b**) Top-ranking AlphaFold3 predictions of human OAS2 p69 in complex with dsRNA 43mer palindromic RNA (Koul *et al*., 2020a) (**a**) or 2×36mer RNA (Donovan *et al*., 2013) (**b**), coloured according to pLDDT scores as indicated by key. **c)** Superposition of OAS1:18mer RNA (Donovan *et al*., 2013) (PDB: 4IG8) with OAS2 p69:2×36mer from (**a**). (**d**) Close up of E570:K399 salt bridge, analogous to that identified in OAS1:18mer (Donovan *et al*., 2013), and displaced R529 and adjacent R533, in proximity of phosphate backbone of RNA; predicted structure superimposed with OAS1:18mer crystal structure (PDB: 4IG8) (Donovan *et al*., 2013) **e)** Position of R374, K378, R529 and R533 to dsRNA in predicted OAS2:dsRNA models. **f)** OAS2 multiple sequence alignments across primate and mammalian orthologues, focusing on human R529 and R533. **g)** A549 cells were modified to express p69 with substitutions predicted to bind RNA, confirmed by Western blotting. **h,i)** HCoV-OC43 ORF1a **h)** and nucleocapsid **i)** levels were measured in cells characterised in **g)**, infected with HCoV-OC43 (MOI 0.01) for 72 h, by RT-qPCR; data shown are the mean of 3 biological replicates.

The OAS2-2×36mer AF3 structure superimposed well with the OAS1-dsRNA crystal structure (root mean square deviation (RMSD) value between 305 pruned atom pairs was 0.685 Å) (**Fig. 4c**), with several conserved side chains stabilising interaction with RNA in both structures. In particular, the OAS2-2×36mer model captured a key conformational change that occurs upon dsRNA binding in OAS1 (Donovan *et al*., 2013). In an apo structure of porcine OAS1 (Hartmann *et al*, 2003), residue R195 (human OAS1 numbering) forms a salt bridge with an internal glutamate (E233). Upon dsRNA binding, distant residue K66 moves 18 Å to exchange places with R195, displacing R195 by 11 Å and causing it to rotate and form hydrogen bonds with backbone phosphates (Donovan *et al*., 2013). In the predicted OAS2-2×36mer structure, the equivalent glutamate (E570) is seen to form a salt bridge with K66-equivalent K399, freeing R529 for interaction with RNA (**Fig. 4d**). Nearby OAS2 residue R533, also conserved in OAS1, similarly forms hydrogen bonds with RNA phosphates (**Fig. 4d**). Double mutation of these residues in porcine OAS1 (R194E/R198E) caused a ∼10,000 reduction in dsRNA-dependent 2-5A catalysis (Hartmann *et al*., 2003). In the AF3 structure, we noticed a nearby cluster of basic residues – R374 and K378 – also face the 2×36mer RNA; these superimpose with a cysteine (C38) or lysine (K42) in OAS1, respectively (**Fig. 4e**). Mutation of the latter in porcine OAS1 (K41E) caused a ∼ 62-fold reduction in dsRNA-dependent 2-5A catalysis (Hartmann *et al*., 2003). The proximity of these four basic side chains in OAS2 (R374, K378, R529 and R533) to RNA was conserved for both dsRNA ligands (**Fig. 4e**), and across all ten ranked structural predictions (**Supplementary Fig. 4f**), further suggesting their relevance in RNA binding. Of note, the displaced R529 and, to a lesser extent, the nearby R533, are conserved in bovine, equine, rodent, porcine and primate OAS2 sequences, although interestingly human R529 is a glutamine in the *Macaca* sequences analysed (**Fig. 4f**).

Guided by these analyses, we introduced a non-conservative R529E/R533E double substitution in OAS2 p69 to ablate interaction with viral RNA. We also generated a double R374E/K378E variant because R374 is adjacent to T375, the equivalent of porcine K38, which is also important for porcine OAS1 activity (Hartmann *et al*., 2003). p69^R374E/K378E^ and p69^R529E/R533E^ mutants were stably expressed in A549 cells; consistently, the expression of the p69^R529E/R533E^ mutant was 2.7-fold lower than p69^WT^ (**Fig. 4g**). We next infected these cells and used qPCR to monitor viral replication. As before, p69 caused a strong (140-fold) inhibition of viral RNA synthesis relative to the RFP control (**Fig. 4h**). This was reduced to a 60-fold inhibition in cells expressing p69^R374E/K378E^ cells, and completely lost in p69^R529E/R533E^ cells, compared to the RFP control (**Fig. 4h**), suggesting a total loss of viral restriction. Similarly, we measured a 200– and 60-fold reduction in N transcripts in p69^WT^ and p69^R374E/K378E^ cells respectively, and no restriction to N transcription in p69^R529E/R533E^ cells, compared to the RFP control (**Fig. 4i**). In summary, structural predictions and sequence analyses support the model that RNA-binding is necessary for OAS2 p69 restriction of HCoV-OC43.

### OAS2 p69 inhibits HCoV-OC43 independently of RNase L

As described above, the antiviral activity of OAS proteins occurs canonically via synthesis of 2-5A, which activates RNase L. Because OC43 encodes a phosphodiesterase that degrades 2-5A (Goldstein *et al*., 2017), we hypothesised that OAS2 p69 inhibits OC43 through a mechanism that does not involve 2-5A. Accordingly, we introduced a mutation to disrupt the catalytic triad of aspartic acids (D408, D410, D481), which collectively coordinate two catalytic metal ions in all enzymes of the nucleotidyl transferase superfamily, including OASes, and are required for 2-5A synthesis (Donovan *et al*., 2013; Koul *et al*., 2020b; Sarkar *et al*, 1999b). OAS2 D481 is also in proximity to the site of the donor ATP molecule (Sarkar *et al*, 2002), and the D481A variant in OAS2, or equivalent D148A in OAS1, are both catalytically inactive (Donovan *et al*., 2013; Koul *et al*., 2020b). In addition, we generated a variant substituted at the putative CFK dimerization residues (CAFAKA) (Ghosh *et al*, 1997). We expressed these variants in A549 cells; p69^CAFAKA^ expression was 2.4-fold lower than that of p69^WT^ whereas p69^D481A^ expression was equivalent to that of p69^WT^ (**Fig. 5a**). At 72 hpi, HCoV-OC43 ORF1a transcripts were 89-, 137– and 74-fold lower in the p69^WT^, p69^D481A^, p69^CAFAKA^ cells respectively, compared to the RFP control (**Fig. 5b**). Similarly, HCoV-OC43 N transcripts were 156-, 190-, 140-fold lower in the p69^WT^, p69^D481A^, p69^CAFAKA^ cells respectively, compared to the RFP control (**Fig. 5c**). The lack of effect of CAFAKA mutation is supported by our structural predictions, where the CFK residues did not appear at the OAS2 dimer interface (**Supplementary Fig. 4e**), as well as recent *in vitro* studies where CAFAKA mutation was insufficient to disrupt OAS2 oligomerisation and activity (Koul *et al*., 2020b). Combined, these data supported our initial hypothesis that the antiviral activity of p69 was not via the canonical 2-5A/RNase L pathway.

**Figure 5.**
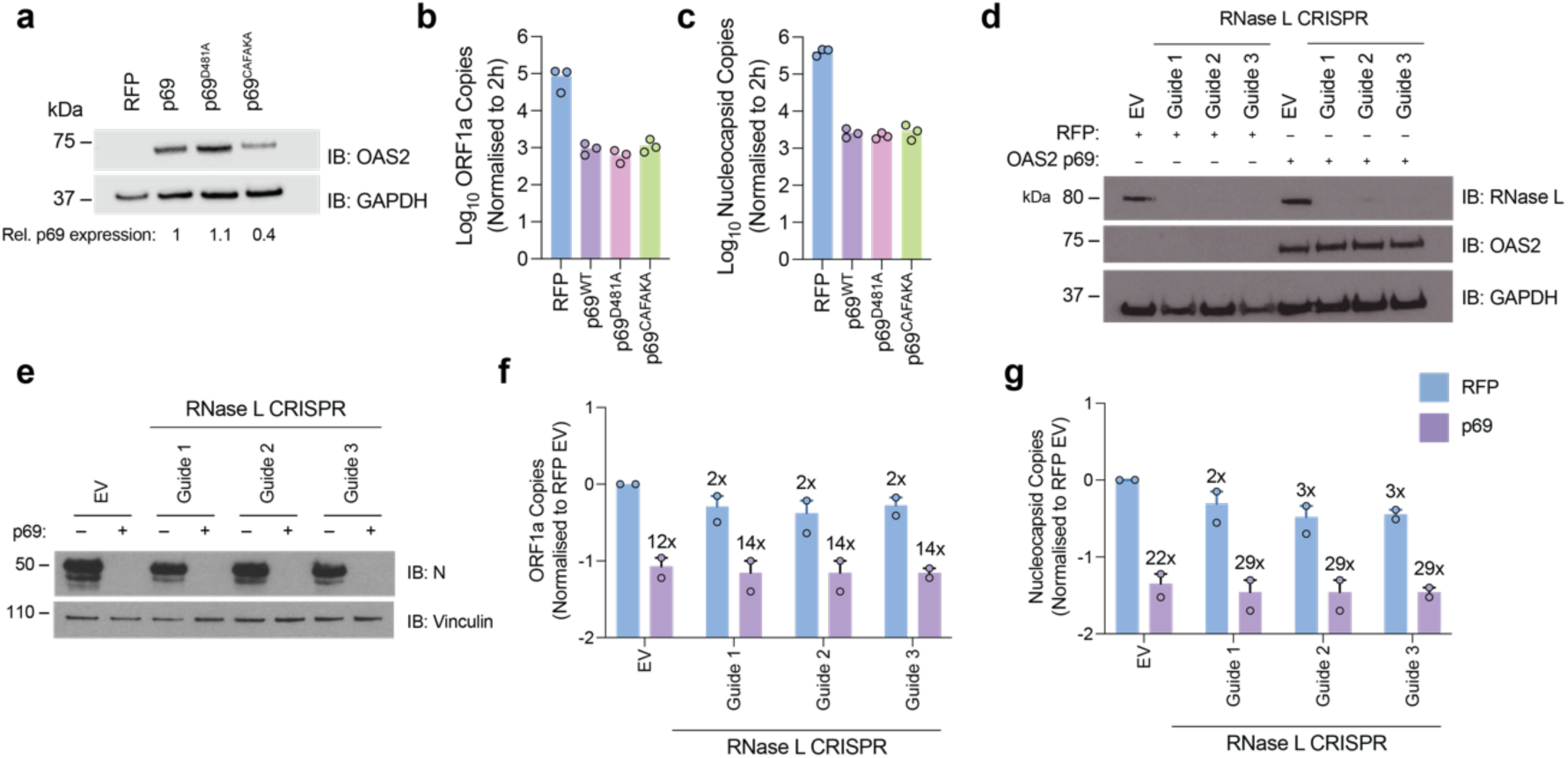
– OAS2 p69 restriction of HCoV-OC43 is RNase L-independent. **a)** A549 cells were modified to express a catalytic mutant of p69 (D481A) or a CFK motif mutant (CAFAKA), confirmed by Western blotting. **b,c)** Cells from (**a**) were infected with HCoV-OC43 for 72 h, and *ORF1a* (**b**) or *nucleocapsid* (**c**) transcripts were quantified by RT-qPCR; data shown are the mean of 3 biological replicates. **d)** A549 cells expressing RFP or OAS2 p69 were depleted of RNase L using three CRISPR guides, and levels of RNase L and OAS2 were confirmed by Western blotting. **e)** Cells from (**d**) infected with HCoV-OC43 for 72 h and levels of N protein determined by Western blotting. **f,g)** Cells from (**d**) were infected with HCoV-OC43 for 48 h, and *ORF1a* (**f**) or *nucleocapsid* (**g**) transcripts were quantified by RT-qPCR; data shown are the mean of two biological replicates. Empty vector (EV).

To test RNase L-independence directly, we stably depleted RNase L from A549 cells expressing RFP or OAS2 p69, using CRISPR/Cas9. Relative RNase L depletion was assessed by Western blot (**Fig. 5d**). The most effective guide RNAs were taken forward into HCoV-OC43 infection assays, and viral replication was monitored by Western blot. Remarkably, despite robust RNase L depletion, the ability of OAS2 p69 to inhibit viral replication was largely unaltered, compared to control cells expressing Cas9 without guide RNA (**Fig. 5e**). We also measured viral ORF1a and N transcripts in these cells at 48 hpi. Despite a modest reduction in p69 potency, these experiments also revealed robust RNase L-independent p69 restriction of viral transcription (**Fig. 5f,g**). Combined with the full activity of the catalytically dead p69^D481A^ variant, these data suggest that OAS2 restricts HCoV-OC43 in a manner independent of the canonical 2’-5’A/RNase L pathway.

### The p71 isoform of OAS2 restricts EMCV in an N-myristoylation and RNase-L dependent manner

Recent data suggests that certain OAS enzymes, like human OAS1, can exhibit both RNase L-dependent and independent restriction mechanisms, in a virus-specific manner (Harioudh *et al*, 2024). To ask whether OAS2 might display RNase L-dependent restriction in alternative scenarios, we tested its ability to restrict the picornavirus cardiovirus A (EMCV). EMCV, like HCoV-OC43, replicates in membranous organelles (Romero-Brey & Bartenschlager, 2014) and was shown to be sensitive to OAS2 (Marié *et al*., 1999).

We infected A549 cells expressing RFP, p69 or p71, with serial dilutions of EMCV. EMCV infectious titres were ∼2.5-fold reduced in cells expressing p71 compared to the RFP control but showed no difference in cells expressing p69 (**Fig. 6a**). Although the difference in titre was modest, the difference in plaque phenotype between the OAS2 isoforms was substantial, with p71 considerably restricting plaque area (**Fig. 6b**). As we found for p69-mediated restriction of HCoV-OC43, p71’s antiviral activity toward EMCV required N-terminal myristoylation, with 15-fold more plaques observed in the presence of the p71^G2A^ variant than myristoylated p71^WT^ (**Fig. 6c**). Moreover, the plaques were considerably larger in the presence of p71^G2A^ than in the presence of p71^WT^ cells (**Fig. 6b**). To test whether EMCV restriction by p71 was also RNase L-independent, we depleted RNase L from p71 expressing A549 cells (**Fig. 6d**), and measured EMCV replication by plaque assay. Unlike p69 and HCoV-OC43, EMCV infectious titre in cells expressing p71 was rescued 10-fold (**Fig. 6e**), with a corresponding enlargement of plaque area (**Fig. 6f**). While we also observed a ∼4-fold increase in EMCV titre following RNase L depletion from control RFP-expressing cells, which we speculate is due to endogenous OAS proteins (Li *et al*, 2016), in this case this occurred without concomitant increase in plaque area (**Fig. 6e,f**). These data therefore demonstrate that OAS2 p71 restricts EMCV via the canonical RNase L-dependent pathway (**Fig. 6g**).

**Figure 6.**
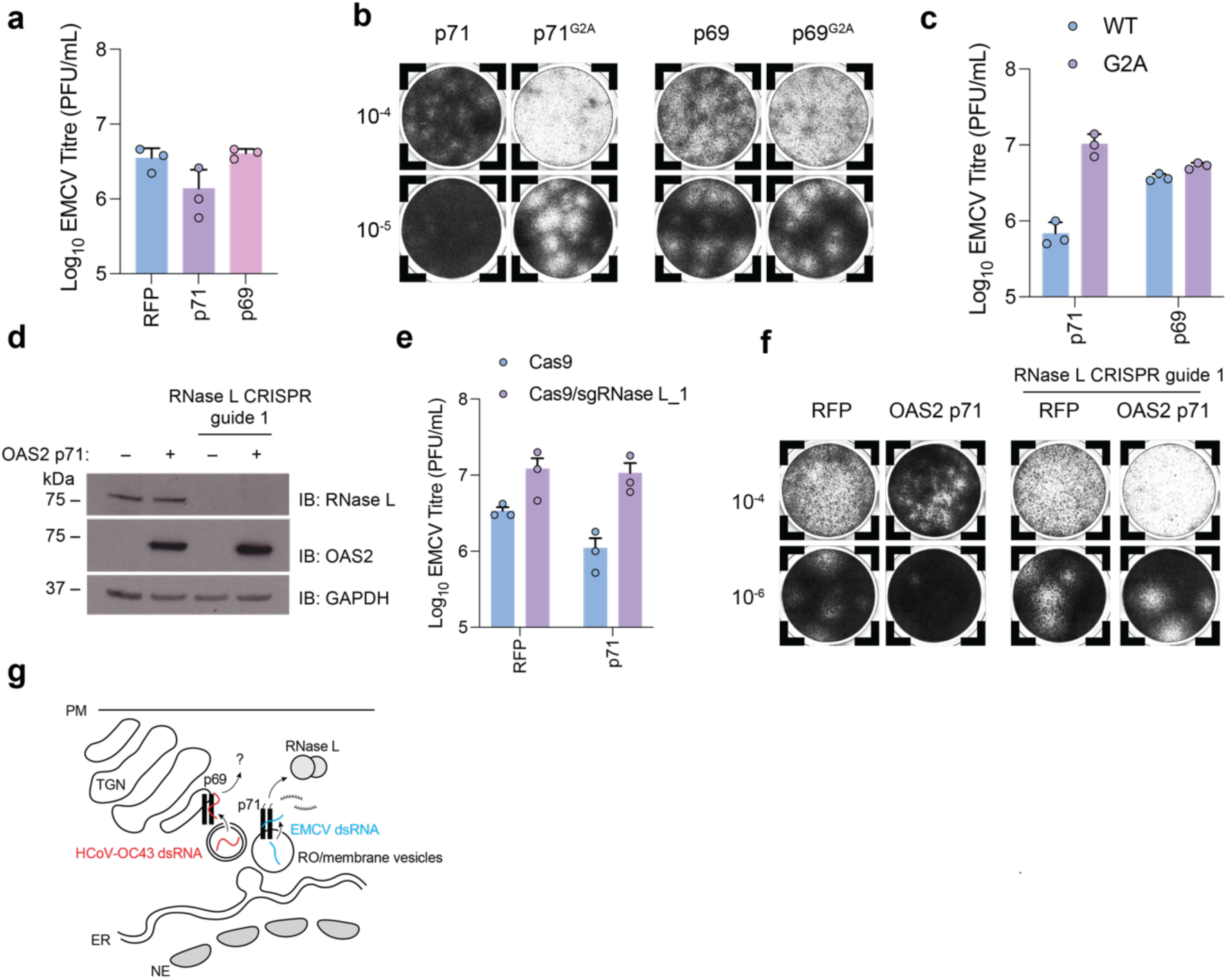
– EMCV is restricted by OAS2 p71 in an RNase L-dependent manner. **a)** Infectious titres of EMCV in cells expressing RFP, p71 or p69 were determined by plaque assay at 72 hpi. **b)** Representative images of EMCV plaques formed in cell lines expressing the OAS2 isoforms or their respective myristoylation (G2A) defective mutants. **c)** Infectious titres of EMCV in cell lines expressing OAS2 isoforms or their respective G2A mutants, determined by plaque assay at 72 hpi. **d)** A549 cells expressing RFP or p71 were depleted of RNase L using three CRISPR guides, and expression levels of RNase L and OAS2 were confirmed by Western blotting. **e)** Infectious titre of EMCV in the cells characterised in (**d**), determined by plaque assay at 72 hpi. **f)** Representative images of EMCV plaques formed in the cell lines characterised in (**d**). **g)** Model for sensing of viral dsRNA by OAS2 p69 and p71 isoforms. Coronaviruses or picornaviruses replicate in or in proximity to single-or double-membraned vesicles (also known as replication organelles, RO) derived from the endoplasmic reticulum (ER), itself continuous with the nuclear envelope (NE). OAS2 isoforms localise to membranes proximal to the trans-Golgi network (TGN). p69 senses dsRNA from HCoV-OC43 and inhibits viral replication by an unknown mechanism; p71 senses dsRNA from EMCV and inhibits viral replication via RNase L. PM, plasma membrane. (**a,c,e**) Data shown are the mean of three biological replicates.

### OAS2 C-terminal tails shape antiviral specificity

The distinct antiviral profiles of OAS2 isoforms p69 and p71 suggest their short C-terminal tails harbour important determinants of restriction. To explore this, we first compared AF3 structural predictions of either isoform in complex with the 2×36mer RNA. pLDDT scores for the p71 tails were low (**Fig. 7a,b, Supplementary Fig. 5b,c**), suggesting they lack a defined binding surface on the OAS2 core scaffold and likely experience significant conformational dynamics. The proximity of these tails to RNA suggests they might even clamp around nucleic acid and provide sequence specificity (**Supplementary Fig. 5b,c**). Although significantly shorter, the p69 4-residue tail is also not confidently modelled (**Fig. 7a,c,d**). Lack of defined structure begins at residue K670 (**Fig. 7d**), and while AF3 is more confident in the position of residues 670-683, this surface-exposed peptide may also reposition in the presence of physiological cofactors. Thus, structural predictions – which accurately predict OAS2:dsRNA interfaces – suggest that the OAS2 C-terminal tails sit in proximity to nucleic acid, potentially explaining how they influence antiviral specificity.

**Figure 7.**
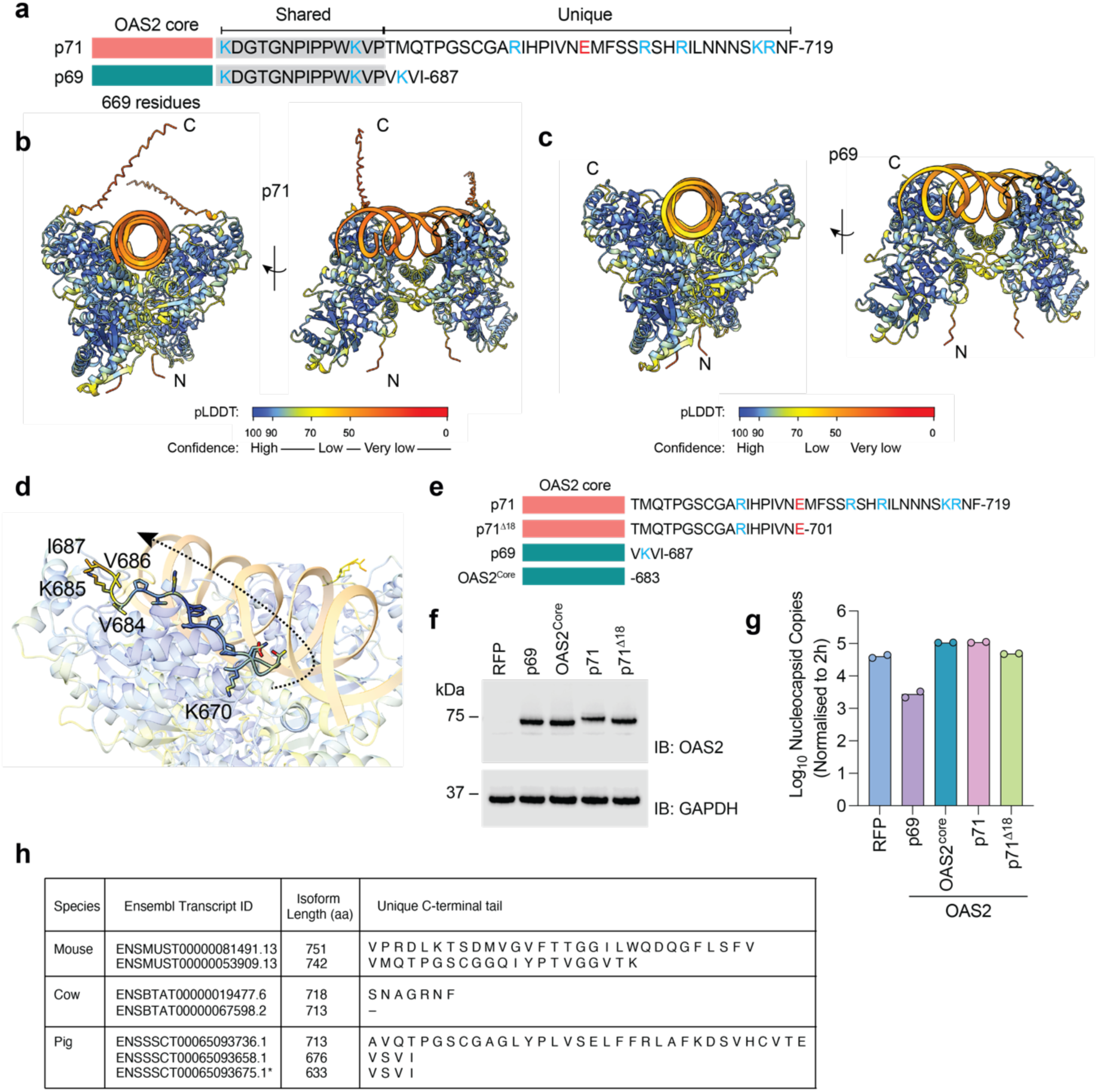
– OAS2 C-terminal tails shape antiviral specificity. **a)** Schematic of the OAS2 isoforms with shared and distinct C-termini sequences indicated. Basic residues are coloured blue, acidic residues are coloured red. **b-c)** AlphaFold3 top-ranked models from Figure 4, of p71 (**b**) or p69 (**c**) with dsRNA (Donovan *et al*., 2013) coloured by pLDDT score, indicated in colour key. OAS2 N– and C-termini are labelled N and C, respectively. **d)** Closeup of OAS2 p69 from structure presented in (**c**), showing side chains of peptide beginning at K670, until terminal residue I687, coloured as in (**c**). The dashed arrow indicates the sequence direction from 670-687. pLDDT scores are high between residues 670-683, and low between residues 684-687. **e)** Schematic of the OAS2 variants generated. Basic residues are coloured blue, acidic residues are coloured red. **f)** A549 cells expressing the OAS2 variants indicated in (**e**) were assessed by Western blot. **g)** Cells characterised in (**f**) were infected with HCoV-OC43 for 48 h (MOI 0.01), and *N* copies quantified by qPCR. Data shown are the mean of two biological replicates. A one-way ANOVA statistical test was used to test significance, threshold p = 0.05. **h)** Ensembl transcripts indicating encoded protein length and within-species sequence divergence at C-termini. *ENSSSCT00065093675.1 lacks an N-terminal peptide encoding Gly2 and so is predicted not to undergo N-terminal myristoylation.

To test this hypothesis, we generated OAS2 variants with truncated tails. For p71, we removed half of the tail (deletion M702-F719, p71^Δ18^), and we generated a second variant where we removed the whole tail region, representing a protein that is present in both p69 and p71 isoforms (residues 1-683, OAS2^core^) (**Fig. 7e**). We expressed these variants in A549 cells (**Fig. 7f**) and infected cells with HCoV-OC43, measuring N transcripts at 48 hpi. Compared to control RFP expressing cells, we observed a 15-fold decrease in genomic+subgenomic (N) transcripts in p69^WT^-expressing cells (**Fig. 7g**). However, in p71^WT^, p71^Δ18^ or OAS2^core^-expressing cells, N transcripts were comparable to those in control cells, suggesting that restriction of HCoV-OC43 is not inhibited by the long p71 tail (**Fig. 7g**). Importantly, these data demonstrate that the short 4-residue tail of p69 is necessary for p69-mediated inhibition of HCoV-OC43.

*OAS2* orthologues also exhibit isoform-specific C-terminal diversity. Across *OAS2* transcripts in pigs, cows and mice, a similar pattern of C-terminal length variation is observed (**Fig. 7h**), suggesting this mechanism of broadening antiviral diversity through alternative splicing might be conserved in mammals.

## Discussion

In this paper, we identify a collection of genes with antiviral activity against seasonal coronavirus HCoV-OC43. Focusing on *OAS2*, we find that the p69 isoform is antiviral towards HCoV-OC43, while the longer p71 isoform is not. Interestingly, the antiviral activity of p69 toward HCoV-OC43 occurred independently of the canonical 2-5A/RNase L axis, aligning OAS2 with other OAS systems recently reported to act independently of RNase L (Harioudh *et al*., 2024; Zhu *et al*, 2014) (**Fig. 6g**). In the context of HCoV-OC43, this alternative antiviral mechanism might provide immune defence in the face of a viral PDE, which is an effective OAS antagonist in related betacoronaviruses like mouse hepatitis virus (Goldstein *et al*., 2017) and was presumed to explain the lack of restriction by paralogue *OAS1* toward HCoV-OC43 (Wickenhagen *et al*., 2021). We found that the p71 isoform is antiviral toward the unrelated positive-sense cardiovirus EMCV, and, as its activity was RNase L-dependent, suggests that p71 acts as a canonical OAS2 sensor in this role. Our data therefore suggest that a single *OAS2* locus provides antiviral diversity through alternative splicing, and that mechanistic plasticity underlies this phenomenon. The concept of ‘sensor’/’restriction factor’ duality first emerged with the discovery that TRIM5 and tetherin – already well-characterised restriction factors – also stimulated NF-κB signaling pathways upon virus detection (Galao *et al*, 2012; Pertel *et al*, 2011). Since then, several other proteins have been shown to exhibit both activities (McEwan *et al*, 2013), even extending to the prototypical sensors like RIG-I (Sato *et al*, 2015). In this context, OAS2 represents an alternative solution to achieving restriction/sensor duality, whereby the distinct functions are not coordinated but rather split across two polypeptides derived from the same gene. Both isoforms are *bona fide* 2-5A synthetases *in vitro* (Marié *et al*., 1999), suggesting RNase L independence is a functional adaptation rather than a catalytic deficit. However, understanding whether restriction *versus* sensing are truly split, i.e. can p69 restrict a virus via 2-5A synthesis (and p71 via RNase L-independent mechanisms) will provide a clearer picture of how malleable the *OAS2* locus is in humans. Testing the replication of an NS2-deleted HCoV-OC43 (Diefenbacher *et al*, 2024) in p69– and p71-expressing cells could address whether any residual RNase L-dependent mechanism is indeed antagonised by this viral PDE.

Importantly, the C-terminal tails harbour critical determinants for virus specificity. Deletion of the short 4-residue tail of p69 was sufficient to abolish its antiviral activity toward HCoV-OC43. Both p69 and p71 C-terminal tails are characterised by periodic spacing of basic residues (and a paucity of acidic residues) (**Fig. 7a**), and we speculate that these tails could influence target RNA selection and binding. Interestingly, an OAS2 single nucleotide polymorphism (SNP) rs15895, which extends the OAS2 p71 tail by 8 residues, is associated with increased susceptibility to DENV infection and severe TBEV infection *in vivo* (Alagarasu *et al*, 2013; Barkhash *et al*, 2010). The ‘A’ allele encodes a stop codon after residue 719, whereas the ‘G’ allele encodes a tryptophan at position 720, followed by 7 additional residues. The original identification of the p71 isoform isolated cDNA from Daudi cells, which encoded the 727 amino acid protein rather than the 719 residue p71 cloned in our study (Marié & Hovanessian, 1992) (**Supplementary Fig. 5d,e**). Daudi cells originate from a Burkitt’s lymphoma patient of African origin, where the rs15895 ‘A’ allele occurs at a lower frequency than European populations (Gokul *et al*, 2023) (**Supplementary Fig. 5f**). Owing to European sampling bias, the ‘A’ allele encoding the 719 residue p71 is thus annotated as the canonical p71 sequence by NCBI and Ensembl and for this reason was the isoform cloned and characterised in our study. Thus, altering the OAS2 tail correlates with virus susceptibility *in vitro* and *in vivo*.

The evolution of RNase L-independent mechanisms for OAS enzymes is perhaps unsurprising considering the diversity of viral PDE enzymes, which have been recurrently acquired by both nidoviruses and rotaviruses via horizonal gene transfer from host to virus (Goldstein *et al*, 2024). Moreover, *OAS* genes are evolutionarily ancient antiviral loci, appearing across eukaryotic taxa – more deeply so than the related cGAS-like receptors (cGLRs) (Culbertson & Levin, 2023; Li *et al*, 2023). Being able to restrict viral replication by diverse enzymatic mechanisms might afford a more robust defence in the face of rapidly evolving RNA viruses, particularly where OAS-RNA binding is sequence or structure specific, as indicated by OAS1 p46 with SARS-CoV-2 (Wickenhagen *et al*., 2021), West Nile Virus (WNV) (Koul *et al*., 2020a), or adenovirus-associated virus RNA (Desai *et al*, 1995). As for the precise antiviral mechanism behind p69 restriction of HCoV-OC43, clues might be found in related OAS systems like human OAS1, which in addition to generating 2-5A, can bind and stabilise IFNβ mRNA, potentiating IFN signalling strength during WNV replication (Harioudh *et al*., 2024). Unlike its single human orthologue, murine Oas1b is devoid of 2-5A synthetase activity (Elbahesh *et al*, 2011), but remains potently antiviral towards WNV (Perelygin *et al*, 2002). Many mouse lab strains are highly susceptible to WNV because they encode a truncated Oas1b protein lacking the C-terminal transmembrane domain that localises Oas1b to the endoplasmic reticulum (Courtney *et al*, 2012; Perelygin *et al*., 2002). While the detail of the RNase L-independent Oas1b restriction mechanism remains unclear, it is thought to involve interaction with the ATP binding cassette protein 3, subfamily F (ABCF3) (Courtney *et al*., 2012). Human OASL is a catalytically inactive paralogue but via its C-terminal ubiquitin-like domains, OASL binds to RIG-I by mimicking di-ubiquitin, enhancing RIG-I signalling (Zhu *et al*., 2014). Human OAS2 is also seen to localise to stress granules (Reineke & Lloyd, 2015), which can coordinate antiviral defence against diverse RNA viruses. Ongoing work is exploring whether OAS2 p69 restriction operates via any of these pathways.

Irrespective of catalytic mechanism, we find that the restriction activity of both OAS2 isoforms is wholly dependent on membrane association, aligning it with the related OAS1s, including human OAS1 p46 (Soveg *et al*., 2021; Wickenhagen *et al*., 2021), horseshoe bat OAS1 (Lytras *et al*, 2023; Wickenhagen *et al*., 2021), and murine Oas1b (Courtney *et al*., 2012). Across these three examples, the method of membrane attachment is distinct, exemplifying the plasticity of the OAS system in achieving a similar goal. Whether N-terminal myristoylation, versus C-terminal prenylation, is particularly important for OAS2 restriction remains unknown – the addition of a CAAX box to an OAS2 G2A myristoylation-deficient variant could reveal whether lipid association by any means is sufficient to support antiviral activity. Alternatively, myristoylation might direct OAS2 to highly specific membrane subdomains that reflect its viral target. Alternatively, the N-terminal lipidation could be important for a protein where RNA binding specificity is dictated by the C-terminal tail, as we hypothesised above. Across OAS systems, the topology of dsRNA sensing remains unclear, given that coronaviruses and cardioviruses likely replicate within single or double-membrane vesicles (DMV); in such cases, vRNA might transits through exit pores to become exposed to OAS enzymes anchored to the cytosolic face of the vesicle (**Fig. 6g**). Alternatively, anchored OASes could infiltrate the DMV during its biogenesis from the endomembrane system.

Antiviral breadth is hardwired into the IFN response. The parallel expression of multiple ISGs provides the cell with a broad palette of antiviral activities against many viral families (McDougal *et al*., 2023; Schoggins *et al*, 2011), even where some ISG products, like TRIM5 (Ganser-Pornillos & Pornillos, 2019), are highly target-specific. Some ISGs independently achieve antiviral breadth by virtue of generic restriction mechanisms; BST-2/tetherin prevents the release of enveloped viral particles belonging to diverse virus families including *Retroviridae*, *Flaviviridae*, *Filoviridae*, *Orthoherpesviridae*, *Peribunyaviridae* and *Coronaviridae* (Hagelauer *et al*, 2023; Neil, 2013; Varela *et al*, 2017), while arguably, OAS enzymes achieve this through the indiscriminate endoribonuclease RNase L. Alternatively, gene duplication provides novel genetic material with which to mould the antiviral portfolio, exemplified by the *OAS1* and *TRIM5* loci in rodents (Elkhateeb *et al*, 2016; Tareen *et al*, 2009)., although whether gene duplication broadens virus specificity remains speculative. Here, we reveal a novel mechanism, where through alternative splicing and mechanistic plasticity, a single *OAS* gene expands the antiviral portfolio. We speculate that this might be an evolutionarily economical method for achieving antiviral breadth, compared to whole gene duplication. As the majority of multi-exon genes undergo alternative splicing (Kjer-Hansen & Weatheritt, 2023), this is likely to be a more common strategy for antiviral defence than currently appreciated.

## Acknowledgements

We thank all members of the Fletcher group for remarks on the manuscript and discussions. We thank Colin Loney for assistance in image cytometry, Joseph Hughes for discussions around OAS2 transcript diversity and Suzannah Rihn for discussion around ISG libraries. This work was supported by funding from a UKRI Future Leaders Fellowship (MR/T043482/1, H.S., A.B., R.H., E.M., A.J.F.), a Wellcome Trust PhD Fellowship (218518/Z/19/Z, E.L.D.), and addition funding from the Medical Research Council (MC_UU_00034/3, M.P.; MR/P022642/1 and MR/V01157X/1, S.J.W., M.T., A.W.).

## Author Contributions

E.L.D., S.J.W. and A.J.F. designed experiments, coordinated the project and wrote the manuscript. E.L.D. performed all infection experiments, including image cytometry, performed molecular cloning, and analysed data, with help from H.S., E.M., A.W. and M.T.. E.L.D. and A.B. performed confocal microscopy experiments. A.J.F. performed AF3 structural prediction analyses and prepared figures. A.B. performed microscopy and associated data analysis. E.L.D. and A.J.F. analysed data. S.J.W. designed the ISG screens, and E.L.D. and S.J.W. grew and prepared material for ISG screens.

## Materials and Methods

### Cells and viruses

HEK-293T, A549, VeroE6 and Huh7.5 cells were propagated in Dulbecco’s Modified Eagle Medium (DMEM; ThermoFisher) supplemented with 10% fetal bovine serum (FBS; ThermoFisher) and 10 μg/mL gentamicin (Melford Laboratories) or 100 U/mL penicillin/100 μg/mL streptomycin (ThermoFisher). Betacoronavirus HCoV-OC43 (VR-1558) was purchased from ATCC and propagated on VeroE6 cells (**Figure 1**) or Huh7.5 cells (all other figures) at 33 °C. Encephalomyocarditis virus (VR-129B) was purchased from ATCC and propagated on VeroE6 cells at 37 °C.

### Retroviral vectors and plasmids

The screening lentiviral vector pSCRPSY (GenBank accession no. KT368137.1) and the modified plasmid pLV-EF1a-IRES-*SfiI*-Puro-TagRFP have been described previously (Wickenhagen *et al*., 2021). cDNA for OAS2 p71 (NM_016817.3) and OAS2 p69 (NM_002535.3) were ordered as gene blocks (IDT DNA) with flanking *SfiI* sites and subcloned into the pLV plasmid. The OAS2-p71^G2A^ and OAS2-p69^G2A^ mutants were generated by PCR amplifying the relevant gene blocks with primers that mutate Gly2 to Ala. OAS2-p71^Δ18^ and OAS2^core^ constructs were generated by PCR amplification from OAS2-p71 templates using forward primer 5’-GCC TCA GAC AGT GGT TCA AAG-3’ and reverse primers 5’-CCG CCC TCG AGG AAT TCG GCC AGA GAG GCC CTA CGG CAC TTT CCA AGG TGG TAT TGG GTT TC-3’ for OAS2^core^ and 5’-CCG CCC TCG AGG AAT TCG GCC AGA GAG GCC CTA CTC ATT GAC AAT AGG ATG GAT CCT AGC-3’ for p71^Δ18^, then introduced into pLV-EF1a-IRES-SfiI using Gibson Assembly (NEB) following the manufacturer’s protocols.

The OAS2-p69^D481A^, OAS2-p69^CAFAKA^, OAS2-p69^R374E/K378E^ and OAS2-p69^R529E/R533E^ mutants were generated by *in vivo* assembly (IVA) using the following overlapping primers. D481A (Fwd: 5’-CCT CAA CGA AAG TGT CAG CTT TGC AGT GCT TCC TGC CTT TAA TGC ACT GGG-3’, Rev: 5’-AAA GCT GAC ACT TTC GTT GAG GAC TTT GG-3’), CAFAKA (Fwd: 5’-GGA ATG GTT ATC CTC TCC CGC CGC CGC GGA TGG GAC TGG AAA CCC AAT ACC ACC-3’, Rev: 5’-GGG AGA GGA TAA CCA TTC CTT TGC TTC-3’), R374E/K378E (Fwd: 5’-ATT GAC AGT GCT GTT AAC ATC ATC GAA ACA TTC CTT GAA GAA AAC TGC TTC CGA CAA TCA ACA G-3’, Rev: 5’-GAT GAT GTT AAC AGC ACT GTC AAT CTG CTC-3’), R529E/R533E(Fwd: 5’-CTG TTT CAC AGT CCT GCA GGA AAA CTT CAT TGA ATC CCG GCC CAC CAA ACT AAA GG-3’, Rev: 5’-CTG CAG GAC TGT GAA ACA GGT AGA AAA C-3’). IVA reactions were performed using 1 ng template, 100 nM primers, 200 μM dNTPs, 3% DMSO, 0.5 μL Herculase II Fusion DNA polymerase (Agilent). Cycle parameters were 95 °C 30 s, 18 cycles of 95 °C 10 s, 60 °C 30 s, 72 °C 3 min. A final hold of 72 °C for 10 min was performed. 1 μL FastDigest DpnI (Thermo Fisher) was added and incubated at 37°c for 15 min. 1 μL reaction was added to 50 μL NEB® 5-alpha cells (NEB C2987H), incubated on ice 15 min, heat shocked at 42 °C 30 s, incubated on ice for 2 min, then 200 μL SOC media (NEB) added and cells recovered at 37 °C, 200 rpm, for 45 min. The complete mixture was plated on LB agar containing 200 μg/mL ampicillin. Colonies were screened by analytical restriction enzyme digest with HindIII (Thermo Fisher) followed by Sanger sequencing.

LentiCRISPRv2-Puro and lentiCRISPRv2-Blast plasmids were used to knockdown expression of OAS2 and RNase L, respectively (Sanjana *et al*, 2014). CRISPR guides were designed using the CHOPCHOP tool (https://chopchop.cbu.uib.no) (Labun *et al*, 2019). Seven guides were subcloned into the lentiCRISPRv2 system and the three best guides that depleted target protein expression were used for subsequent experiments. The following guides were used: OAS2 (guide 1: 5-GTC TTA AGA GGC AAC TCC GA-3’, guide 2: 5’-GGA CGG AAA ACA GTC TTA AG-3’, guide 3: 5’-GCT TAC TCA GAG CGT TGA AGG-3’) and RNase L (guide 1: 5’-GCC GAG TTG CTG TGC AAA CG-3’, guide 2: 5’-TTA TCC TCG CAG CGA TTG CG-3’, guide 3: 5’-TAT AGG ACG CTT CGG AAT GT-3’) (Labun *et al*., 2019).

Retroviral vectors used for cell modification were produced by transient transfection in HEK-293T using the vector plasmid, HIV-1 gag-pol (pNLGP) and vesicular stomatitis virus glycoprotein expression plasmid (pVSVg), as described previously (Rihn *et al*, 2019). A549 cells were transduced with vector-containing supernatants filtered using a 0.45 μm-pore-size filter. Transduced cells were selected with 2 μg/mL puromycin (Melford Laboratories) or 5 μg/mL blasticidin (Melford Laboratories).

### Arrayed ISG expression screening

ISG lentiviral libraries encoding human, macaque and bovine genes have been described previously (Hardy *et al*, 2023; Wickenhagen *et al*., 2021). Briefly, libraries were generated in HEK-293T seeded in 96-well plates at a density of 3.5×10^5^ cells/well. HEK-293T were transfected with pSCRPSY, pNLGP and pVSVg at a ratio of 125 ng:25 ng:5 ng in the presence of Polyethylenimine (PEI, Polysciences). Supernatant was collected 48, 72 and 96 h post-transfection and replaced with fresh medium.

To perform the multi-species library screen, A549 cells were seeded in 96-well Pheno plates (PerkinElmer) at a density of 5×10^3^ cells/well and grown overnight. Cells were transduced with the 50 μL ISG lentiviral libraries (one ISG per well) and spinoculated at 500 xg for 1 h. 48 h post-transduction, cells were infected with HCoV-OC43 diluted in DMEM supplemented with 2% FBS. After incubation at 33 °C for 72 h, cells were fixed with a final concentration of 2% formaldehyde for 30 minutes, washed with PBS, and then permeabilised with 0.2% Triton X-100 (Sigma-Aldrich) for 5 min. After cells were washed and blocked with 1% Bovine Serum Albumin (Sigma-Aldrich) for 1 h, cells were incubated with 1 μg/mL of mouse anti-dsRNA IgG2a (10010500, Nordic-MuBio) overnight at 4 °C. After repeating washing and blocking steps, cells were incubated with 2 μg/mL goat anti-mouse IgG2a AlexaFluor488 (A-21131, Invitrogen) and 5 μg/mL Hoechst 33342 (Invitrogen) for 1 h. Screening plates were scanned and analysed using the Celigo Imaging Cytometer (Nexcelom Biosciences). Transduced and infected cells were gated using FlowJo v10.8. Data was normalised to the mean of each species library, and z-scores for each gene were calculated. ISGs with <10% transduction efficiency were excluded from hit selection.

Miniscreen libraries were generated as before. To measure cytotoxicity and ISRE induction, A549-ISRE::GFP cells were transduced with the 50 μL miniscreen library supernatant and spinoculated for 1 h at 500 xg. Supernatant was collected 120 h post-transduction and tested with the CytoTox-Glo™ Cytotoxicity assay (Promega), following manufacturer’s protocols. Cells were disassociated using 0.5% trypsin-EDTA (ThermoFisher) before fixation with formaldehyde. GFP+ cells were measured using the Guava EasyCyte flow cytometer (MilliPore); 10000 events/well were acquired and data were analysed using FlowJo v10.8. Data was normalised to SCRPSY-EMPTY controls.

### Virus infections

For HCoV-OC43 plaque assays, modified A549 cells were seeded at 4×10^5^ cells/well in a 12-well plate and grown to confluency overnight. Cells were inoculated with 250 μL HCoV-OC43 serially diluted 10-fold in serum-free DMEM and incubated at 33 °C for 1 h. An overlay of 1:1 Avicel (1.2%; FMC) and 2× MEM (Gibco), supplemented with L-Glutamine, Sodium Bicarbonate 7.5% (ThermoFisher), gentamicin and 4% FBS was then added. Cells were incubated for 5 days, fixed with 4% Formaldehyde solution, washed twice with PBS (Gibco), and stained with Coomassie Blue for plaque visualisation. The same conditions were used for EMCV plaque assays but were incubated for 3 days at 37 °C.

For infections for RT-qPCR and western blot analysis, MOIs were calculated using virus titre (PFU/mL) calculated by plaque assay on A549 cells. Unless otherwise stated, A549 derivative cells were infected at MOI 0.01 and incubated for 72 h prior to cell collection. Human interferon beta 1b (IFNβ; #11415-1) and alpha H2 (IFNɑ14; #11145-1) were obtained from Stratech Scientific.

### Western Blot Analyses

Cells were lysed on ice in a 1% NP-40 buffer containing protease and phosphatase inhibitors. After centrifugation, NuPAGE™ LDS Sample buffer containing 2-Mercaptoethanol was added. Proteins were separated on NuPage 4-12% BisTris polyacrylamide gels and transferred onto nitrocellulose membranes using the iBlot 2 (Invitrogen). After blocking in 5% milk (Sigma-Aldrich), membranes were probed against GAPDH (AM4300, Invitrogen), vinculin (13901S, Cell Signalling Technologies), OAS2 (19279-1-AP, Proteintech), RNase L (27281, Cell Signalling Technologies) or HCoV-OC43 nucleocapsid (DA116, MRC Protein Phosphorylation and Ubiquitylation Unit). Membranes were then stained with species IgG-specific secondary antibodies conjugated to horseradish peroxidase: donkey anti-sheep (A16041, Invitrogen), goat anti-rabbit (7074, Cell Signalling Technologies) and horse anti-mouse (7076, Cell Signalling Technologies). After addition of Pierce ECL substrate (Thermo Scientific), membranes were scanned using the LICOR Odyssey XF scanner or on the SRX-101A Medical Film Processor (Konica Minolta).

### RT-qPCR

Infected cells were lysed with TRIzol (Invitrogen), and the aqueous RNA layer was collected following chloroform precipitation. RNA was purified using the RNeasy mini kit (Qiagen), with on-column DNase treatment (Qiagen), following manufacturer’s instructions. cDNA was synthesised using SuperScript IV with random hexamer primers (Invitrogen). Host and viral gene expression was measured using TaqMan Fast Universal Master Mix (Applied BioSystems) and specific TaqMan probes listed below on the QuantStudio 3 Real-Time PCR machine (Thermo Fisher Scientific). Using the 2^-△△Ct^ method, viral transcript levels were normalised to ACTB and then normalised to input viral transcripts at 2 h in the respective cell line or 48 h/72 h in the control cell line. The following primers and probes were used:

ACTB(Hs01060665_g1, Thermo Fisher Scientific), HCoV-OC43_N_Fwd: 5’-CTA CTT CGC GCA CAT CCA G-3’, HCoV-OC43_N_Rev: 5’-GTC AGG TGT TAC ACC AGA GG-3’, HCoV-OC43_N_Probe: 5’-AGC CTC TAG TGC AGG ATC GCG TAG-3’, HCoV-OC43_ORF1a_Fwd: 5’-ATG TGG TGT AAA GCA GGA AC-3’, HCoV-OC43_ORF1a_Rev: 5’-GCA AGA ACA GTC CAC GGT ATA-3’, HCoV-OC43_ORF1a_Probe: 5’-TAC TGG TCT GGA CGC TGT TAT GC-3’.

### Immunofluorescence

A549 or modified cells were seeded at a density of 5×10^4^ cells/well onto glass coverslips. Cells were pre-treated with 1000 U/mL IFNβ or infected with HCoV-OC43 at MOI 10 for 24 h. After washing with PBS and fixing with 4% formaldehyde for 15 min at room temperature, cells were permeabilised in buffer containing 0.2% Bovine serum albumin and 0.2% Triton X-100 in PBS for 20 minutes at room temperature. The following antibodies, diluted in permeabilisation buffer, were used: rabbit polyclonal anti-OAS2 (19279-1-AP), mouse monoclonal anti-dsRNA (10010500, NordicMuBio), mouse monoclonal anti-58K (MA3-027, ThermoFisher), mouse monoclonal anti-calnexin (A160041, ThermoFisher), goat anti-mouse IgG2a Alexa Fluor™ 488 (A21131, ThermoFisher), Goat anti-rabbit IgG (H+L), Alexa Fluor™ 568 (A32731, ThermoFisher), goat anti-rabbit IgG (H+L) Alexa Fluor™ 568 (A11011, ThermoFisher), goat anti-mouse IgG (H+L) AlexaFluor™ 488 (A11001, ThermoFisher). Coverslips were incubated for 1 h with primary antibody followed by secondary antibody and Hoechst for 1 h. Maximum intensity projection images of cell monolayers were acquired with an Airyscan Fast detector fitted to a Zeiss LSM880 confocal microscope. Acquired images were analysed using ZEN software.

### Software

OAS2-dsRNA complex structural predictions were generated using the AlphaFold3 public server (www.alphafoldserver.com) (Abramson *et al*., 2024). Structural analyses and molecular graphics were performed with UCSF ChimeraX (version 1.8) (Meng *et al*, 2023). Protein sequences were analysed using Clustal Omega (Madeira *et al*, 2024) and AliView (Larsson, 2014). OAS2 sequence orthologues were obtained from the NCBI orthologues server. Graphs were generated using GraphPad Prism v10 (www.graphpad.com). Figures were prepared in Adobe Illustrator 2025. The schematic illustration in Supplementary Figure 1 was generated using BioRender software (https://app.biorender.com/).

## References

1. Abramson J, Adler J, Dunger J, Evans R, Green T, Pritzel A, Ronneberger O, Willmore L, Ballard AJ, Bambrick J et al (2024) Accurate structure prediction of biomolecular interactions with AlphaFold 3. Nature 630: 493–500

2. Alagarasu K, Honap T, Damle IM, Mulay AP, Shah PS, Cecilia D (2013) Polymorphisms in the oligoadenylate synthetase gene cluster and its association with clinical outcomes of dengue virus infection. Infection, Genetics and Evolution 14: 390–395

3. Barkhash AV, Perelygin AA, Babenko VN, Myasnikova NG, Pilipenko PI, Romaschenko AG, Voevoda AG, Brinton MA (2010) Variability in the 2′–5′-Oligoadenylate Synthetase Gene Cluster Is Associated with Human Predisposition to Tick-Borne Encephalitis Virus-Induced Disease. The Journal of Infectious Diseases 202: 1813–1818

4. Bastard P, Rosen LB, Zhang Q, Michailidis E, Hoffmann HH, Zhang Y, Dorgham K, Philippot Q, Rosain J, Béziat V et al (2020) Autoantibodies against type I IFNs in patients with life-threatening COVID-19. Science 370

5. Bloom GS, Brashear TA (1989) A Novel 58-kDa Protein Associates with the Golgi Apparatus and Microtubules. Journal of Biological Chemistry 264: 16083–16092

6. Borgese N, Aggujaro D, Carrera P, Pietrini G, Bassetti M (1996) A role for N-myristoylation in protein targeting: NADH-cytochrome b5 reductase requires myristic acid for association with outer mitochondrial but not ER membranes. Journal of Cell Biology 135: 1501–1513

7. Chen YG, Hur S (2022) Cellular origins of dsRNA, their recognition and consequences. Nature Reviews Molecular Cell Biology 23: 286–301

8. Consortium GT (2015) Human genomics. The Genotype-Tissue Expression (GTEx) pilot analysis: multitissue gene regulation in humans. Science 348: 648–660

9. Corman VM, Muth D, Niemeyer D, Drosten C (2018) Hosts and Sources of Endemic Human Coronaviruses. In: Advances in Virus Research, Kielian M., Mettenleiter T.C., Roossinck M.J. (eds.) pp. 163–188. Academic Press:

10. Courtney SC, Di H, Stockman BM, Liu H, Scherbik SV, Brinton MA (2012) Identification of Novel Host Cell Binding Partners of Oas1b, the Protein Conferring Resistance to Flavivirus-Induced Disease in Mice. Journal of Virology 86: 7953–7963

11. Culbertson EM, Levin TC (2023) Eukaryotic CD-NTase, STING, and viperin proteins evolved via domain shuffling, horizontal transfer, and ancient inheritance from prokaryotes. PLoS Biol 21: e3002436

12. Dalskov L, Gad HH, Hartmann R (2023) Viral recognition and the antiviral interferon response. The EMBO Journal 42: e112907

13. Danziger O, Patel RS, DeGrace EJ, Rosen MR, Rosenberg BR (2022) Inducible CRISPR activation screen for interferon-stimulated genes identifies OAS1 as a SARS-CoV-2 restriction factor. PLoS Pathog 18: e1010464

14. Deichaite I, Casson LP, Ling HP, Resh MD (1988) In vitro synthesis of pp60v-src: myristylation in a cell-free system. Mol Cell Biol 8: 4295–4301

15. Desai SY, Patel RC, Sen GC, Malhotra P, Ghadge GD, Thimmapaya B (1995) Activation of Interferon-inducible 2′ −5′ Oligoadenylate Synthetase by Adenoviral VAI RNA (∗). Journal of Biological Chemistry 270: 3454–3461

16. Desiere F, Deutsch EW, King NL, Nesvizhskii AI, Mallick P, Eng J, Chen S, Eddes J, Loevenich SN, Aebersold R (2006) The PeptideAtlas project. Nucleic Acids Res 34: D655–658

17. Diefenbacher MV, Baric TJ, Martinez DR, Baric RS, Catanzaro NJ, Sheahan TP (2024) A nano-luciferase expressing human coronavirus OC43 for countermeasure development. Virus Research 339: 199286

18. Dong B, Silverman RH (1995) 2-5A-dependent RNase Molecules Dimerize during Activation by 2-5A (∗). Journal of Biological Chemistry 270: 4133–4137

19. Donovan J, Dufner M, Korennykh A (2013) Structural basis for cytosolic double-stranded RNA surveillance by human oligoadenylate synthetase 1. Proceedings of the National Academy of Sciences 110: 1652–1657

20. Drosten C, Günther S, Preiser W, van der Werf S, Brodt HR, Becker S, Rabenau H, Panning M, Kolesnikova L, Fouchier RA et al (2003) Identification of a novel coronavirus in patients with severe acute respiratory syndrome. N Engl J Med 348: 1967–1976

21. Elbahesh H, Jha BK, Silverman RH, Scherbik SV, Brinton MA (2011) The Flvr-encoded murine oligoadenylate synthetase 1b (Oas1b) suppresses 2-5A synthesis in intact cells. Virology 409: 262–270

22. Elkhateeb E, Tag-El-Din-Hassan HT, Sasaki N, Torigoe D, Morimatsu M, Agui T (2016) The role of mouse 2′,5′-oligoadenylate synthetase 1 paralogs. Infection, Genetics and Evolution 45: 393–401

23. Froggatt HM, Harding AT, Chaparian RR, Heaton NS (2021) ETV7 limits antiviral gene expression and control of influenza viruses. Science Signaling 14: eabe1194

24. Galao RP, Le Tortorec A, Pickering S, Kueck T, Neil SJ (2012) Innate sensing of HIV-1 assembly by Tetherin induces NFkappaB-dependent proinflammatory responses. Cell Host Microbe 12: 633–644

25. Ganser-Pornillos BK, Pornillos O (2019) Restriction of HIV-1 and other retroviruses by TRIM5. Nat Rev Microbiol 17: 546–556

26. Gaunt ER, Hardie A, Claas EC, Simmonds P, Templeton KE (2010) Epidemiology and clinical presentations of the four human coronaviruses 229E, HKU1, NL63, and OC43 detected over 3 years using a novel multiplex real-time PCR method. J Clin Microbiol 48: 2940–2947

27. Ghosh A, Sarkar SN, Guo W, Bandyopadhyay S, Sen GC (1997) Enzymatic Activity of 2′–5′-Oligoadenylate Synthetase Is Impaired by Specific Mutations that Affect Oligomerization of the Protein*. Journal of Biological Chemistry 272: 33220–33226

28. Ghosh A, Sarkar SN, Sen GC (2000) Cell Growth Regulatory and Antiviral Effects of the P69 Isozyme of 2−5 (A) Synthetase. Virology 266: 319–328

29. Gokul A, Arumugam T, Ramsuran V (2023) Genetic Ethnic Differences in Human 2′-5′-Oligoadenylate Synthetase and Disease Associations: A Systematic Review. Genes 14: 527

30. Goldstein SA, Feeley TM, Babler KM, Hilbert ZA, Downhour DM, Moshiri N, Elde NC (2024) Hidden evolutionary constraints dictate the retention of coronavirus accessory genes. Current Biology 34: 5685–5696.e5683

31. Goldstein SA, Thornbrough JM, Zhang R, Jha BK, Li Y, Elliott R, Quiroz-Figueroa K, Chen AI, Silverman RH, Weiss SR (2017) Lineage A Betacoronavirus NS2 Proteins and the Homologous Torovirus Berne pp1a Carboxy-Terminal Domain Are Phosphodiesterases That Antagonize Activation of RNase L. Journal of Virology 91: 10.1128/jvi.02201-02216

32. Hagelauer E, Lotke R, Kmiec D, Hu D, Hohner M, Stopper S, Nchioua R, Kirchhoff F, Sauter D, Schindler M (2023) Tetherin Restricts SARS-CoV-2 despite the Presence of Multiple Viral Antagonists. Viruses 15

33. Hambleton S, Goodbourn S, Young DF, Dickinson P, Mohamad SMB, Valappil M, McGovern N, Cant AJ, Hackett SJ, Ghazal P et al (2013) STAT2 deficiency and susceptibility to viral illness in humans. Proceedings of the National Academy of Sciences 110: 3053–3058

34. Hardy A, Bakshi S, Furnon W, MacLean O, Gu Q, Varjak M, Varela M, Aziz MA, Shaw AE, Pinto RM et al (2023) The Timing and Magnitude of the Type I Interferon Response Are Correlated with Disease Tolerance in Arbovirus Infection. mBio 14: e0010123

35. Harioudh MK, Perez J, Chong Z, Nair S, So L, McCormick KD, Ghosh A, Shao L, Srivastava R, Soveg F et al (2024) Oligoadenylate synthetase 1 displays dual antiviral mechanisms in driving translational shutdown and protecting interferon production. Immunity 57: 446–461.e447

36. Hartmann R, Justesen J, Sarkar SN, Sen GC, Yee VC (2003) Crystal Structure of the 2′-Specific and Double-Stranded RNA-Activated Interferon-Induced Antiviral Protein 2′-5′-Oligoadenylate Synthetase. Molecular Cell 12: 1173–1185

37. Jackson CB, Farzan M, Chen B, Choe H (2022) Mechanisms of SARS-CoV-2 entry into cells. Nature Reviews Molecular Cell Biology 23: 3–20

38. Johnson DR, Bhatnagar RS, Knoll LJ, Gordon JI (1994) GENETIC AND BIOCHEMICAL STUDIES OF PROTEIN N-MYRISTOYLATION. Annual Review of Biochemistry 63: 869–914

39. Kjer-Hansen P, Weatheritt RJ (2023) The function of alternative splicing in the proteome: rewiring protein interactomes to put old functions into new contexts. Nat Struct Mol Biol 30: 1844–1856

40. Koul A, Deo S, Booy EP, Orriss GL, Genung M, McKenna SA (2020a) Impact of double-stranded RNA characteristics on the activation of human 2′–5′-oligoadenylate synthetase 2 (OAS2). Biochemistry and Cell Biology 98: 70–82

41. Koul A, Gemmill D, Lubna N, Meier M, Krahn N, Booy EP, Stetefeld J, Patel TR, McKenna SA (2020b) Structural and Hydrodynamic Characterization of Dimeric Human Oligoadenylate Synthetase 2. Biophysical Journal 118: 2726–2740

42. Koul A, Hui LT, Lubna N, McKenna SA (2024) Distinct domain organization and diversity of 2′-5′-oligoadenylate synthetases. Biochemistry and Cell Biology 102: 305–318

43. Kristiansen H, Gad HH, Eskildsen-Larsen S, Despres P, Hartmann R (2010) The Oligoadenylate Synthetase Family: An Ancient Protein Family with Multiple Antiviral Activities. Journal of Interferon & Cytokine Research 31: 41–47

44. Labun K, Montague TG, Krause M, Torres Cleuren YN, Tjeldnes H, Valen E (2019) CHOPCHOP v3: expanding the CRISPR web toolbox beyond genome editing. Nucleic Acids Res 47: W171–W174

45. Lamborn IT, Jing H, Zhang Y, Drutman SB, Abbott JK, Munir S, Bade S, Murdock HM, Santos CP, Brock LG et al (2017) Recurrent rhinovirus infections in a child with inherited MDA5 deficiency. J Exp Med 214: 1949–1972

46. Larsson A (2014) AliView: a fast and lightweight alignment viewer and editor for large datasets. Bioinformatics 30: 3276–3278

47. Latinne A, Hu B, Olival KJ, Zhu G, Zhang L-B, Li H, Chmura AA, Field HE, Zambrana-Torrelio C, Epstein JH et al (2024) Origin and cross-species transmission of bat coronaviruses in China. Nature Communications 15: 10705

48. Lee D, Le Pen J, Yatim A, Dong B, Aquino Y, Ogishi M, Pescarmona R, Talouarn E, Rinchai D, Zhang P et al (2023) Inborn errors of OAS-RNase L in SARS-CoV-2-related multisystem inflammatory syndrome in children. Science 379: eabo3627

49. Li Y, Banerjee S, Wang Y, Goldstein SA, Dong B, Gaughan C, Silverman RH, Weiss SR (2016) Activation of RNase L is dependent on OAS3 expression during infection with diverse human viruses. Proc Natl Acad Sci U S A 113: 2241–2246

50. Li Y, Slavik KM, Toyoda HC, Morehouse BR, de Oliveira Mann CC, Elek A, Levy S, Wang Z, Mears KS, Liu J et al (2023) cGLRs are a diverse family of pattern recognition receptors in innate immunity. Cell 186: 3261–3276 e3220

51. Lytras S, Wickenhagen A, Sugrue E, Stewart DG, Swingler S, Sims A, Jackson Ireland H, Davies EL, Ludlam EM, Li Z et al (2023) Resurrection of 2′-5′-oligoadenylate synthetase 1 (OAS1) from the ancestor of modern horseshoe bats blocks SARS-CoV-2 replication. PLOS Biology 21: e3002398

52. Mac Kain A, Maarifi G, Aicher S-M, Arhel N, Baidaliuk A, Munier S, Donati F, Vallet T, Tran QD, Hardy A et al (2022) Identification of DAXX as a restriction factor of SARS-CoV-2 through a CRISPR/Cas9 screen. Nature Communications 13: 2442

53. Madeira F, Madhusoodanan N, Lee J, Eusebi A, Niewielska A, Tivey ARN, Lopez R, Butcher S (2024) The EMBL-EBI Job Dispatcher sequence analysis tools framework in 2024. Nucleic acids research 52: W521–W525

54. Maitra RK, Li G, Xiao W, Dong B, Torrence PF, Silverman RH (1995) Catalytic Cleavage of an RNA Target by 2–5A Antisense and RNase L (∗). Journal of Biological Chemistry 270: 15071–15075

55. Marié I, Blanco J, Rebouillat D, Hovanessian AG (1997) 69-kDa and 100-kDa Isoforms of Interferon-Induced (2′-5′)Oligoadenylate Synthetase Exhibit Differential Catalytic Parameters. European Journal of Biochemistry 248: 558–566

56. Marié I, Hovanessian AG (1992) The 69-kDa 2-5A synthetase is composed of two homologous and adjacent functional domains. Journal of Biological Chemistry 267: 9933–9939

57. Marié I, Rebouillat D, Hovanessian AG (1999) The expression of both domains of the 69/71 kDa 2’,5’ oligoadenylate synthetase generates a catalytically active enzyme and mediates an anti-viral response. Eur J Biochem 262: 155–165

58. Marié I, Svab J, Robert N, Galabru J, Hovanessian AG (1990) Differential expression and distinct structure of 69– and 100-kDa forms of 2-5A synthetase in human cells treated with interferon. Journal of Biological Chemistry 265: 18601–18607

59. Marx S, Kummerer BM, Grutzner C, Kato H, Schlee M, Renn M, Bartok E, Hartmann G (2022) RIG-I-induced innate antiviral immunity protects mice from lethal SARS-CoV-2 infection. Mol Ther Nucleic Acids 27: 1225–1234

60. McDougal MB, De Maria AM, Ohlson MB, Kumar A, Xing C, Schoggins JW (2023) Interferon inhibits a model RNA virus via a limited set of inducible effector genes. EMBO reports 24: e56901

61. McEwan WA, Tam JC, Watkinson RE, Bidgood SR, Mallery DL, James LC (2013) Intracellular antibody-bound pathogens stimulate immune signaling via the Fc receptor TRIM21. Nat Immunol 14: 327–336

62. Meng EC, Goddard TD, Pettersen EF, Couch GS, Pearson ZJ, Morris JH, Ferrin TE (2023) UCSF ChimeraX: Tools for structure building and analysis. Protein Sci 32: e4792

63. Neil SJ (2013) The antiviral activities of tetherin. Curr Top Microbiol Immunol 371: 67–104

64. Perelygin AA, Scherbik SV, Zhulin IB, Stockman BM, Li Y, Brinton MA (2002) Positional cloning of the murine flavivirus resistance gene. Proceedings of the National Academy of Sciences 99: 9322–9327

65. Pertel T, Hausmann S, Morger D, Zuger S, Guerra J, Lascano J, Reinhard C, Santoni FA, Uchil PD, Chatel L et al (2011) TRIM5 is an innate immune sensor for the retrovirus capsid lattice. Nature 472: 361–365

66. Pfaender S, Mar KB, Michailidis E, Kratzel A, Boys IN, V’kovski P, Fan W, Kelly JN, Hirt D, Ebert N et al (2020) LY6E impairs coronavirus fusion and confers immune control of viral disease. Nature Microbiology 5: 1330–1339

67. Reineke LC, Lloyd RE (2015) The stress granule protein G3BP1 recruits protein kinase R to promote multiple innate immune antiviral responses. J Virol 89: 2575–2589

68. Rihn SJ, Aziz MA, Stewart DG, Hughes J, Turnbull ML, Varela M, Sugrue E, Herd CS, Stanifer M, Sinkins SP et al (2019) TRIM69 Inhibits Vesicular Stomatitis Indiana Virus. J Virol 93

69. Romero-Brey I, Bartenschlager R (2014) Membranous Replication Factories Induced by Plus-Strand RNA Viruses. Viruses 6: 2826–2857

70. Sa Ribero M, Jouvenet N, Dreux M, Nisole S (2020) Interplay between SARS-CoV-2 and the type I interferon response. PLoS Pathog 16: e1008737

71. Sanjana NE, Shalem O, Zhang F (2014) Improved vectors and genome-wide libraries for CRISPR screening. Nat Methods 11: 783–784

72. Sarkar SN, Bandyopadhyay S, Ghosh A, Sen GC (1999a) Enzymatic Characteristics of Recombinant Medium Isozyme of 2′-5′ Oligoadenylate Synthetase *. Journal of Biological Chemistry 274: 1848–1855

73. Sarkar SN, Ghosh A, Wang H-W, Sung S-S, Sen GC (1999b) The Nature of the Catalytic Domain of 2’-5’-Oligoadenylate Synthetases. Journal of Biological Chemistry 274: 25535–25542

74. Sarkar SN, Miyagi M, Crabb JW, Sen GC (2002) Identification of the Substrate-binding Sites of 2′-5′-Oligoadenylate Synthetase *. Journal of Biological Chemistry 277: 24321–24330

75. Sato S, Li K, Kameyama T, Hayashi T, Ishida Y, Murakami S, Watanabe T, Iijima S, Sakurai Y, Watashi K et al (2015) The RNA sensor RIG-I dually functions as an innate sensor and direct antiviral factor for hepatitis B virus. Immunity 42: 123–132

76. Schneider WM, Luna JM, Hoffmann HH, Sanchez-Rivera FJ, Leal AA, Ashbrook AW, Le Pen J, Ricardo-Lax I, Michailidis E, Peace A et al (2021) Genome-Scale Identification of SARS-CoV-2 and Pan-coronavirus Host Factor Networks. Cell 184: 120–132 e114

77. Schoggins JW (2019) Interferon-Stimulated Genes: What Do They All Do? Annual Review of Virology 6: 567–584

78. Schoggins JW, Wilson SJ, Panis M, Murphy MY, Jones CT, Bieniasz P, Rice CM (2011) A diverse range of gene products are effectors of the type I interferon antiviral response. Nature 472: 481–485

79. Schwartz SL, Conn GL (2019) RNA regulation of the antiviral protein 2′-5′-oligoadenylate synthetase. WIREs RNA 10: e1534

80. Shaw AE, Hughes J, Gu Q, Behdenna A, Singer JB, Dennis T, Orton RJ, Varela M, Gifford RJ, Wilson SJ et al (2017) Fundamental properties of the mammalian innate immune system revealed by multispecies comparison of type I interferon responses. PLOS Biology 15: e2004086

81. Shaw B, Gatherer D (2023) Candidate historical events for the emergence of Human Coronavirus OC43: A critical reassessment of the molecular evidence. PLoS One 18: e0285481

82. Silverman RH (2007) Viral encounters with 2’,5’-oligoadenylate synthetase and RNase L during the interferon antiviral response. J Virol 81: 12720–12729

83. Soveg FW, Schwerk J, Gokhale NS, Cerosaletti K, Smith JR, Pairo-Castineira E, Kell AM, Forero A, Zaver SA, Esser-Nobis K et al (2021) Endomembrane targeting of human OAS1 p46 augments antiviral activity. eLife 10: e71047

84. Tareen SU, Sawyer SL, Malik HS, Emerman M (2009) An expanded clade of rodent Trim5 genes. Virology 385: 473–483

85. Tate EW, Soday L, de la Lastra AL, Wang M, Lin H (2024) Protein lipidation in cancer: mechanisms, dysregulation and emerging drug targets. Nat Rev Cancer 24: 240–260

86. Thorne LG, Reuschl AK, Zuliani-Alvarez L, Whelan MVX, Turner J, Noursadeghi M, Jolly C, Towers GJ (2021) SARS-CoV-2 sensing by RIG-I and MDA5 links epithelial infection to macrophage inflammation. EMBO J 40: e107826

87. V’kovski P, Kratzel A, Steiner S, Stalder H, Thiel V (2021) Coronavirus biology and replication: implications for SARS-CoV-2. Nature Reviews Microbiology 19: 155–170

88. Varela M, Piras IM, Mullan C, Shi X, Tilston-Lunel NL, Pinto RM, Taggart A, Welch SR, Neil SJD, Kreher F et al (2017) Sensitivity to BST-2 restriction correlates with Orthobunyavirus host range. Virology 509: 121–130

89. Vijgen L, Keyaerts E, Moës E, Thoelen I, Wollants E, Lemey P, Vandamme A-M, Van Ranst M (2005) Complete Genomic Sequence of Human Coronavirus OC43: Molecular Clock Analysis Suggests a Relatively Recent Zoonotic Coronavirus Transmission Event. Journal of Virology 79: 1595–1604

90. Wickenhagen A, Sugrue E, Lytras S, Kuchi S, Noerenberg M, Turnbull ML, Loney C, Herder V, Allan J, Jarmson I et al (2021) A prenylated dsRNA sensor protects against severe COVID-19. Science 374: eabj3624

91. Zaki AM, van Boheemen S, Bestebroer TM, Osterhaus AD, Fouchier RA (2012) Isolation of a novel coronavirus from a man with pneumonia in Saudi Arabia. N Engl J Med 367: 1814–1820

92. Zhang SY, Jouanguy E, Ugolini S, Smahi A, Elain G, Romero P, Segal D, Sancho-Shimizu V, Lorenzo L, Puel A et al (2007) TLR3 deficiency in patients with herpes simplex encephalitis. Science 317: 1522–1527

93. Zhu J, Zhang Y, Ghosh A, Cuevas Rolando A, Forero A, Dhar J, Ibsen Mikkel S, Schmid-Burgk Jonathan L, Schmidt T, Ganapathiraju Madhavi K et al (2014) Antiviral Activity of Human OASL Protein Is Mediated by Enhancing Signaling of the RIG-I RNA Sensor. Immunity 40: 936–948

94. Zhu N, Zhang D, Wang W, Li X, Yang B, Song J, Zhao X, Huang B, Shi W, Lu R et al (2020) A Novel Coronavirus from Patients with Pneumonia in China, 2019. N Engl J Med 382: 727–733

