## Supplementary Figures for "Alternative splicing broadens antiviral diversity at the human *OAS2* locus"

[‡]These authors contributed equally

### Supplementary Information

**Supplementary Figure 1 - Scoring RNA virus infection by immunostaining dsRNA**

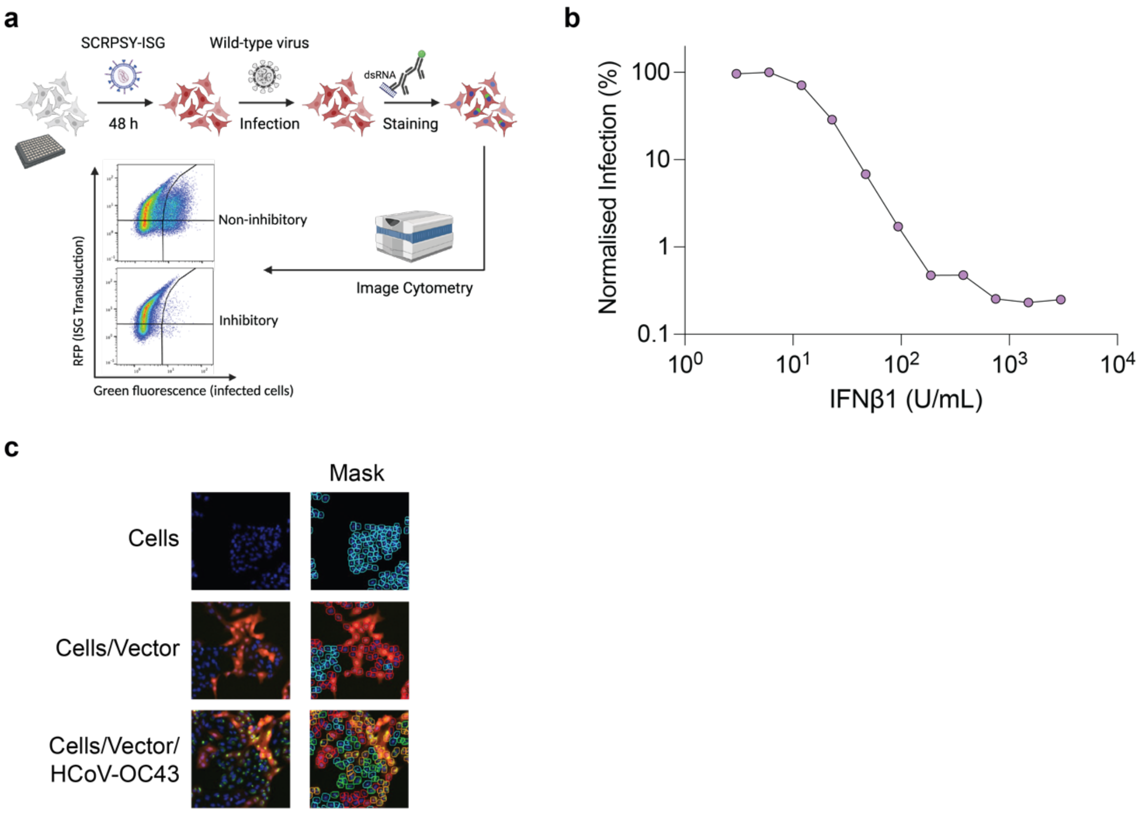


**Supplementary Figure 1 – Quantifying RNA virus infection by immunostaining dsRNA**. **a)** Schematic diagram of screening method used for the arrayed ISG expression screen used in (**Fig. 1d-g**). **b)** A549 cells were pre-treated with IFNβ1 for 24 h, infected with HCoV-OC43 for 72 h, and stained for dsRNA prior to quantifying dsRNA+ cells by image cytometry. **c)** Representative images of control wells in the ISG screen (**Fig. 1d**) gated for RFP and dsRNA expression levels using an image cytometer.

**Supplementary Figure 2**

**a**

| Peptide | OAS2 isoform | Instances | Cell types | Reference |
|---|---|---|---|---|
| VPTMQTPGSCGAR | p71 | 125 | HeLa, Jurkats | Zhang, J Proteome Research, 2018 |
| IHPIVNEMFSSR | p71 | 93 | HeLa, Jurkats, DCs | Rieckmann, Nat Immunology, 2017 |
| ILNNNSKR | p71 | 12 | Patient-derived xenografts (breast cancer) | Udeshi, Nat Comms, 2020 |
| DGTGNPIPPWKVPVK | p69 | 13 | HeLa | Zhang, J Proteome Research, 2018 |

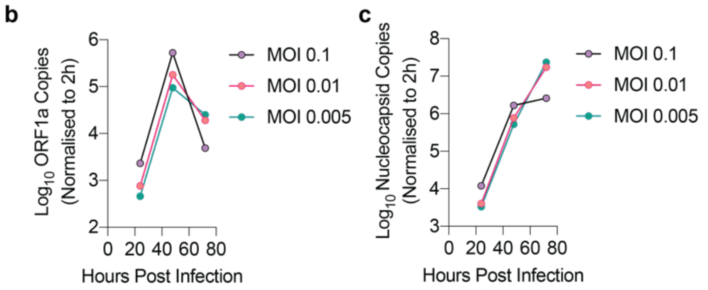


**Supplementary Figure 2 – Presence of endogenous OAS2 isoforms. a)** The Human Peptide Atlas (https://peptideatlas.org/) was searched for peptides from the OAS2 p71 and p69 C-terminus, generated by trypsin cleavage. **b)** HCoV-OC43 *ORF1a* transcript levels in A549 cells infected with HCoV-OC43 were quantified at multiple timepoints, by RT-qPCR. **c)** HCoV-OC43 *nucleocapsid* transcript levels in A549 cells infected with HCoV-OC43 were quantified at multiple timepoints, by RT-qPCR.

**Supplementary Figure 3 – N-terminal myristoylation of OAS2 is required for antiviral activity**

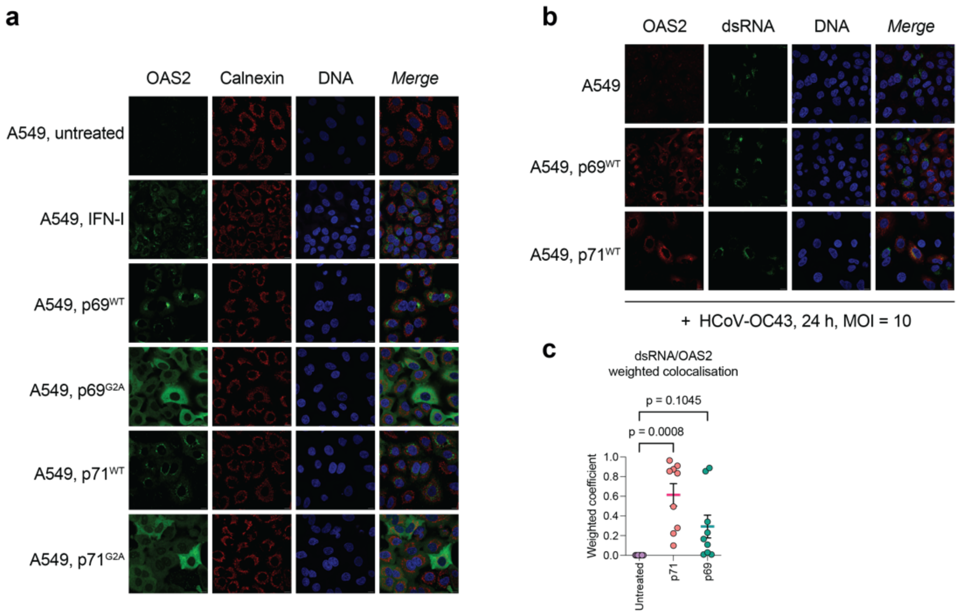


**Supplementary Figure 3 – N-terminal myristoylation activity is required for antiviral activity. a)** A549 cells stimulated with 1000 U/mL IFNβ, or A549 cells expressing p71, p69, $p71^{G2A}$ or $p69^{G2A}$, were immunostained with anti-OAS2 (green), anti-calnexin (red) and Hoechst (blue). **b)** Representative immunofluorescence images of A549 cells expressing p71 or p69 infected with HCoV-OC43 (MOI = 10) for 24 h and immunostained for OAS2 (red), dsRNA (green) and Hoechst (blue). **c)** Weighted coefficients between p71 and p69 isoforms and dsRNA; a one-way ANOVA test was used to assess significance, threshold p = 0.05. Each data point represents a separate region of interest from a representative experiment.

**Supplementary Figure 4 – Predicted RNA binding residues are necessary for OAS2 antiviral activity**

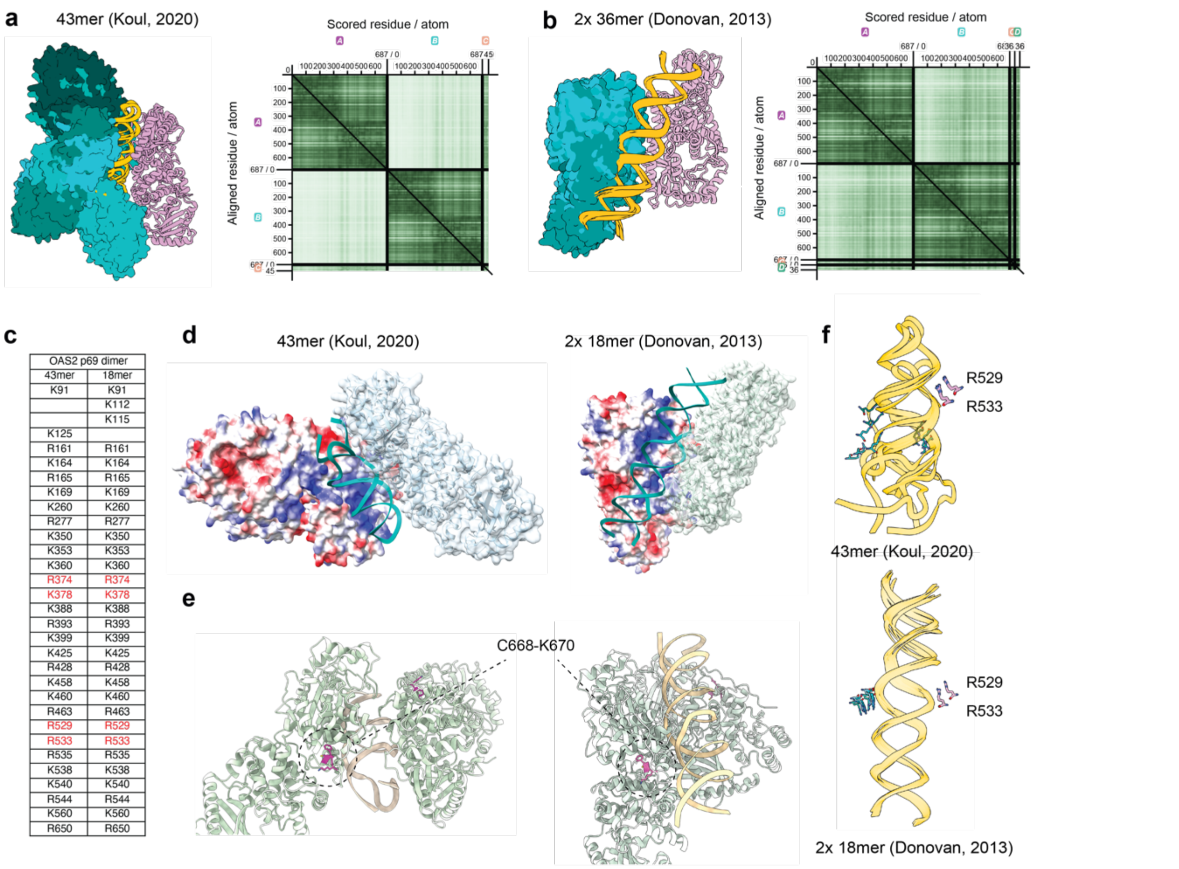


| OAS2 p69 dimer | |
|---|---|
| 43mer | 18mer |
| K91 | K91 |
| | K112 |
| | K115 |
| K125 | |
| R161 | R161 |
| K164 | K164 |
| R165 | R165 |
| K169 | K169 |
| K260 | K260 |
| R277 | R277 |
| K350 | K350 |
| K353 | K353 |
| K360 | K360 |
| R374 | R374 |
| K378 | K378 |
| K388 | K388 |
| R393 | R393 |
| K399 | K399 |
| K425 | K425 |
| R428 | R428 |
| K458 | K458 |
| K460 | K460 |
| R463 | R463 |
| R529 | R529 |
| R533 | R533 |
| R535 | R535 |
| K538 | K538 |
| K540 | K540 |
| R544 | R544 |
| K560 | K560 |
| R650 | R650 |

**Supplementary Figure 4. Predicted RNA binding residues are necessary for OAS2 antiviral activity. a,b)** Superposition of all 5 AlphaFold3 structural predictions of OAS2 p69 with each dsRNA, as in (**Fig. 4a,b**). One OAS2 chain was aligned across all 5 structures, to demonstrate the relative orientations of the second OAS2 chain. Predictions were more similar in (**b**) than in (**a**). Predicted Aligned Error (PAE) plots for the top-ranking structures are shown, which indicate high confidence in the conformation of individual OAS2 structures, and lower confidence in the relative position of one OAS2 chain to the other. **c)** List of basic amino acids within 5 Å of dsRNA in top-ranking structures. **d)** Electrostatic surface representation of top-ranking AlphaFold3 predictions of OAS2 p69-dsRNA structures from (**Fig. 4**), blue is basic, red is acidic. **e)** Location of triplet C668-K670 in top-ranking AlphaFold3 predictions of OAS2 p69-dsRNA structures from (**Fig. 4**), showing that these residues do not sit at a predicted OAS2 dimer interface. **f)** Superposition of all 5 AlphaFold3 predictions of OAS2 p69-dsRNA structures from (**Fig. 4**), showing just dsRNA and OAS2 residues R374, K378, R529 and R533 and their proximity in all structures.

**Supplementary Figure 5 – OAS2 C-terminal tails shape antiviral specificity**

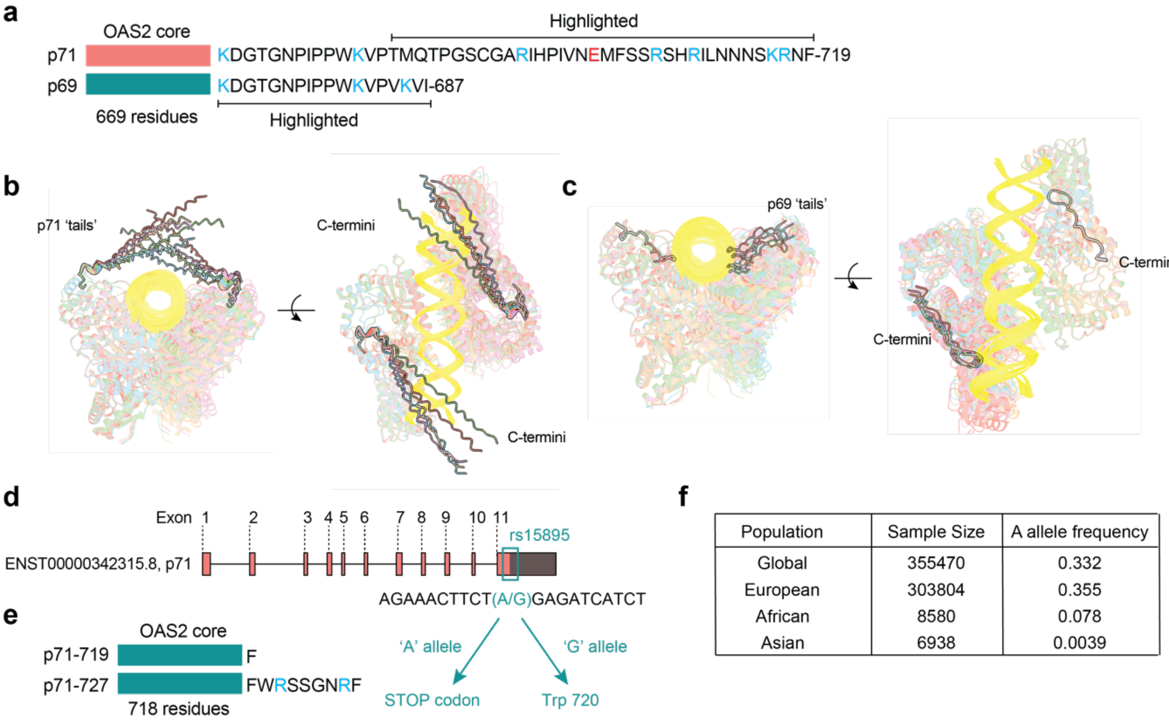


| Population | Sample Size | A allele frequency |
|---|---|---|
| Global | 355470 | 0.332 |
| European | 303804 | 0.355 |
| African | 8580 | 0.078 |
| Asian | 6938 | 0.0039 |

**Supplementary Figure 5 – OAS2 C-terminal tail shapes antiviral specificity. a)** OAS2 p69 and p71 C-terminal tails, indicating basic residues in blue, acidic residues in red. Regions highlighted in the structures in (**b-c**) are indicated. **b,c)** Superposition of all 5 OAS2 p71 (**b**) and p69 (**c**) AlphaFold3 structure predictions from (**Fig. 4a,b)**. The highlighted tail peptides begin at residues T684 for OAS2 p71, and K670 for p69, as indicated in (**a**); note that sequence K670-P683 is shared between p69 and p71 isoforms, thus the region highlighted for p69 is also partly present in p71. **d)** Representation of the exon structure of OAS2 p71 with SNP rs15895 resulting in a premature stop codon occurring in the canonical p71 sequence (NM_016817.3). Bright colours are coding sequence, shaded region is non-coding sequence. **e)** Schematic showing the sequences of the p71 C-termini resulting from SNP rs15895. **f)** Prevalence of the 'A' allele of SNP rs15895 recorded for different human populations, obtained from the National Centre for Biotechnology Information (NCBI), https://www.ncbi.nlm.nih.gov/snp/ (accessed 22 January 2025).
