## Supplementary Table 1 for "Alternative splicing broadens antiviral diversity at the human *OAS2* locus"

| Species | Unique No | Gene Name | Transduction_72h | VirusGrowth_72h | zscore | Miniscreen |
|---|---|---|---|---|---|---|
| Macaque | 930 | IFNB1 | 23.9 | 0.2 | -3.03 | yes |
| Macaque | 641 | IRF7 | 30.5 | 0.8 | -3.01 | yes |
| Macaque | 738 | PDGFRL | 86.1 | 1.0 | -3.00 | yes |
| Macaque | 640 | IRF7 | 24.7 | 1.2 | -3.00 | yes |
| Macaque | 701 | LY6E | 80.6 | 1.2 | -3.00 | yes |
| Macaque | 636 | ETV6 | 84.9 | 1.4 | -2.99 | yes |
| Macaque | 678 | OAS2 | 74.0 | 1.6 | -2.99 | yes |
| Cow | 1109 | IRF9 | 0.0 | 0.0 | -2.72 | no |
| Cow | 1001 | IFIH1 | 0.0 | 0.0 | -2.72 | no |
| Macaque | 967 | CTSS | 28.9 | 11.6 | -2.68 | yes |
| Macaque | 638 | TNFRSF10A | 15.4 | 13.3 | -2.63 | yes |
| Cow | 1055 | LY6E | 23.6 | 5.0 | -2.59 | yes |
| Macaque | 684 | TLK2 | 3.4 | 19.5 | -2.44 | no |
| Macaque | 692 | DDX58 | 31.2 | 19.8 | -2.43 | yes |
| Human | 11 | IFIH1 | 0.5 | 0.0 | -2.43 | no |
| Human | 99 | APOBEC3G | 0.0 | 0.0 | -2.43 | no |
| Human | 401 | PCGF5 | 0.0 | 0.0 | -2.43 | no |
| Human | 1928 | DRAM1 | 20.2 | 0.2 | -2.43 | yes |
| Macaque | 685 | TLK2 | 4.2 | 20.5 | -2.41 | no |
| Human | 192 | LY6E | 89.0 | 0.9 | -2.41 | yes |
| Macaque | 791 | BCL3 | 95.8 | 20.7 | -2.41 | yes |
| Human | 180 | DDX58 | 49.2 | 1.5 | -2.39 | yes |
| Human | 163 | OAS2 | 73.9 | 1.8 | -2.39 | yes |
| Human | 181 | MAP3K14 | 37.0 | 2.0 | -2.38 | yes |
| Human | 117 | IRF7 | 35.6 | 2.2 | -2.38 | yes |
| Human | 397 | IL28RA | 78.6 | 2.3 | -2.37 | yes |
| Human | 112 | ETV6 | 89.6 | 2.5 | -2.37 | yes |
| Human | 486 | PAX5 | 75.7 | 3.4 | -2.35 | yes |
| Human | 521 | ZBTB42 | 75.2 | 4.3 | -2.33 | yes |
| Human | 104 | LAMP3 | 94.2 | 4.8 | -2.31 | yes |
| Human | 427 | CTSS | 82.8 | 5.0 | -2.31 | yes |
| Cow | 1042 | OAS2 | 60.5 | 16.1 | -2.28 | yes |
| Human | 484 | RORB | 91.7 | 6.6 | -2.27 | yes |
| Human | 103 | LAMP3 | 90.1 | 6.7 | -2.27 | yes |
| Cow | 1258 | CHST1 | 61.7 | 17.3 | -2.25 | yes |
| Macaque | 796 | ARHGEF3 | 67.7 | 27.2 | -2.21 | yes |
| Human | 418 | PARP10 | 4.2 | 9.3 | -2.21 | no |
| Macaque | 570 | CMAH | 50.7 | 27.7 | -2.19 | yes |
| Macaque | 668 | SLFN12 | 0.8 | 28.7 | -2.16 | no |
| Human | 115 | TNFRSF10A | 13.6 | 11.5 | -2.15 | yes |
| Human | 529 | IRF4 | 95.0 | 12.1 | -2.14 | yes |
| Human | 278 | IRF1 | 75.1 | 13.2 | -2.11 | yes |
| Human | 293 | BCL3 | 96.5 | 14.0 | -2.09 | yes |
| Human | 169 | TLK2 | 2.8 | 14.1 | -2.09 | no |
| Human | 247 | RIPK2 | 92.3 | 15.4 | -2.06 | yes |
| Macaque | 611 | TNFSF10 | 79.6 | 33.3 | -2.02 | yes |
| Macaque | 633 | LAMP3 | 54.8 | 33.4 | -2.02 | yes |

| | | | | | | |
|---|---|---|---|---|---|---|
| Human | 286 | EHD4 | 65.1 | 17.6 | -2.00 | yes |
| Macaque | 826 | MICB | 75.9 | 36.2 | -1.94 | no |
| Cow | 1028 | SLFN12L | 1.0 | 28.9 | -1.94 | no |
| Cow | 1154 | MITD1 | 0.6 | 30.5 | -1.89 | no |
| Cow | 1012 | IFITM1 | 3.1 | 30.5 | -1.89 | no |
| Macaque | 699 | ANKFY1 | 17.8 | 38.6 | -1.86 | no |
| Cow | 1237 | MS4A8 | 35.2 | 31.6 | -1.86 | no |
| Macaque | 917 | TMEM173 | 85.0 | 38.7 | -1.86 | no |
| Macaque | 778 | IRF1 | 55.9 | 39.1 | -1.85 | no |
| Human | 303 | C5orf39 | 43.7 | 24.3 | -1.84 | no |
| Cow | 1278 | MAP2K6 | 89.8 | 33.4 | -1.81 | no |
| Cow | 1249 | ESR1 | 35.0 | 33.6 | -1.81 | no |
| Human | 188 | MAP3K5 | 55.2 | 26.3 | -1.79 | no |
| Human | 59 | SOCS1 | 94.7 | 26.4 | -1.79 | no |
| Macaque | 589 | RNF24 | 69.9 | 41.1 | -1.79 | no |
| Human | 187 | ANKFY1 | 46.7 | 27.2 | -1.77 | no |
| Human | 248 | RIPK2 | 89.6 | 27.7 | -1.76 | no |
| Macaque | 890 | MLKL | 4.0 | 43.3 | -1.72 | no |
| Cow | 1020 | SOCS1 | 73.2 | 36.9 | -1.72 | no |
| Human | 321 | FNDC3B | 24.3 | 30.5 | -1.69 | no |
| Cow | 1103 | TNFAIP3 | 41.7 | 38.0 | -1.69 | no |
| Human | 149 | SLFN12 | 1.7 | 30.7 | -1.69 | no |
| Macaque | 798 | C5orf39 | 2.1 | 44.6 | -1.68 | no |
| Macaque | 592 | SAT1 | 0.1 | 44.7 | -1.68 | no |
| Macaque | 943 | SAMD9L | 0.8 | 44.9 | -1.67 | no |
| Human | 313 | CEBPD | 90.9 | 31.6 | -1.66 | no |
| Cow | 1171 | HS3ST1 | 86.6 | 40.3 | -1.62 | no |
| Human | 335 | TNFAIP3 | 82.7 | 34.9 | -1.58 | no |
| Cow | 1123 | SAMD9 | 0.1 | 41.9 | -1.58 | no |
| Human | 518 | FAM111A | 0.9 | 35.3 | -1.57 | no |
| Human | 294 | GALNT2 | 63.9 | 35.6 | -1.56 | no |
| Cow | 1184 | PARM1 | 86.1 | 42.6 | -1.56 | no |
| Human | 218 | PIM3 | 86.1 | 35.9 | -1.56 | no |
| Cow | 1006 | OAS1 | 1.1 | 43.1 | -1.55 | no |
| Macaque | 582 | IFITM1 | 63.2 | 49.0 | -1.55 | no |
| Macaque | 926 | MOBKL2C | 62.0 | 49.5 | -1.53 | no |
| Human | 342 | B4GALT5 | 75.9 | 37.8 | -1.51 | no |
| Macaque | 893 | CLDN23 | 80.4 | 50.2 | -1.51 | no |
| Macaque | 547 | PSMB9 | 75.4 | 50.6 | -1.50 | no |
| Human | 86 | ETV7 | 91.0 | 38.5 | -1.50 | no |
| Human | 113 | NFIL3 | 80.6 | 40.2 | -1.45 | no |
| Macaque | 966 | HERC5 | 40.2 | 53.6 | -1.41 | no |
| Macaque | 726 | PIM3 | 87.8 | 53.9 | -1.40 | no |
| Human | 101 | IDO1 | 61.4 | 42.4 | -1.40 | no |
| Macaque | 828 | AKT3 | 68.2 | 55.1 | -1.36 | no |
| Cow | 1092 | IRF1 | 46.3 | 50.5 | -1.35 | no |
| Macaque | 918 | TMEM62 | 75.1 | 56.0 | -1.34 | no |
| Cow | 1095 | EHD4 | 57.0 | 51.2 | -1.33 | no |

| | | | | | |
|---|---|---|---|---|---|
| Human | 239 | CREB3L3 | 80.0 | 45.4 | -1.33 no |
| Human | 515 | GCNT1 | 89.9 | 45.4 | -1.33 no |
| Macaque | 612 | TNFSF10 | 41.7 | 56.4 | -1.32 no |
| Human | 353 | CD163 | 10.9 | 45.8 | -1.32 no |
| Human | 340 | MICB | 82.6 | 45.9 | -1.32 no |
| Cow | 1150 | ZFPM2 | 25.4 | 52.0 | -1.31 no |
| Macaque | 953 | HIST1H2BB | 82.4 | 57.0 | -1.30 no |
| Human | 47 | MAFF | 78.1 | 46.9 | -1.29 no |
| Macaque | 818 | LIPA | 60.1 | 57.5 | -1.29 no |
| Macaque | 711 | STARD5 | 70.6 | 57.6 | -1.29 no |
| Macaque | 569 | DEFB1 | 92.9 | 58.0 | -1.27 no |
| Cow | 1217 | DMRT2 | 16.7 | 53.7 | -1.26 no |
| Human | 45 | MAFF | 82.3 | 48.4 | -1.25 no |
| Human | 344 | AKT3 | 84.4 | 48.8 | -1.25 no |
| Human | 46 | MAFF | 83.8 | 50.3 | -1.21 no |
| Cow | 1142 | BTC | 63.2 | 55.7 | -1.20 no |
| Macaque | 561 | IFIH1 | 31.1 | 60.5 | -1.20 no |
| Human | 513 | ZFPM2 | 47.2 | 50.7 | -1.20 no |
| Human | 116 | RASGEF1B | 76.6 | 51.2 | -1.19 no |
| Cow | 1216 | B3GALT2 | 57.1 | 56.6 | -1.18 no |
| Human | 74 | CD80 | 88.0 | 51.5 | -1.18 no |
| Cow | 1191 | RNASEL | 0.3 | 56.9 | -1.17 no |
| Macaque | 815 | FNDC3B | 10.5 | 61.4 | -1.17 no |
| Human | 266 | HK2 | 18.1 | 53.1 | -1.14 no |
| Cow | 1010 | CXCL10 | 89.0 | 58.4 | -1.13 no |
| Human | 95 | G6PC | 94.1 | 53.7 | -1.13 no |
| Human | 417 | SAMD9L | 0.1 | 54.0 | -1.12 no |
| Cow | 1000 | RTP4 | 8.7 | 59.1 | -1.11 no |
| Macaque | 799 | IRF2 | 49.3 | 63.4 | -1.11 no |
| Human | 7 | MX1 | 58.4 | 54.5 | -1.11 no |
| Human | 530 | MOB3C | 82.0 | 54.5 | -1.11 no |
| Cow | 1007 | OAS1 | 59.2 | 59.5 | -1.10 no |
| Human | 411 | SLCO5A1 | 54.9 | 54.9 | -1.10 no |
| Human | 172 | STAT3 | 68.1 | 55.2 | -1.09 no |
| Human | 453 | BTC | 91.1 | 55.5 | -1.08 no |
| Human | 139 | FUT4 | 64.2 | 55.5 | -1.08 no |
| Macaque | 707 | ISG20 | 62.1 | 64.6 | -1.08 no |
| Human | 202 | STARD5 | 86.9 | 55.9 | -1.07 no |
| Human | 330 | IL15RA | 95.5 | 56.0 | -1.07 no |
| Macaque | 780 | HLA-X | 70.4 | 64.8 | -1.07 no |
| Macaque | 584 | C4orf32 | 14.7 | 65.3 | -1.05 no |
| Cow | 1299 | ZNF710 | 53.2 | 61.3 | -1.05 no |
| Human | 48 | SAT1 | 0.5 | 56.7 | -1.05 no |
| Human | 301 | ARHGEF3 | 78.8 | 56.7 | -1.05 no |
| Macaque | 660 | CEACAM1 | 57.9 | 65.4 | -1.05 no |
| Macaque | 653 | OASL | 12.4 | 65.7 | -1.04 no |
| Cow | 1014 | CXCL9 | 90.4 | 62.0 | -1.04 no |
| Cow | 1023 | TNFSF10 | 60.5 | 62.0 | -1.03 no |

| | | | | | | |
|---|---|---|---|---|---|---|
| Human | 284 | TREX1 | 88.4 | 57.5 | -1.03 | no |
| Cow | 1226 | PLCXD2 | 72.3 | 62.3 | -1.03 | no |
| Cow | 1308 | MYADM | 93.4 | 62.4 | -1.02 | no |
| Human | 147 | THBD | 68.4 | 58.3 | -1.01 | no |
| Macaque | 976 | HIST2H2AA4 | 81.3 | 66.7 | -1.01 | no |
| Macaque | 600 | COMMD3 | 85.0 | 66.7 | -1.01 | no |
| Cow | 1063 | XAF1 | 72.1 | 62.9 | -1.01 | no |
| Human | 93 | ARG2 | 91.7 | 59.4 | -0.99 | no |
| Human | 384 | OAS3 | 5.3 | 59.4 | -0.99 | no |
| Human | 140 | CYP1B1 | 85.4 | 59.6 | -0.98 | no |
| Human | 246 | FAM134B | 87.2 | 59.8 | -0.98 | no |
| Cow | 1169 | GTF2B | 0.1 | 64.2 | -0.98 | no |
| Cow | 1065 | P2RY6 | 2.8 | 64.6 | -0.96 | no |
| Macaque | 618 | MAFB | 95.1 | 68.3 | -0.96 | no |
| Human | 283 | SSBP3 | 86.7 | 60.5 | -0.96 | no |
| Macaque | 819 | BLZF1 | 83.5 | 68.5 | -0.95 | no |
| Cow | 1038 | GBP5 | 74.1 | 65.1 | -0.95 | no |
| Cow | 1064 | LGALS9 | 90.0 | 65.1 | -0.95 | no |
| Human | 508 | SLFN11 | 2.2 | 61.0 | -0.95 | no |
| Macaque | 909 | GUCA1C | 66.8 | 69.1 | -0.94 | no |
| Cow | 1153 | MOB3C | 65.9 | 65.7 | -0.93 | no |
| Macaque | 851 | GCA | 0.0 | 69.3 | -0.93 | no |
| Cow | 1107 | STAT1 | 64.5 | 65.8 | -0.93 | no |
| Human | 6 | IFITM3 | 58.8 | 61.7 | -0.93 | no |
| Macaque | 964 | BTN3A1 | 47.4 | 69.4 | -0.93 | no |
| Human | 199 | ISG20 | 76.6 | 62.2 | -0.92 | no |
| Human | 1934 | NLRC5 | 3.0 | 62.2 | -0.92 | no |
| Macaque | 913 | HIST1H2AC | 82.6 | 70.0 | -0.91 | no |
| Macaque | 773 | STAP1 | 76.6 | 70.0 | -0.91 | no |
| Human | 72 | TNFSF10 | 92.1 | 62.6 | -0.91 | no |
| Macaque | 642 | IRF7 | 64.7 | 70.1 | -0.91 | no |
| Macaque | 891 | CLDN23 | 75.7 | 70.1 | -0.91 | no |
| Human | 455 | BATF3 | 81.6 | 62.7 | -0.91 | no |
| Macaque | 585 | NRN1 | 88.5 | 70.3 | -0.90 | no |
| Macaque | 591 | MAFF | 79.8 | 70.3 | -0.90 | no |
| Macaque | 586 | NRN1 | 71.9 | 70.4 | -0.90 | no |
| Human | 403 | MLKL | 48.3 | 63.2 | -0.90 | no |
| Cow | 1207 | ACKR4 | 64.1 | 67.1 | -0.90 | no |
| Human | 125 | SERPING1 | 95.1 | 63.4 | -0.89 | no |
| Human | 168 | EXT1 | 68.2 | 63.7 | -0.88 | no |
| Macaque | 872 | MT1E | 81.3 | 71.1 | -0.88 | no |
| Macaque | 716 | SCO2 | 79.7 | 71.1 | -0.88 | no |
| Cow | 1291 | MGST1 | 85.5 | 67.9 | -0.87 | no |
| Cow | 1029 | FCGR1A | 87.6 | 67.9 | -0.87 | no |
| Macaque | 661 | CYP1B1 | 75.1 | 71.4 | -0.87 | no |
| Macaque | 639 | RASGEF1B | 50.6 | 71.6 | -0.86 | no |
| Cow | 1208 | ABHD1 | 72.1 | 68.5 | -0.86 | no |
| Macaque | 643 | SCARB2 | 36.7 | 71.8 | -0.86 | no |

| | | | | | | |
|---|---|---|---|---|---|---|
| Human | 1933 | IFITM3 | 80.5 | 65.2 | -0.85 | no |
| Macaque | 576 | CCL19 | 87.2 | 72.3 | -0.84 | no |
| Cow | 1215 | MIC1 | 82.9 | 69.3 | -0.84 | no |
| Human | 505 | RAD9A | 91.3 | 65.7 | -0.83 | no |
| Human | 275 | BLVRA | 88.2 | 65.8 | -0.83 | no |
| Human | 42 | RNF24 | 73.8 | 65.9 | -0.83 | no |
| Macaque | 710 | STARD5 | 74.1 | 72.8 | -0.83 | no |
| Human | 229 | APOL1 | 80.3 | 66.2 | -0.82 | no |
| Macaque | 635 | PLIN2 | 73.0 | 73.1 | -0.82 | no |
| Cow | 1254 | SULT1C4 | 9.2 | 70.3 | -0.81 | no |
| Human | 118 | SCARB2 | 88.3 | 66.8 | -0.81 | no |
| Macaque | 821 | IL15RA | 90.0 | 73.4 | -0.81 | no |
| Human | 88 | TFEC | 87.7 | 66.9 | -0.81 | no |
| Macaque | 765 | MASTL | 42.9 | 73.5 | -0.80 | no |
| Macaque | 916 | TAPBPL | 74.5 | 73.5 | -0.80 | no |
| Macaque | 957 | CA5B | 82.3 | 73.6 | -0.80 | no |
| Human | 367 | N4BP1 | 33.9 | 67.1 | -0.80 | no |
| Cow | 1022 | BATF2 | 80.4 | 70.7 | -0.80 | no |
| Macaque | 870 | RSAD2 | 56.1 | 73.9 | -0.79 | no |
| Cow | 1158 | ARNTL2 | 83.7 | 71.1 | -0.79 | no |
| Human | 182 | MOV10 | 0.1 | 67.6 | -0.79 | no |
| Human | 426 | HERC5 | 42.5 | 67.7 | -0.78 | no |
| Human | 5 | GBP1 | 86.1 | 67.7 | -0.78 | no |
| Human | 69 | GCH1 | 72.7 | 67.7 | -0.78 | no |
| Cow | 1119 | MLKL | 22.5 | 71.6 | -0.77 | no |
| Macaque | 982 | TSLP | 81.9 | 74.6 | -0.77 | no |
| Cow | 1253 | ICAM2 | 60.3 | 71.8 | -0.77 | no |
| Human | 222 | FFAR2 | 53.0 | 68.5 | -0.77 | no |
| Macaque | 766 | MASTL | 41.8 | 74.8 | -0.77 | no |
| Macaque | 951 | LDB1 | 62.8 | 74.9 | -0.76 | no |
| Macaque | 767 | HK2 | 18.1 | 74.9 | -0.76 | no |
| Macaque | 867 | ADAR | 75.2 | 75.0 | -0.76 | no |
| Cow | 1138 | RFX5 | 48.0 | 72.2 | -0.76 | no |
| Human | 215 | CASP7 | 90.7 | 68.9 | -0.75 | no |
| Macaque | 814 | HERC6 | 34.0 | 75.3 | -0.75 | no |
| Cow | 1099 | IRF2 | 43.0 | 72.5 | -0.75 | no |
| Human | 108 | ZBP1 | 74.4 | 69.2 | -0.75 | no |
| Human | 16 | IFI27 | 86.0 | 69.3 | -0.75 | no |
| Human | 290 | PPM1K | 80.7 | 69.5 | -0.74 | no |
| Macaque | 873 | MT1E | 67.7 | 75.6 | -0.74 | no |
| Macaque | 782 | TREX1 | 79.4 | 75.6 | -0.74 | no |
| Cow | 1091 | NMI | 1.1 | 72.9 | -0.74 | no |
| Macaque | 623 | ETV7 | 66.1 | 75.7 | -0.74 | no |
| Human | 23 | SNN | 90.9 | 69.9 | -0.73 | no |
| Cow | 1277 | CSRNP2 | 44.6 | 73.1 | -0.73 | no |
| Human | 65 | BATF2 | 95.6 | 70.2 | -0.73 | no |
| Human | 40 | NRN1 | 84.0 | 70.2 | -0.72 | no |
| Macaque | 888 | PRAP1 | 83.8 | 76.2 | -0.72 | no |

| | | | | | | |
|---|---|---|---|---|---|---|
| Macaque | 882 | ENPP2 | 63.6 | 76.5 | -0.71 | no |
| Cow | 1168 | FOXS1 | 87.0 | 74.0 | -0.71 | no |
| Cow | 1025 | TNFSF13B | 81.9 | 74.0 | -0.71 | no |
| Macaque | 863 | CLEC4D | 82.7 | 76.9 | -0.70 | no |
| Human | 366 | NCF1 | 91.4 | 71.3 | -0.70 | no |
| Human | 170 | TRIM56 | 6.5 | 71.3 | -0.70 | no |
| Macaque | 952 | LDB1 | 79.2 | 77.2 | -0.69 | no |
| Human | 382 | ATP10D | 53.0 | 71.5 | -0.69 | no |
| Macaque | 649 | ANGPTL1 | 65.5 | 77.3 | -0.69 | no |
| Macaque | 931 | RHOB | 43.3 | 77.3 | -0.69 | no |
| Macaque | 637 | NFIL3 | 25.3 | 77.4 | -0.68 | no |
| Human | 19 | MT1L | 91.9 | 71.9 | -0.68 | no |
| Human | 30 | CCL8 | 91.3 | 72.0 | -0.68 | no |
| Human | 485 | C5AR2 | 81.0 | 72.4 | -0.67 | no |
| Macaque | 939 | SAMD9 | 0.7 | 77.9 | -0.67 | no |
| Macaque | 737 | USP18 | 55.0 | 77.9 | -0.67 | no |
| Human | 120 | IFIT5 | 83.6 | 72.5 | -0.67 | no |
| Human | 220 | P2RY6 | 65.6 | 72.6 | -0.67 | no |
| Human | 43 | C9ofr19 | 86.1 | 72.6 | -0.67 | no |
| Cow | 1117 | RSAD2 | 74.1 | 75.6 | -0.66 | no |
| Human | 20 | DEFB1 | 91.4 | 72.7 | -0.66 | no |
| Human | 332 | UNC84B | 61.8 | 72.9 | -0.66 | no |
| Macaque | 971 | TCL6 | 87.9 | 78.3 | -0.66 | no |
| Cow | 1204 | ZNFX1 | 0.1 | 76.0 | -0.65 | no |
| Human | 24 | CCL5 | 95.7 | 73.1 | -0.65 | no |
| Human | 300 | TIMP1 | 91.7 | 73.1 | -0.65 | no |
| Macaque | 724 | XAF1 | 82.4 | 78.5 | -0.65 | no |
| Macaque | 992 | GMPR2 | 64.4 | 78.6 | -0.65 | no |
| Human | 314 | SAMD4A | 90.0 | 73.4 | -0.65 | no |
| Macaque | 920 | SLCO5A1 | 62.8 | 78.8 | -0.64 | no |
| Human | 216 | XAF1 | 88.0 | 73.6 | -0.64 | no |
| Macaque | 631 | IFI44L | 71.1 | 78.9 | -0.64 | no |
| Human | 308 | MT1F | 90.0 | 73.8 | -0.64 | no |
| Cow | 1019 | TMEM140 | 89.9 | 76.8 | -0.63 | no |
| Human | 107 | PUS1 | 65.7 | 74.0 | -0.63 | no |
| Human | 361 | CCND3 | 83.9 | 74.1 | -0.63 | no |
| Human | 393 | ABCA9 | 4.5 | 74.3 | -0.62 | no |
| Cow | 1083 | TAP2 | 3.8 | 77.1 | -0.62 | no |
| Macaque | 928 | APOBEC3F | 79.6 | 79.5 | -0.62 | no |
| Human | 1922 | RNF19B | 58.7 | 74.5 | -0.62 | no |
| Human | 531 | DNAJC1 | 62.6 | 74.5 | -0.62 | no |
| Human | 324 | LIPA | 71.6 | 74.7 | -0.62 | no |
| Cow | 1118 | ZC3HAV1 | 0.0 | 77.4 | -0.61 | no |
| Human | 76 | FBXO6 | 89.1 | 74.9 | -0.61 | no |
| Human | 143 | SPTLC2 | 67.4 | 75.0 | -0.61 | no |
| Macaque | 672 | AXUD1 | 61.5 | 80.0 | -0.61 | no |
| Human | 174 | PHF15 | 59.1 | 75.1 | -0.61 | no |
| Macaque | 801 | CCNA1 | 33.5 | 80.1 | -0.60 | no |

| | | | | | | |
|---|---|---|---|---|---|---|
| Human | 391 | CLEC4D | 91.1 | 75.4 | -0.60 | no |
| Human | 15 | OAS1 | 70.9 | 75.4 | -0.60 | no |
| Cow | 1132 | CD47 | 78.4 | 78.1 | -0.60 | no |
| Human | 493 | RAB20 | 94.9 | 75.5 | -0.60 | no |
| Macaque | 941 | CD58 | 48.8 | 80.5 | -0.59 | no |
| Macaque | 630 | IDO1 | 49.1 | 80.5 | -0.59 | no |
| Human | 358 | IRF9 | 74.9 | 75.7 | -0.59 | no |
| Cow | 1281 | DAXX | 34.0 | 78.3 | -0.59 | no |
| Cow | 1230 | LOC524181 | 75.1 | 78.4 | -0.59 | no |
| Macaque | 632 | IFI44L | 69.8 | 80.8 | -0.58 | no |
| Macaque | 949 | ARTS-1 | 45.5 | 80.8 | -0.58 | no |
| Cow | 1046 | DTX3L | 51.0 | 78.7 | -0.58 | no |
| Human | 327 | MT1A | 92.3 | 76.2 | -0.58 | no |
| Cow | 1133 | ERAP1 | 55.0 | 78.7 | -0.58 | no |
| Cow | 1126 | PSMB10 | 40.3 | 78.8 | -0.58 | no |
| Human | 60 | C10orf10 | 87.9 | 76.3 | -0.58 | no |
| Cow | 1021 | RNF114 | 77.2 | 78.9 | -0.57 | no |
| Human | 82 | CCDC92 | 89.5 | 76.4 | -0.57 | no |
| Human | 298 | TRIM25 | 92.8 | 76.5 | -0.57 | no |
| Cow | 1245 | CHI3L1 | 32.0 | 79.1 | -0.57 | no |
| Human | 323 | ADM | 86.6 | 76.7 | -0.57 | no |
| Human | 379 | AQP9 | 77.3 | 76.7 | -0.57 | no |
| Macaque | 959 | F3 | 72.0 | 81.4 | -0.56 | no |
| Human | 363 | PTMA | 71.7 | 76.8 | -0.56 | no |
| Human | 128 | C22orf28 | 68.6 | 77.0 | -0.56 | no |
| Human | 489 | GUCY1A3 | 2.5 | 77.1 | -0.56 | no |
| Human | 85 | SLFN12L | 87.3 | 77.2 | -0.55 | no |
| Human | 162 | DHX58 | 60.4 | 77.2 | -0.55 | no |
| Cow | 1077 | RNF19B | 50.0 | 79.7 | -0.55 | no |
| Macaque | 830 | ABLIM3 | 83.5 | 81.8 | -0.55 | no |
| Human | 231 | SERPINE1 | 85.4 | 77.4 | -0.55 | no |
| Human | 270 | MTHFD2L | 91.0 | 77.6 | -0.54 | no |
| Human | 41 | RNASE4 | 88.9 | 77.6 | -0.54 | no |
| Human | 150 | DCP1A | 82.5 | 77.8 | -0.54 | no |
| Human | 8 | PLSCR1 | 85.4 | 77.8 | -0.54 | no |
| Human | 375 | IFIT2 | 75.8 | 77.8 | -0.54 | no |
| Human | 190 | C2orf31 | 89.1 | 77.8 | -0.54 | no |
| Macaque | 899 | GSDMDC1 | 67.5 | 82.3 | -0.54 | no |
| Cow | 1213 | CITED1 | 81.6 | 80.3 | -0.54 | no |
| Cow | 1018 | BST2 | 42.4 | 80.3 | -0.53 | no |
| Human | 370 | MAX | 89.3 | 78.0 | -0.53 | no |
| Human | 173 | PLEKHA4 | 39.9 | 78.0 | -0.53 | no |
| Cow | 1214 | PI15 | 70.3 | 80.5 | -0.53 | no |
| Human | 129 | PRAME | 79.5 | 78.2 | -0.53 | no |
| Macaque | 675 | SAMHD1 | 73.9 | 82.7 | -0.53 | no |
| Human | 154 | AXUD1 | 81.4 | 78.4 | -0.52 | no |
| Human | 200 | FLJ39739 | 93.7 | 78.5 | -0.52 | no |
| Macaque | 934 | RXFP1 | 71.1 | 82.8 | -0.52 | no |

| | | | | | | |
|---|---|---|---|---|---|---|
| Cow | 1209 | CSDC2 | 84.2 | 80.9 | -0.52 | no |
| Macaque | 860 | GBP3 | 60.7 | 82.9 | -0.52 | no |
| Macaque | 845 | SLC25A30 | 72.7 | 82.9 | -0.52 | no |
| Cow | 1256 | C1H3orf38 | 12.4 | 81.2 | -0.51 | no |
| Human | 96 | MAB21L2 | 86.5 | 79.0 | -0.51 | no |
| Cow | 1225 | THEM6 | 31.6 | 81.3 | -0.51 | no |
| Cow | 1205 | GJA5 | 52.5 | 81.4 | -0.51 | no |
| Human | 214 | CD38 | 90.4 | 79.2 | -0.51 | no |
| Cow | 1267 | GDAP2 | 80.4 | 81.5 | -0.50 | no |
| Macaque | 613 | TNFSF13B | 76.9 | 83.4 | -0.50 | no |
| Cow | 1246 | IL31RA | 78.6 | 81.5 | -0.50 | no |
| Cow | 1201 | TIFA | 69.6 | 81.7 | -0.50 | no |
| Human | 328 | MT1H | 91.6 | 79.6 | -0.50 | no |
| Macaque | 566 | IFI27 | 78.0 | 83.7 | -0.50 | no |
| Human | 331 | DNAPTP6 | 73.1 | 79.8 | -0.49 | no |
| Human | 105 | SMAD3 | 83.4 | 80.0 | -0.49 | no |
| Human | 100 | FAM46C | 79.3 | 80.1 | -0.48 | no |
| Cow | 1157 | ARHGAP26 | 1.5 | 82.2 | -0.48 | no |
| Cow | 1220 | FLVCR2 | 85.8 | 82.4 | -0.48 | no |
| Cow | 1234 | GXYLT2 | 91.4 | 82.4 | -0.48 | no |
| Macaque | 578 | CCL8 | 45.2 | 84.4 | -0.47 | no |
| Macaque | 575 | CXCL10 | 75.3 | 84.5 | -0.47 | no |
| Cow | 1147 | ICAM1 | 40.5 | 82.7 | -0.47 | no |
| Cow | 1224 | FEZ1 | 45.0 | 82.8 | -0.47 | no |
| Human | 44 | LMO2 | 88.3 | 80.7 | -0.47 | no |
| Human | 362 | ODC1 | 70.4 | 81.0 | -0.46 | no |
| Macaque | 795 | TIMP1 | 84.4 | 84.9 | -0.46 | no |
| Human | 351 | STAT2 | 68.7 | 81.1 | -0.46 | no |
| Human | 412 | TMEM229B | 92.9 | 81.3 | -0.46 | no |
| Human | 337 | NT5C3 | 83.7 | 81.3 | -0.45 | no |
| Human | 49 | BST2 | 60.9 | 81.4 | -0.45 | no |
| Macaque | 988 | HSPA6 | 48.4 | 85.1 | -0.45 | no |
| Macaque | 721 | CASP7 | 80.5 | 85.2 | -0.45 | no |
| Macaque | 937 | HORMAD2 | 82.4 | 85.2 | -0.45 | no |
| Human | 207 | PMM2 | 77.8 | 81.6 | -0.45 | no |
| Human | 446 | SSTR2 | 87.3 | 81.6 | -0.45 | no |
| Human | 281 | HLA-E | 79.1 | 81.9 | -0.44 | no |
| Human | 273 | STAP1 | 85.8 | 81.9 | -0.44 | no |
| Cow | 1102 | IL15RA | 83.9 | 84.0 | -0.44 | no |
| Human | 341 | DUSP5 | 75.0 | 82.1 | -0.44 | no |
| Human | 354 | FLT1 | 61.1 | 82.2 | -0.43 | no |
| Human | 292 | LGMN | 86.3 | 82.2 | -0.43 | no |
| Human | 1924 | RTP4 | 81.1 | 82.3 | -0.43 | no |
| Human | 306 | IRF2 | 70.4 | 82.4 | -0.43 | no |
| Macaque | 986 | AXL | 65.0 | 85.9 | -0.43 | no |
| Cow | 1075 | TRIM5 | 3.9 | 84.4 | -0.42 | no |
| Cow | 1156 | ANKRD35 | 1.7 | 84.4 | -0.42 | no |
| Macaque | 654 | GTPBP2 | 54.3 | 86.1 | -0.42 | no |

| | | | | | | |
|---|---|---|---|---|---|---|
| Human | 28 | CCL19 | 91.0 | 82.6 | -0.42 | no |
| Human | 26 | CXCL11 | 92.1 | 82.7 | -0.42 | no |
| Cow | 1269 | CNDP1 | 48.5 | 84.5 | -0.42 | no |
| Macaque | 963 | BTN3A2 | 61.0 | 86.1 | -0.42 | no |
| Human | 189 | S100A8 | 90.9 | 82.7 | -0.42 | no |
| Cow | 1164 | CDKN2AIP | 79.9 | 84.6 | -0.42 | no |
| Cow | 1262 | DCP2 | 67.6 | 84.6 | -0.42 | no |
| Macaque | 940 | CD58 | 67.7 | 86.2 | -0.42 | no |
| Human | 4 | IFI6 | 63.0 | 82.8 | -0.42 | no |
| Macaque | 892 | CLDN23 | 71.1 | 86.3 | -0.41 | no |
| Cow | 1279 | LMO6 | 3.4 | 84.8 | -0.41 | no |
| Macaque | 568 | PMAIP1 | 88.0 | 86.4 | -0.41 | no |
| Macaque | 549 | PSMB8 | 0.0 | 86.4 | -0.41 | no |
| Cow | 1057 | ISG15 | 73.8 | 84.9 | -0.41 | no |
| Human | 194 | UBE2L6 | 92.8 | 83.1 | -0.41 | no |
| Human | 134 | PFKFB3 | 88.5 | 83.1 | -0.41 | no |
| Human | 102 | IFI44L | 70.1 | 83.2 | -0.41 | no |
| Macaque | 950 | PSMB10 | 78.2 | 86.6 | -0.41 | no |
| Cow | 1094 | TREX1 | 72.6 | 85.1 | -0.41 | no |
| Human | 127 | SIRPA | 85.9 | 83.4 | -0.40 | no |
| Human | 1914 | BTN3A3 | 45.8 | 83.5 | -0.40 | no |
| Human | 467 | CXCL16 | 85.6 | 83.6 | -0.40 | no |
| Cow | 1218 | ATAD1 | 58.6 | 85.7 | -0.39 | no |
| Macaque | 842 | CCND3 | 58.0 | 87.2 | -0.39 | no |
| Cow | 1252 | BHLHE41 | 76.4 | 85.8 | -0.39 | no |
| Cow | 1090 | MYD88 | 89.8 | 85.8 | -0.39 | no |
| Cow | 1251 | BHLHE41 | 59.1 | 86.0 | -0.38 | no |
| Human | 157 | SAMHD1 | 68.4 | 84.3 | -0.38 | no |
| Human | 390 | MB21D1 | 73.7 | 84.4 | -0.38 | no |
| Macaque | 836 | PRKD2 | 51.8 | 87.5 | -0.38 | no |
| Human | 250 | ULK4 | 75.8 | 84.6 | -0.37 | no |
| Cow | 1140 | CMPK2 | 10.7 | 86.3 | -0.37 | no |
| Macaque | 708 | ISG20 | 74.7 | 87.7 | -0.37 | no |
| Cow | 1135 | TAPBP | 67.2 | 86.4 | -0.37 | no |
| Human | 304 | RASSF4 | 85.2 | 84.8 | -0.37 | no |
| Human | 3 | PSMB8 | 87.1 | 84.8 | -0.37 | no |
| Cow | 1242 | PVRL2 | 83.5 | 86.4 | -0.37 | no |
| Cow | 1259 | IRF3 | 0.0 | 86.5 | -0.37 | no |
| Macaque | 650 | SERPING1 | 74.7 | 88.1 | -0.36 | no |
| Human | 279 | GJA4 | 82.9 | 85.2 | -0.36 | no |
| Human | 33 | B2M | 91.6 | 85.2 | -0.36 | no |
| Human | 260 | ZNF107 | 10.5 | 85.5 | -0.35 | no |
| Cow | 1104 | FAM46A | 23.5 | 87.1 | -0.35 | no |
| Human | 167 | DTX3L | 73.8 | 85.6 | -0.35 | no |
| Human | 264 | CMTR1 | 75.2 | 85.7 | -0.35 | no |
| Macaque | 657 | PCTK2 | 74.5 | 88.7 | -0.34 | no |
| Human | 193 | LINCR | 90.8 | 86.0 | -0.34 | no |
| Macaque | 579 | GLRX | 66.0 | 88.8 | -0.34 | no |

| | | | | | | |
|---|---|---|---|---|---|---|
| Human | 533 | SERTAD1 | 60.2 | 86.1 | -0.34 | no |
| Macaque | 925 | TMEM229B | 83.0 | 89.0 | -0.34 | no |
| Human | 1930 | PLAC8 | 90.2 | 86.2 | -0.34 | no |
| Human | 36 | CXCL9 | 93.1 | 86.3 | -0.33 | no |
| Macaque | 822 | DNAPTP6 | 66.1 | 89.1 | -0.33 | no |
| Macaque | 689 | PLEKHA4 | 43.1 | 89.2 | -0.33 | no |
| Human | 158 | NDC80 | 79.0 | 86.5 | -0.33 | no |
| Macaque | 663 | HPSE | 81.2 | 89.2 | -0.33 | no |
| Macaque | 874 | MT1E | 74.0 | 89.3 | -0.32 | no |
| Human | 32 | VAMP5 | 84.5 | 86.7 | -0.32 | no |
| Macaque | 727 | PNRC1 | 88.2 | 89.3 | -0.32 | no |
| Human | 67 | CCDC109B | 90.0 | 86.8 | -0.32 | no |
| Macaque | 656 | PFKFB3 | 80.3 | 89.5 | -0.32 | no |
| Macaque | 581 | B2M | 72.5 | 89.6 | -0.31 | no |
| Cow | 1295 | FLT3LG | 65.9 | 88.5 | -0.31 | no |
| Macaque | 954 | THEMIS2 | 71.8 | 89.7 | -0.31 | no |
| Human | 63 | CD9 | 82.5 | 87.2 | -0.31 | no |
| Cow | 1078 | GBP4 | 19.3 | 88.6 | -0.31 | no |
| Macaque | 837 | DDX60 | 30.8 | 89.8 | -0.31 | no |
| Macaque | 753 | TRIM5 | 51.1 | 89.8 | -0.31 | no |
| Macaque | 981 | TNFSF18 | 86.8 | 89.8 | -0.31 | no |
| Macaque | 933 | RXFP1 | 51.4 | 89.8 | -0.31 | no |
| Macaque | 898 | GIMAP2 | 80.0 | 89.9 | -0.31 | no |
| Macaque | 843 | ODC1 | 53.2 | 89.9 | -0.31 | no |
| Macaque | 850 | SLC16A1 | 57.5 | 89.9 | -0.31 | no |
| Human | 210 | SPSB1 | 86.6 | 87.5 | -0.30 | no |
| Cow | 1275 | KRT75 | 48.7 | 88.9 | -0.30 | no |
| Human | 297 | TRAFD1 | 73.2 | 87.6 | -0.30 | no |
| Cow | 1034 | TRIM14 | 5.6 | 89.0 | -0.30 | no |
| Cow | 1174 | JSP.1 | 25.3 | 89.0 | -0.30 | no |
| Human | 326 | BLZF1 | 79.8 | 87.7 | -0.30 | no |
| Cow | 1173 | IQCD | 30.4 | 89.2 | -0.29 | no |
| Cow | 1297 | CLEC2D | 74.1 | 89.2 | -0.29 | no |
| Cow | 1222 | EDN1 | 81.5 | 89.3 | -0.29 | no |
| Human | 443 | RFX5 | 95.0 | 88.0 | -0.29 | no |
| Cow | 1054 | UBA7 | 54.3 | 89.3 | -0.29 | no |
| Macaque | 958 | SLC2A12 | 59.0 | 90.5 | -0.29 | no |
| Human | 203 | RAB27A | 31.0 | 88.1 | -0.29 | no |
| Macaque | 563 | CCDC75 | 28.9 | 90.6 | -0.29 | no |
| Human | 348 | STAT1 | 84.4 | 88.3 | -0.28 | no |
| Human | 525 | SHOX2 | 75.2 | 88.3 | -0.28 | no |
| Human | 71 | TNFAIP6 | 85.1 | 88.3 | -0.28 | no |
| Cow | 1016 | SAT1 | 0.6 | 89.6 | -0.28 | no |
| Cow | 1264 | TNFRSF1B | 91.8 | 89.6 | -0.28 | no |
| Human | 211 | STEAP4 | 86.5 | 88.4 | -0.28 | no |
| Human | 501 | EGF | 76.6 | 88.4 | -0.28 | no |
| Cow | 1273 | C2 | 9.7 | 90.0 | -0.27 | no |
| Cow | 1120 | IL18BP | 87.2 | 90.1 | -0.27 | no |

| | | | | | | |
|---|---|---|---|---|---|---|
| Human | 269 | FER1L3 | 37.9 | 88.9 | -0.27 | no |
| Cow | 1241 | ADRB2 | 36.2 | 90.1 | -0.27 | no |
| Human | 89 | LRG1 | 90.7 | 88.9 | -0.27 | no |
| Cow | 1084 | MX2 | 60.2 | 90.2 | -0.27 | no |
| Macaque | 732 | USP18 | 77.3 | 91.2 | -0.27 | no |
| Human | 299 | IL1RN | 90.6 | 89.0 | -0.27 | no |
| Cow | 1219 | F2RL1 | 52.8 | 90.2 | -0.27 | no |
| Human | 77 | GEM | 86.3 | 89.4 | -0.26 | no |
| Macaque | 573 | CCL4 | 89.4 | 91.6 | -0.26 | no |
| Macaque | 729 | HLA-X | 67.0 | 91.6 | -0.25 | no |
| Cow | 1289 | NUAK2 | 74.8 | 90.7 | -0.25 | no |
| Macaque | 625 | MKX | 38.0 | 91.7 | -0.25 | no |
| Cow | 1041 | DHX58 | 42.6 | 90.8 | -0.25 | no |
| Human | 54 | COMMD3 | 92.0 | 89.8 | -0.25 | no |
| Human | 291 | CPT1A | 70.7 | 89.8 | -0.25 | no |
| Cow | 1244 | TBX20 | 82.5 | 91.0 | -0.25 | no |
| Macaque | 886 | HIST1H3D | 72.2 | 91.9 | -0.25 | no |
| Human | 68 | LGALS3 | 80.7 | 89.9 | -0.24 | no |
| Macaque | 764 | PNPT1 | 27.0 | 92.0 | -0.24 | no |
| Human | 392 | LEPR | 46.8 | 90.0 | -0.24 | no |
| Cow | 1260 | RUNX3 | 65.3 | 91.1 | -0.24 | no |
| Human | 81 | BCL2L14 | 85.6 | 90.1 | -0.24 | no |
| Macaque | 820 | FNDC4 | 90.1 | 92.1 | -0.24 | no |
| Cow | 995 | IFI6 | 74.4 | 91.2 | -0.24 | no |
| Cow | 1294 | MAD2L2 | 28.7 | 91.2 | -0.24 | no |
| Human | 165 | SP110 | 82.2 | 90.3 | -0.24 | no |
| Human | 195 | KIAA0040 | 95.8 | 90.3 | -0.24 | no |
| Macaque | 900 | IL20RB | 73.1 | 92.3 | -0.23 | no |
| Human | 360 | IL15 | 88.8 | 90.5 | -0.23 | no |
| Macaque | 783 | TREX1 | 71.3 | 92.7 | -0.22 | no |
| Macaque | 794 | IL1RN | 63.5 | 92.8 | -0.22 | no |
| Human | 387 | WHDC1 | 86.9 | 91.0 | -0.22 | no |
| Macaque | 688 | PLEKHA4 | 43.5 | 92.8 | -0.22 | no |
| Cow | 1097 | TRAFD1 | 8.2 | 92.0 | -0.22 | no |
| Human | 511 | CA13 | 92.5 | 91.1 | -0.22 | no |
| Human | 53 | ANKRD22 | 80.7 | 91.1 | -0.22 | no |
| Cow | 1266 | ZDHHC14 | 89.0 | 92.1 | -0.21 | no |
| Cow | 997 | GBP1 | 61.1 | 92.1 | -0.21 | no |
| Human | 121 | TYMP | 48.3 | 91.2 | -0.21 | no |
| Human | 183 | UBA7 | 45.1 | 91.2 | -0.21 | no |
| Cow | 1238 | TM6SF2 | 76.1 | 92.2 | -0.21 | no |
| Human | 503 | CASP10 | 88.0 | 91.3 | -0.21 | no |
| Human | 258 | MX2 | 58.9 | 91.3 | -0.21 | no |
| Macaque | 789 | CPT1A | 41.0 | 93.1 | -0.21 | no |
| Macaque | 932 | LIPC | 81.7 | 93.1 | -0.21 | no |
| Cow | 1040 | GBP2 | 36.9 | 92.3 | -0.21 | no |
| Human | 309 | MT1M | 93.8 | 91.5 | -0.21 | no |
| Macaque | 596 | HESX1 | 78.9 | 93.2 | -0.21 | no |

| | | | | | | |
|---|---|---|---|---|---|---|
| Macaque | 858 | FLJ23556 | 87.5 | 93.2 | -0.21 | no |
| Cow | 1074 | IFIT1 | 72.1 | 92.5 | -0.20 | no |
| Macaque | 781 | SSBP3 | 67.5 | 93.4 | -0.20 | no |
| Macaque | 624 | APOL6 | 74.6 | 93.4 | -0.20 | no |
| Cow | 1235 | BPNT1 | 67.5 | 92.8 | -0.20 | no |
| Human | 372 | GCA | 75.4 | 92.0 | -0.20 | no |
| Macaque | 733 | USP18 | 68.8 | 93.6 | -0.19 | no |
| Human | 236 | IFI44 | 78.1 | 92.1 | -0.19 | no |
| Macaque | 715 | SCO2 | 66.7 | 93.7 | -0.19 | no |
| Macaque | 835 | PRKD2 | 56.6 | 93.8 | -0.19 | no |
| Human | 311 | THOC4 | 86.7 | 92.3 | -0.19 | no |
| Cow | 1062 | CASP7 | 71.6 | 93.1 | -0.19 | no |
| Macaque | 878 | IGFBP3 | 71.0 | 93.9 | -0.19 | no |
| Macaque | 965 | BTN3A1 | 53.9 | 93.9 | -0.19 | no |
| Human | 274 | NAPA | 92.6 | 92.4 | -0.19 | no |
| Cow | 1195 | SP140L | 53.9 | 93.2 | -0.19 | no |
| Cow | 996 | GBP1 | 35.3 | 93.2 | -0.18 | no |
| Human | 179 | ENPP1 | 52.2 | 92.5 | -0.18 | no |
| Macaque | 960 | HRASLS2 | 80.9 | 94.0 | -0.18 | no |
| Human | 1926 | OAS1(p42) | 48.3 | 92.6 | -0.18 | no |
| Macaque | 833 | STAT1 | 53.2 | 94.1 | -0.18 | no |
| Macaque | 839 | IRF9 | 71.7 | 94.1 | -0.18 | no |
| Macaque | 922 | SLC16A4 | 72.2 | 94.2 | -0.18 | no |
| Cow | 1292 | RGS16 | 79.6 | 93.6 | -0.17 | no |
| Human | 78 | EPSTI1 | 73.6 | 92.9 | -0.17 | no |
| Macaque | 906 | C9orf52 | 50.7 | 94.4 | -0.17 | no |
| Cow | 1233 | DDIT4L | 31.1 | 93.8 | -0.17 | no |
| Macaque | 739 | TXNIP | 17.8 | 94.5 | -0.17 | no |
| Human | 217 | LGALS9 | 94.2 | 93.2 | -0.17 | no |
| Human | 1942 | DNAJC13 | 0.7 | 93.3 | -0.16 | no |
| Human | 307 | CCNA1 | 75.3 | 93.4 | -0.16 | no |
| Human | 133 | PFKFB3 | 91.9 | 93.4 | -0.16 | no |
| Macaque | 571 | C15orf48 | 81.4 | 94.8 | -0.16 | no |
| Macaque | 725 | LGALS9 | 86.0 | 94.9 | -0.16 | no |
| Cow | 1255 | C1H21orf91 | 1.0 | 94.3 | -0.16 | no |
| Cow | 1247 | VCPIP1 | 25.7 | 94.3 | -0.16 | no |
| Cow | 1155 | AKAP7 | 69.5 | 94.3 | -0.16 | no |
| Human | 439 | ERAP1 | 83.7 | 93.7 | -0.15 | no |
| Cow | 1210 | KDR | 21.5 | 94.4 | -0.15 | no |
| Macaque | 681 | IFI16 | 42.2 | 95.0 | -0.15 | no |
| Human | 488 | GALNT3 | 3.4 | 93.9 | -0.15 | no |
| Cow | 1125 | PARP9 | 0.6 | 94.7 | -0.15 | no |
| Cow | 1151 | IL7 | 74.1 | 94.7 | -0.14 | no |
| Macaque | 970 | c5orf56 | 86.2 | 95.3 | -0.14 | no |
| Human | 409 | TMEM62 | 74.0 | 94.1 | -0.14 | no |
| Human | 534 | ATAD1 | 70.2 | 94.2 | -0.14 | no |
| Human | 110 | PLIN2 | 83.6 | 94.2 | -0.14 | no |
| Macaque | 744 | FKBP5 | 31.9 | 95.3 | -0.14 | no |

| | | | | | | |
|---|---|---|---|---|---|---|
| Human | 491 | ICAM1 | 84.9 | 94.3 | -0.14 | no |
| Human | 242 | PCTK3 | 93.4 | 94.4 | -0.14 | no |
| Macaque | 904 | IL18BP | 88.9 | 95.5 | -0.14 | no |
| Macaque | 857 | JAK2 | 55.0 | 95.5 | -0.14 | no |
| Macaque | 690 | PHF15 | 41.1 | 95.5 | -0.14 | no |
| Macaque | 735 | USP18 | 66.5 | 95.6 | -0.13 | no |
| Human | 221 | IGFBP2 | 86.0 | 94.6 | -0.13 | no |
| Human | 423 | SLC2A12 | 78.1 | 94.6 | -0.13 | no |
| Macaque | 772 | STAP1 | 77.4 | 95.7 | -0.13 | no |
| Macaque | 907 | MAP2 | 67.0 | 95.7 | -0.13 | no |
| Cow | 1146 | GUCY1A3 | 70.6 | 95.3 | -0.13 | no |
| Human | 322 | IFI30 | 92.7 | 94.7 | -0.13 | no |
| Human | 201 | RGS1 | 90.3 | 94.7 | -0.13 | no |
| Cow | 1192 | SERPINB2 | 85.4 | 95.3 | -0.13 | no |
| Cow | 1114 | MB21D1 | 23.3 | 95.5 | -0.12 | no |
| Macaque | 746 | WARS | 68.0 | 96.0 | -0.12 | no |
| Macaque | 714 | PMM2 | 52.4 | 96.0 | -0.12 | no |
| Human | 504 | CCDC184 | 83.6 | 95.3 | -0.11 | no |
| Human | 465 | GSDMC | 83.2 | 95.4 | -0.11 | no |
| Macaque | 674 | SAMHD1 | 73.2 | 96.3 | -0.11 | no |
| Human | 479 | TRIM69 | 85.9 | 95.4 | -0.11 | no |
| Human | 385 | HES4 | 93.6 | 95.5 | -0.11 | no |
| Human | 272 | CD274 | 85.4 | 95.6 | -0.11 | no |
| Cow | 1268 | NOSTRIN | 2.9 | 96.2 | -0.10 | no |
| Cow | 1172 | IFI47 | 58.2 | 96.2 | -0.10 | no |
| Cow | 1270 | SNTB1 | 0.5 | 96.2 | -0.10 | no |
| Human | 305 | HELZ2 | 91.3 | 95.8 | -0.10 | no |
| Human | 52 | GPX2 | 89.1 | 96.0 | -0.10 | no |
| Macaque | 847 | NCF1 | 66.8 | 96.9 | -0.10 | no |
| Cow | 1059 | ISG20 | 34.4 | 96.5 | -0.09 | no |
| Cow | 1056 | UBE2L6 | 57.4 | 96.6 | -0.09 | no |
| Human | 79 | UPP2 | 82.2 | 96.2 | -0.09 | no |
| Human | 206 | HEG1 | 87.9 | 96.2 | -0.09 | no |
| Cow | 1186 | PLA2G7 | 68.5 | 96.9 | -0.08 | no |
| Cow | 1033 | PLIN2 | 60.4 | 97.0 | -0.08 | no |
| Macaque | 752 | TRIM5 | 44.3 | 97.4 | -0.08 | no |
| Human | 526 | IL7 | 93.4 | 96.8 | -0.08 | no |
| Cow | 998 | MX1 | 64.2 | 97.1 | -0.08 | no |
| Human | 233 | VEGFC | 80.6 | 96.9 | -0.08 | no |
| Cow | 1081 | GBP4 | 61.4 | 97.3 | -0.07 | no |
| Macaque | 601 | SOCS2 | 79.6 | 97.6 | -0.07 | no |
| Cow | 1176 | MAP1LC3C | 84.5 | 97.4 | -0.07 | no |
| Cow | 1122 | IFNB1 | 84.4 | 97.5 | -0.07 | no |
| Human | 509 | ODF3B | 91.9 | 97.3 | -0.07 | no |
| Macaque | 761 | TAP2 | 30.8 | 97.9 | -0.07 | no |
| Human | 62 | CRP | 86.5 | 97.4 | -0.06 | no |
| Human | 22 | C15orf48 | 91.1 | 97.4 | -0.06 | no |
| Macaque | 627 | MAB21L2 | 79.0 | 98.0 | -0.06 | no |

| | | | | | | |
|---|---|---|---|---|---|---|
| Cow | 1008 | IFI27 | 49.6 | 97.8 | -0.06 | no |
| Cow | 1113 | MB21D1 | 29.8 | 97.8 | -0.06 | no |
| Human | 219 | PNRC1 | 88.9 | 97.7 | -0.06 | no |
| Macaque | 969 | HIST2H2AA3 | 71.8 | 98.2 | -0.05 | no |
| Human | 365 | SLC25A30 | 85.0 | 98.0 | -0.05 | no |
| Macaque | 859 | FLJ23556 | 85.7 | 98.4 | -0.05 | no |
| Cow | 1287 | PLEKHA7 | 7.8 | 98.3 | -0.05 | no |
| Human | 124 | ANGPTL1 | 85.5 | 98.1 | -0.05 | no |
| Cow | 1274 | TLE4 | 54.6 | 98.3 | -0.04 | no |
| Cow | 1257 | GRINA | 73.3 | 98.5 | -0.04 | no |
| Human | 369 | TCF7L2 | 62.6 | 98.3 | -0.04 | no |
| Macaque | 703 | UBE2L6 | 53.2 | 98.8 | -0.04 | no |
| Human | 395 | OGFR | 35.2 | 98.5 | -0.04 | no |
| Macaque | 559 | PLSCR1 | 84.8 | 98.8 | -0.04 | no |
| Human | 145 | CES1 | 73.6 | 98.6 | -0.03 | no |
| Human | 25 | CCL4 | 94.6 | 98.6 | -0.03 | no |
| Macaque | 924 | PSME2 | 70.4 | 99.0 | -0.03 | no |
| Macaque | 572 | CCL5 | 84.9 | 99.0 | -0.03 | no |
| Human | 234 | CNP | 78.4 | 98.8 | -0.03 | no |
| Human | 254 | GBP4 | 79.0 | 98.8 | -0.03 | no |
| Macaque | 902 | TLR1 | 70.3 | 99.1 | -0.03 | no |
| Human | 520 | GNB4 | 86.0 | 99.1 | -0.02 | no |
| Human | 171 | CFB | 64.7 | 99.2 | -0.02 | no |
| Macaque | 605 | CD9 | 62.9 | 99.4 | -0.02 | no |
| Macaque | 608 | GCH1 | 44.8 | 99.4 | -0.02 | no |
| Human | 1935 | TAPBP | 84.9 | 99.2 | -0.02 | no |
| Macaque | 948 | PARP9 | 40.6 | 99.4 | -0.02 | no |
| Human | 334 | ZNF295 | 33.6 | 99.3 | -0.02 | no |
| Cow | 1229 | EPHX2 | 25.8 | 99.4 | -0.02 | no |
| Human | 64 | RNF114 | 89.2 | 99.5 | -0.01 | no |
| Human | 359 | CLEC2B | 84.6 | 99.6 | -0.01 | no |
| Human | 238 | FKBP5 | 44.9 | 99.6 | -0.01 | no |
| Cow | 999 | PLSCR2 | 36.8 | 99.6 | -0.01 | no |
| Macaque | 551 | IFI6 | 57.7 | 99.7 | -0.01 | no |
| Cow | 1108 | STAT2 | 2.5 | 99.7 | -0.01 | no |
| Macaque | 879 | ACSL5 | 17.5 | 99.7 | -0.01 | no |
| Human | 524 | GTPBP1 | 75.4 | 99.7 | -0.01 | no |
| Macaque | 617 | EPSTI1 | 63.4 | 99.9 | 0.00 | no |
| Human | 130 | OASL | 22.3 | 99.9 | 0.00 | no |
| Cow | 1263 | PPP2R3C | 27.8 | 100.0 | 0.00 | no |
| Macaque | 923 | SLC16A4 | 68.3 | 100.0 | 0.00 | no |
| Cow | 1052 | DDX58 | 33.7 | 100.2 | 0.01 | no |
| Human | 161 | PADI2 | 74.6 | 100.3 | 0.01 | no |
| Human | 122 | IFNGR1 | 76.2 | 100.3 | 0.01 | no |
| Cow | 1196 | SP140L | 75.1 | 100.3 | 0.01 | no |
| Macaque | 887 | PRAP1 | 77.4 | 100.4 | 0.01 | no |
| Macaque | 914 | Hist | 78.5 | 100.4 | 0.01 | no |
| Human | 277 | NMI | 84.4 | 100.6 | 0.01 | no |

| | | | | | | |
|---|---|---|---|---|---|---|
| Human | 87 | APOL6 | 61.2 | 100.6 | 0.01 | no |
| Human | 83 | APOL2 | 79.2 | 100.6 | 0.02 | no |
| Human | 271 | RARRES3 | 89.4 | 100.8 | 0.02 | no |
| Cow | 1069 | CNP | 38.7 | 100.7 | 0.02 | no |
| Macaque | 645 | IFIT5 | 56.3 | 100.7 | 0.02 | no |
| Cow | 1223 | CMTR2 | 61.6 | 100.8 | 0.02 | no |
| Human | 350 | NCOA3 | 18.3 | 100.9 | 0.02 | no |
| Cow | 1182 | MTSS1 | 77.6 | 100.8 | 0.02 | no |
| Human | 333 | NPAS2 | 70.9 | 101.0 | 0.02 | no |
| Cow | 1004 | OAS1 | 35.7 | 101.1 | 0.03 | no |
| Human | 138 | CEACAM1 | 0.6 | 101.2 | 0.03 | no |
| Human | 398 | RBM25 | 24.4 | 101.3 | 0.03 | no |
| Macaque | 548 | PSMB8 | 83.4 | 101.1 | 0.03 | no |
| Human | 388 | GBP3 | 80.8 | 101.4 | 0.04 | no |
| Macaque | 700 | DYNLT1 | 77.3 | 101.2 | 0.04 | no |
| Human | 244 | IFIT1 | 67.3 | 101.6 | 0.04 | no |
| Human | 61 | FAM125B | 90.2 | 101.7 | 0.04 | no |
| Macaque | 743 | IFI44 | 52.1 | 101.4 | 0.04 | no |
| Human | 175 | ABTB2 | 77.4 | 101.7 | 0.04 | no |
| Cow | 1015 | LMO2 | 39.8 | 101.6 | 0.04 | no |
| Human | 232 | TMEM49 | 67.1 | 101.9 | 0.05 | no |
| Human | 462 | ASPHD2 | 90.3 | 101.9 | 0.05 | no |
| Cow | 1272 | HECTD2 | 59.0 | 101.8 | 0.05 | no |
| Macaque | 723 | XAF1 | 80.8 | 101.6 | 0.05 | no |
| Human | 226 | GMPR | 76.5 | 102.1 | 0.05 | no |
| Human | 280 | MCL1 | 64.1 | 102.1 | 0.05 | no |
| Cow | 993 | PSMB9 | 79.9 | 101.9 | 0.05 | no |
| Macaque | 698 | ANKFY1 | 45.4 | 101.7 | 0.05 | no |
| Cow | 1039 | GBP2 | 52.4 | 101.9 | 0.05 | no |
| Human | 27 | CXCL10 | 89.4 | 102.2 | 0.05 | no |
| Macaque | 560 | RTP4 | 63.3 | 101.8 | 0.05 | no |
| Human | 394 | RBCK1 | 83.6 | 102.3 | 0.06 | no |
| Human | 368 | CASP1 | 79.8 | 102.3 | 0.06 | no |
| Human | 285 | SERPINB9 | 87.7 | 102.3 | 0.06 | no |
| Macaque | 588 | RNF24 | 73.8 | 101.9 | 0.06 | no |
| Human | 432 | HCP5 | 78.7 | 102.5 | 0.06 | no |
| Macaque | 987 | RGL1 | 67.2 | 102.0 | 0.06 | no |
| Cow | 1143 | CIITA | 1.3 | 102.3 | 0.06 | no |
| Human | 51 | HESX1 | 90.3 | 102.6 | 0.06 | no |
| Macaque | 577 | CCL2 | 81.2 | 102.2 | 0.07 | no |
| Cow | 1098 | TRIM25 | 29.6 | 102.4 | 0.07 | no |
| Human | 198 | ATF3 | 95.9 | 102.7 | 0.07 | no |
| Macaque | 786 | PPM1K | 66.8 | 102.2 | 0.07 | no |
| Human | 213 | IFI35 | 81.3 | 102.8 | 0.07 | no |
| Human | 517 | TEX29 | 81.6 | 102.8 | 0.07 | no |
| Human | 329 | FNDC4 | 90.7 | 102.9 | 0.07 | no |
| Macaque | 693 | MOV10 | 30.0 | 102.3 | 0.07 | no |
| Macaque | 812 | TAGAP | 65.4 | 102.3 | 0.07 | no |

| | | | | | | |
|---|---|---|---|---|---|---|
| Macaque | 968 | ZSCAN12 | 28.1 | 102.3 | 0.07 | no |
| Cow | 1035 | IRF7 | 0.5 | 102.6 | 0.07 | no |
| Macaque | 803 | HLA-X | 68.6 | 102.4 | 0.07 | no |
| Human | 539 | IL22RA1 | 89.0 | 103.0 | 0.07 | no |
| Human | 1912 | BTN3A1 | 81.1 | 103.1 | 0.07 | no |
| Macaque | 607 | LGALS3 | 73.6 | 102.6 | 0.08 | no |
| Human | 399 | RSAD2 | 57.4 | 103.3 | 0.08 | no |
| Macaque | 682 | IFI16 | 59.2 | 102.8 | 0.08 | no |
| Human | 319 | HERC6 | 40.4 | 103.5 | 0.08 | no |
| Macaque | 697 | BUB1 | 22.4 | 102.8 | 0.08 | no |
| Human | 98 | FCGR1A | 74.5 | 103.5 | 0.09 | no |
| Cow | 1303 | CBLN3 | 57.1 | 103.1 | 0.09 | no |
| Macaque | 587 | RNASE4 | 86.3 | 102.9 | 0.09 | no |
| Human | 287 | PSCD1 | 69.7 | 103.7 | 0.09 | no |
| Cow | 1076 | TRIM5 | 22.6 | 103.3 | 0.09 | no |
| Human | 376 | HLA-C | 84.0 | 103.8 | 0.09 | no |
| Macaque | 962 | BTN3A2 | 38.1 | 103.1 | 0.09 | no |
| Cow | 1300 | SCIN | 18.2 | 103.4 | 0.09 | no |
| Macaque | 677 | DHX58 | 36.2 | 103.1 | 0.09 | no |
| Human | 251 | CRY1 | 71.4 | 103.9 | 0.10 | no |
| Human | 31 | GLRX | 78.5 | 104.0 | 0.10 | no |
| Cow | 1011 | B2M | 22.3 | 103.7 | 0.10 | no |
| Macaque | 885 | PLA2G4C | 59.7 | 103.3 | 0.10 | no |
| Human | 428 | KCTD14 | 81.8 | 104.2 | 0.10 | no |
| Human | 475 | CYP2J2 | 93.3 | 104.2 | 0.10 | no |
| Macaque | 935 | FLRT3 | 63.4 | 103.4 | 0.10 | no |
| Human | 347 | C1S | 65.6 | 104.2 | 0.10 | no |
| Macaque | 565 | OAS1 | 42.5 | 103.4 | 0.10 | no |
| Cow | 1290 | GYPC | 56.7 | 103.9 | 0.11 | no |
| Human | 38 | IFITM2 | 93.4 | 104.4 | 0.11 | no |
| Macaque | 552 | GBP1 | 47.0 | 103.5 | 0.11 | no |
| Macaque | 804 | HLA-X | 24.6 | 103.6 | 0.11 | no |
| Cow | 1106 | STAT1 | 38.0 | 104.1 | 0.11 | no |
| Human | 528 | IRF4 | 88.3 | 104.6 | 0.11 | no |
| Cow | 1181 | HLA-ortholog | 11.4 | 104.2 | 0.11 | no |
| Macaque | 846 | APOBEC3A | 71.7 | 103.7 | 0.11 | no |
| Macaque | 564 | IFIT3 | 54.9 | 103.8 | 0.11 | no |
| Macaque | 946 | B3GALNT1 | 57.0 | 103.8 | 0.12 | no |
| Macaque | 769 | RARRES3 | 79.6 | 103.9 | 0.12 | no |
| Macaque | 741 | VEGFC | 55.1 | 103.9 | 0.12 | no |
| Cow | 1261 | DCP2 | 36.5 | 104.4 | 0.12 | no |
| Cow | 1243 | TCAF2 | 28.2 | 104.4 | 0.12 | no |
| Macaque | 840 | CLEC2B | 67.8 | 104.0 | 0.12 | no |
| Macaque | 834 | STAT2 | 65.6 | 104.1 | 0.12 | no |
| Macaque | 658 | GK | 23.9 | 104.1 | 0.13 | no |
| Macaque | 864 | LEPR | 62.1 | 104.2 | 0.13 | no |
| Macaque | 683 | DTX3L | 60.5 | 104.2 | 0.13 | no |
| Human | 1938 | ZNFX1 | 9.6 | 105.3 | 0.13 | no |

| | | | | | | |
|---|---|---|---|---|---|---|
| Cow | 1044 | SP110 | 31.7 | 104.7 | 0.13 | no |
| Macaque | 944 | LYRM2 | 63.9 | 104.3 | 0.13 | no |
| Human | 537 | RBMS1 | 91.0 | 105.4 | 0.13 | no |
| Macaque | 910 | NCOA7 | 49.3 | 104.3 | 0.13 | no |
| Human | 245 | TRIM5 | 70.6 | 105.6 | 0.13 | no |
| Human | 381 | FLJ23556 | 86.2 | 105.6 | 0.14 | no |
| Macaque | 912 | CKB | 77.5 | 104.5 | 0.14 | no |
| Human | 1921 | PLSCR1 | 88.3 | 105.7 | 0.14 | no |
| Cow | 1271 | PARP14 | 2.4 | 105.1 | 0.14 | no |
| Cow | 1051 | TLR3 | 50.5 | 105.2 | 0.14 | no |
| Macaque | 590 | LMO2 | 78.3 | 104.9 | 0.15 | no |
| Macaque | 877 | SPP1 | 66.9 | 105.0 | 0.15 | no |
| Macaque | 919 | TRIM22 | 56.6 | 105.0 | 0.15 | no |
| Human | 205 | CD74 | 91.4 | 106.3 | 0.15 | no |
| Macaque | 736 | USP18 | 70.5 | 105.1 | 0.16 | no |
| Human | 470 | ACTN2 | 50.5 | 106.5 | 0.16 | no |
| Cow | 1003 | IFIT3 | 66.2 | 106.0 | 0.16 | no |
| Human | 468 | CCL7 | 33.7 | 106.7 | 0.16 | no |
| Cow | 1067 | APOL3 | 61.0 | 106.0 | 0.16 | no |
| Cow | 1080 | GBP4 | 52.5 | 106.1 | 0.17 | no |
| Human | 1931 | TRIM5 | 0.2 | 107.1 | 0.17 | no |
| Human | 1940 | XIRP1 | 1.2 | 107.1 | 0.17 | no |
| Human | 204 | DDIT4 | 88.9 | 107.1 | 0.17 | no |
| Cow | 1185 | PIK3CD | 11.9 | 106.4 | 0.17 | no |
| Macaque | 734 | USP18 | 72.4 | 105.8 | 0.18 | no |
| Human | 21 | CMAH | 85.5 | 107.2 | 0.18 | no |
| Human | 519 | HOXD3 | 71.3 | 107.2 | 0.18 | no |
| Macaque | 903 | SQRDL | 68.2 | 105.8 | 0.18 | no |
| Human | 405 | GIMAP2 | 83.0 | 107.3 | 0.18 | no |
| Human | 55 | SOCS2 | 88.7 | 107.4 | 0.18 | no |
| Human | 191 | DYNLT1 | 90.2 | 107.4 | 0.18 | no |
| Human | 447 | GBP7 | 75.4 | 107.4 | 0.18 | no |
| Cow | 1179 | PTCHD3 | 30.3 | 106.7 | 0.18 | no |
| Human | 29 | CCL2 | 91.0 | 107.5 | 0.18 | no |
| Cow | 1296 | DAPP1 | 35.7 | 106.7 | 0.18 | no |
| Macaque | 991 | GMPR2 | 60.4 | 106.2 | 0.19 | no |
| Human | 383 | ARHGAP17 | 53.5 | 107.8 | 0.19 | no |
| Human | 57 | C4orf33 | 93.4 | 107.9 | 0.19 | no |
| Macaque | 615 | FBXO6 | 56.4 | 106.4 | 0.19 | no |
| Cow | 1005 | OAS1 | 86.5 | 107.1 | 0.19 | no |
| Human | 406 | IL18BP | 0.0 | 108.0 | 0.19 | no |
| Cow | 1265 | PAPD4 | 48.5 | 107.1 | 0.19 | no |
| Macaque | 776 | MYD88 | 71.2 | 106.4 | 0.19 | no |
| Human | 312 | C5orf27 | 93.5 | 108.1 | 0.20 | no |
| Human | 209 | SCO2 | 65.9 | 108.2 | 0.20 | no |
| Macaque | 621 | APOL2 | 67.7 | 106.6 | 0.20 | no |
| Cow | 1282 | RHPN2 | 30.1 | 107.4 | 0.20 | no |
| Macaque | 889 | PRAP1 | 67.4 | 106.7 | 0.20 | no |

| | | | | | | |
|---|---|---|---|---|---|---|
| Macaque | 807 | SAMD4A | 59.6 | 106.7 | 0.20 | no |
| Human | 197 | ISG15 | 88.6 | 108.6 | 0.21 | no |
| Human | 119 | PI4K2B | 67.7 | 108.6 | 0.21 | no |
| Macaque | 599 | ANKRD22 | 64.9 | 106.9 | 0.21 | no |
| Human | 1925 | c9orf52 | 67.2 | 109.0 | 0.22 | no |
| Macaque | 728 | HLA-X | 57.0 | 107.2 | 0.22 | no |
| Human | 131 | GTPBP2 | 75.5 | 109.0 | 0.22 | no |
| Human | 425 | HRASLS2 | 75.7 | 109.1 | 0.22 | no |
| Cow | 1202 | TMEM100 | 73.1 | 108.2 | 0.22 | no |
| Human | 295 | OPTN | 79.8 | 109.2 | 0.22 | no |
| Human | 433 | LOC400759 | 79.8 | 109.2 | 0.22 | no |
| Cow | 1030 | IDO1 | 1.5 | 108.3 | 0.22 | no |
| Cow | 1283 | EYA1 | 0.0 | 108.3 | 0.22 | no |
| Macaque | 770 | RARRES3 | 80.3 | 107.4 | 0.23 | no |
| Macaque | 620 | CCDC92 | 62.0 | 107.5 | 0.23 | no |
| Cow | 1285 | NREP | 68.2 | 108.4 | 0.23 | no |
| Human | 1929 | OAS3 | 0.3 | 109.5 | 0.23 | no |
| Macaque | 679 | PARP12 | 35.2 | 107.6 | 0.23 | no |
| Cow | 1079 | GBP4 | 67.8 | 108.6 | 0.23 | no |
| Human | 288 | CLEC4A | 91.3 | 109.7 | 0.23 | no |
| Human | 377 | GAK | 88.0 | 109.7 | 0.24 | no |
| Cow | 1301 | ADGRG6 | 30.0 | 108.7 | 0.24 | no |
| Human | 66 | GZMB | 88.8 | 109.8 | 0.24 | no |
| Cow | 1068 | USP18 | 73.2 | 108.9 | 0.24 | no |
| Human | 507 | THSD1 | 83.7 | 109.9 | 0.24 | no |
| Human | 136 | GK | 50.1 | 109.9 | 0.24 | no |
| Cow | 1017 | BST2 | 14.7 | 109.0 | 0.24 | no |
| Macaque | 961 | FLJ31033 | 3.9 | 108.1 | 0.25 | no |
| Macaque | 806 | HLA-X | 51.9 | 108.2 | 0.25 | no |
| Human | 449 | ZMYND15 | 58.2 | 110.2 | 0.25 | no |
| Cow | 1187 | PLAC8 | 87.0 | 109.2 | 0.25 | no |
| Human | 155 | GBP2 | 81.5 | 110.5 | 0.26 | no |
| Human | 471 | FAM26F | 81.2 | 110.5 | 0.26 | no |
| Cow | 1124 | PARP10 | 0.3 | 109.5 | 0.26 | no |
| Macaque | 817 | ADM | 77.4 | 108.6 | 0.26 | no |
| Macaque | 626 | ARG2 | 73.0 | 108.7 | 0.26 | no |
| Macaque | 673 | GBP2 | 20.3 | 108.7 | 0.26 | no |
| Cow | 1136 | MORC3 | 29.2 | 109.7 | 0.27 | no |
| Human | 542 | KLHDC7B | 69.8 | 110.9 | 0.27 | no |
| Macaque | 554 | GBP1 | 46.7 | 108.9 | 0.27 | no |
| Cow | 1089 | CD274 | 57.2 | 110.0 | 0.27 | no |
| Macaque | 802 | CCNA1 | 48.7 | 109.0 | 0.27 | no |
| Human | 459 | ARL9 | 85.4 | 111.5 | 0.28 | no |
| Macaque | 908 | C3AR1 | 57.2 | 109.3 | 0.28 | no |
| Cow | 1144 | KIF5C | 25.1 | 110.3 | 0.28 | no |
| Cow | 1232 | VWA5A | 55.2 | 110.4 | 0.28 | no |
| Macaque | 805 | HLA-X | 61.7 | 109.4 | 0.28 | no |
| Human | 94 | CCR1 | 78.0 | 111.7 | 0.29 | no |

| | | | | | | |
|---|---|---|---|---|---|---|
| Human | 257 | TAP2 | 31.6 | 111.8 | 0.29 | no |
| Human | 135 | PCTK2 | 84.6 | 111.8 | 0.29 | no |
| Macaque | 687 | STAT3 | 12.2 | 109.6 | 0.29 | no |
| Cow | 1032 | ZBP1 | 3.1 | 110.7 | 0.29 | no |
| Human | 73 | TNFSF13B | 84.7 | 112.1 | 0.29 | no |
| Human | 438 | CD47 | 67.9 | 112.1 | 0.29 | no |
| Human | 9 | PLSCR2 | 91.8 | 112.2 | 0.30 | no |
| Human | 114 | NUP50 | 73.9 | 112.2 | 0.30 | no |
| Macaque | 915 | PANX1 | 62.7 | 109.8 | 0.30 | no |
| Cow | 1121 | TAPBPL | 64.0 | 111.0 | 0.30 | no |
| Human | 262 | TAP1 | 49.9 | 112.3 | 0.30 | no |
| Macaque | 824 | NT5C3 | 56.6 | 109.9 | 0.30 | no |
| Macaque | 855 | ADAMDEC1 | 70.9 | 110.0 | 0.30 | no |
| Human | 408 | TAPBPL | 0.0 | 112.5 | 0.30 | no |
| Macaque | 884 | ENPP2 | 68.1 | 110.0 | 0.30 | no |
| Macaque | 856 | ADAMDEC1 | 70.1 | 110.1 | 0.31 | no |
| Macaque | 709 | RGS1 | 48.2 | 110.1 | 0.31 | no |
| Cow | 1026 | C19orf66 | 8.6 | 111.4 | 0.31 | no |
| Human | 10 | RTP4 | 74.1 | 112.8 | 0.31 | no |
| Human | 13 | CCDC75 | 83.0 | 112.8 | 0.31 | no |
| Human | 34 | SAA1 | 91.9 | 112.8 | 0.31 | no |
| Human | 2 | PSMB9 | 89.6 | 112.8 | 0.31 | no |
| Human | 310 | HLA-F | 84.1 | 112.9 | 0.31 | no |
| Human | 540 | GLYATL1 | 75.2 | 112.9 | 0.31 | no |
| Macaque | 616 | GEM | 79.2 | 110.3 | 0.31 | no |
| Human | 263 | CMTR1 | 82.6 | 113.0 | 0.32 | no |
| Human | 343 | FAM46A | 63.3 | 113.1 | 0.32 | no |
| Human | 492 | SLC6A9 | 69.0 | 113.2 | 0.32 | no |
| Macaque | 876 | PCGF5 | 32.0 | 110.6 | 0.32 | no |
| Human | 442 | NUB1 | 76.3 | 113.4 | 0.33 | no |
| Macaque | 702 | LY6E | 70.4 | 110.8 | 0.33 | no |
| Cow | 1305 | GBP6 | 58.2 | 112.1 | 0.33 | no |
| Cow | 1127 | SP100 | 55.6 | 112.2 | 0.33 | no |
| Human | 160 | CTCFL | 68.7 | 113.7 | 0.33 | no |
| Human | 144 | EIF3EIP | 85.9 | 113.9 | 0.34 | no |
| Cow | 1130 | APOBEC3B | 4.0 | 112.5 | 0.34 | no |
| Human | 464 | SLITRK2 | 84.4 | 114.0 | 0.34 | no |
| Macaque | 722 | XAF1 | 80.5 | 111.3 | 0.34 | no |
| Cow | 1009 | C15orf48 | 3.5 | 112.7 | 0.35 | no |
| Human | 413 | APOBEC3D | 80.8 | 114.3 | 0.35 | no |
| Macaque | 619 | BCL2L14 | 71.1 | 111.6 | 0.35 | no |
| Macaque | 644 | PI4K2B | 42.6 | 111.6 | 0.35 | no |
| Human | 35 | IFITM1 | 70.1 | 114.6 | 0.36 | no |
| Human | 543 | RUFY4 | 62.7 | 114.7 | 0.36 | no |
| Human | 494 | HLA-DOB | 74.2 | 114.8 | 0.36 | no |
| Macaque | 754 | TRIMCYP | 64.5 | 111.9 | 0.36 | no |
| Human | 1918 | IFITM1 | 88.4 | 115.1 | 0.37 | no |
| Macaque | 717 | SPSB1 | 62.4 | 112.1 | 0.37 | no |

| | | | | | | |
|---|---|---|---|---|---|---|
| Macaque | 583 | RPL22 | 54.2 | 112.2 | 0.37 | no |
| Macaque | 606 | RNF114 | 79.8 | 112.3 | 0.37 | no |
| Cow | 1293 | TM4SF1 | 58.9 | 113.7 | 0.37 | no |
| Macaque | 955 | THEMIS2 | 70.1 | 112.4 | 0.38 | no |
| Macaque | 868 | ADAR | 57.8 | 112.4 | 0.38 | no |
| Macaque | 866 | ADAR | 22.6 | 112.5 | 0.38 | no |
| Macaque | 947 | IL4I1 | 66.6 | 112.5 | 0.38 | no |
| Macaque | 793 | TRAFD1 | 46.2 | 112.5 | 0.38 | no |
| Macaque | 841 | IL15 | 71.5 | 112.5 | 0.38 | no |
| Human | 512 | HAVCR2 | 93.8 | 115.7 | 0.38 | no |
| Human | 14 | IFIT3 | 75.8 | 115.8 | 0.38 | no |
| Macaque | 614 | TNFSF13B | 60.6 | 112.7 | 0.39 | no |
| Human | 267 | KIAA1618 | 59.8 | 115.9 | 0.39 | no |
| Macaque | 676 | NDC80 | 32.3 | 113.0 | 0.39 | no |
| Human | 346 | ABLIM3 | 92.4 | 116.4 | 0.40 | no |
| Cow | 1112 | HES4 | 61.4 | 114.7 | 0.40 | no |
| Macaque | 558 | MX1 | 67.4 | 113.2 | 0.40 | no |
| Human | 473 | TSPAN33 | 92.2 | 116.7 | 0.41 | no |
| Human | 500 | TMEM106A | 85.5 | 116.7 | 0.41 | no |
| Cow | 1194 | SP140L | 45.7 | 115.0 | 0.41 | no |
| Human | 357 | IL6ST | 68.5 | 116.8 | 0.41 | no |
| Human | 407 | C3AR1 | 82.9 | 116.9 | 0.41 | no |
| Cow | 1134 | AZI2 | 1.5 | 115.1 | 0.41 | no |
| Cow | 1145 | CYP2J2 | 12.8 | 115.1 | 0.41 | no |
| Cow | 1231 | VWA5A | 76.9 | 115.3 | 0.42 | no |
| Cow | 1037 | NAMPT | 63.1 | 115.3 | 0.42 | no |
| Human | 452 | GRIP2 | 39.0 | 117.1 | 0.42 | no |
| Cow | 1152 | IRF4 | 0.0 | 115.4 | 0.42 | no |
| Human | 373 | JUNB | 85.9 | 117.3 | 0.42 | no |
| Cow | 1240 | SDS | 0.0 | 115.5 | 0.42 | no |
| Cow | 1110 | IL15 | 1.2 | 115.5 | 0.42 | no |
| Cow | 1298 | TAF3 | 2.7 | 115.5 | 0.42 | no |
| Cow | 1288 | BMK | 37.7 | 115.6 | 0.43 | no |
| Human | 153 | GBP5 | 28.1 | 117.6 | 0.43 | no |
| Macaque | 862 | MB21D1 | 39.5 | 114.1 | 0.43 | no |
| Macaque | 777 | NMI | 55.7 | 114.2 | 0.43 | no |
| Human | 1920 | OAS2 | 10.5 | 117.8 | 0.43 | no |
| Macaque | 823 | ZNF295 | 22.0 | 114.3 | 0.43 | no |
| Human | 522 | PHACTR4 | 77.8 | 117.9 | 0.43 | no |
| Human | 97 | CSDA | 77.7 | 118.0 | 0.44 | no |
| Human | 404 | CLDN23 | 81.2 | 118.1 | 0.44 | no |
| Cow | 1061 | IFI35 | 55.3 | 116.4 | 0.45 | no |
| Human | 230 | TXNIP | 41.0 | 118.5 | 0.45 | no |
| Human | 538 | WASHC4 | 45.3 | 118.7 | 0.46 | no |
| Human | 185 | BUB1 | 61.1 | 118.8 | 0.46 | no |
| Cow | 1165 | DRAM1 | 61.3 | 116.8 | 0.46 | no |
| Macaque | 787 | PPM1K | 58.6 | 115.1 | 0.46 | no |
| Human | 389 | IL1R | 76.8 | 119.0 | 0.46 | no |

| | | | | | | |
|---|---|---|---|---|---|---|
| Macaque | 598 | PHF11 | 76.5 | 115.3 | 0.46 | no |
| Macaque | 718 | IMPA2 | 50.6 | 115.3 | 0.46 | no |
| Cow | 1159 | ATXN3 | 44.1 | 117.2 | 0.47 | no |
| Macaque | 956 | CA5B | 78.8 | 115.4 | 0.47 | no |
| Macaque | 597 | GPX2 | 47.4 | 115.5 | 0.47 | no |
| Human | 499 | TMEM106A | 91.2 | 119.4 | 0.47 | no |
| Human | 296 | SLC15A3 | 84.1 | 119.4 | 0.47 | no |
| Cow | 1049 | ABTB2 | 5.9 | 117.5 | 0.48 | no |
| Macaque | 719 | IFI35 | 69.8 | 115.7 | 0.48 | no |
| Macaque | 897 | LGALS3BP | 74.8 | 115.7 | 0.48 | no |
| Cow | 1082 | GBP4 | 55.2 | 117.7 | 0.48 | no |
| Human | 448 | IFIT1B | 61.9 | 119.9 | 0.48 | no |
| Human | 466 | STX11 | 89.7 | 119.9 | 0.48 | no |
| Macaque | 788 | PPM1K | 60.8 | 116.0 | 0.48 | no |
| Cow | 1096 | SLC15A3 | 59.0 | 117.9 | 0.49 | no |
| Cow | 1085 | PNPT1 | 0.1 | 118.1 | 0.49 | no |
| Human | 386 | AHNAK2 | 56.3 | 120.3 | 0.49 | no |
| Macaque | 763 | PNPT1 | 21.4 | 116.4 | 0.50 | no |
| Human | 451 | EXOC3L1 | 49.5 | 120.4 | 0.50 | no |
| Human | 137 | SLC1A1 | 65.9 | 120.5 | 0.50 | no |
| Macaque | 747 | TRIM21 | 54.4 | 116.4 | 0.50 | no |
| Macaque | 905 | PSME1 | 63.4 | 116.4 | 0.50 | no |
| Cow | 1189 | PTPRE | 51.7 | 118.4 | 0.50 | no |
| Human | 224 | HLA-G | 72.8 | 120.6 | 0.50 | no |
| Human | 176 | EPAS1 | 63.9 | 120.7 | 0.50 | no |
| Human | 410 | TRIM22 | 69.8 | 120.7 | 0.50 | no |
| Human | 1923 | RSAD2 | 45.6 | 120.8 | 0.51 | no |
| Human | 495 | TESK2 | 63.3 | 120.8 | 0.51 | no |
| Human | 481 | RASGRP3 | 76.7 | 120.8 | 0.51 | no |
| Macaque | 895 | LGALS3BP | 66.1 | 116.7 | 0.51 | no |
| Macaque | 774 | NAPA | 27.0 | 116.8 | 0.51 | no |
| Cow | 1111 | IFIT2 | 65.4 | 118.7 | 0.51 | no |
| Human | 12 | EIF2AK2 | 41.5 | 121.0 | 0.51 | no |
| Human | 80 | MAFB | 93.4 | 121.1 | 0.51 | no |
| Cow | 1048 | PHF15 | 0.3 | 118.9 | 0.51 | no |
| Human | 90 | FAM70A | 90.4 | 121.1 | 0.51 | no |
| Macaque | 800 | IRF2 | 2.7 | 117.2 | 0.52 | no |
| Macaque | 610 | TNFAIP6 | 57.1 | 117.2 | 0.52 | no |
| Macaque | 705 | ISG15 | 66.3 | 117.2 | 0.52 | no |
| Human | 228 | PDGFRL | 81.3 | 121.5 | 0.52 | no |
| Cow | 1088 | TDRD7 | 29.5 | 119.4 | 0.53 | no |
| Human | 56 | CD69 | 84.9 | 121.7 | 0.53 | no |
| Macaque | 580 | VAMP5 | 59.8 | 117.4 | 0.53 | no |
| Cow | 1129 | HERC5 | 12.4 | 119.5 | 0.53 | no |
| Cow | 1160 | BOLA-DMB | 81.2 | 119.8 | 0.54 | no |
| Cow | 1199 | TGM1 | 43.6 | 119.9 | 0.54 | no |
| Cow | 1131 | RBM43 | 29.6 | 120.0 | 0.54 | no |
| Macaque | 706 | ATF3 | 89.3 | 118.0 | 0.55 | no |

| | | | | | | |
|---|---|---|---|---|---|---|
| Macaque | 550 | IFI6 | 46.4 | 118.0 | 0.55 | no |
| Human | 126 | ALDH1A1 | 64.8 | 122.5 | 0.55 | no |
| Human | 458 | PIK3AP1 | 75.1 | 122.5 | 0.55 | no |
| Macaque | 667 | PXK | 44.5 | 118.1 | 0.55 | no |
| Macaque | 865 | OGFR | 10.7 | 118.1 | 0.55 | no |
| Macaque | 875 | PCGF5 | 59.2 | 118.2 | 0.55 | no |
| Human | 477 | KCNT2 | 30.1 | 122.7 | 0.55 | no |
| Cow | 1280 | CUL4B | 0.8 | 120.3 | 0.55 | no |
| Macaque | 629 | FAM46C | 34.1 | 118.4 | 0.56 | no |
| Macaque | 797 | ERLIN1 | 1.8 | 118.5 | 0.56 | no |
| Human | 352 | PRKD2 | 72.9 | 123.1 | 0.56 | no |
| Cow | 1304 | CXCR4 | 74.1 | 120.8 | 0.57 | no |
| Macaque | 553 | GBP1 | 47.3 | 118.7 | 0.57 | no |
| Macaque | 628 | APOBEC3G | 8.7 | 118.7 | 0.57 | no |
| Human | 261 | PNPT1 | 63.8 | 123.4 | 0.57 | no |
| Human | 422 | THEMIS2 | 67.4 | 123.5 | 0.57 | no |
| Macaque | 696 | BUB1 | 38.6 | 118.9 | 0.57 | no |
| Human | 338 | BAG1 | 64.4 | 123.8 | 0.58 | no |
| Human | 166 | IFI16 | 71.7 | 123.8 | 0.58 | no |
| Cow | 1058 | ATF3 | 51.4 | 121.3 | 0.58 | no |
| Macaque | 973 | APOBEC3B | 43.4 | 119.1 | 0.58 | no |
| Human | 325 | CX3CL1 | 71.1 | 123.9 | 0.58 | no |
| Cow | 1170 | HAS3 | 60.2 | 121.8 | 0.59 | no |
| Macaque | 680 | IFI16 | 0.4 | 119.6 | 0.59 | no |
| Human | 156 | BIRC3 | 70.8 | 124.5 | 0.59 | no |
| Human | 461 | LRRTM2 | 0.0 | 124.6 | 0.60 | no |
| Human | 146 | SQLE | 88.0 | 124.8 | 0.60 | no |
| Human | 445 | CMPK2 | 81.3 | 125.1 | 0.61 | no |
| Cow | 1101 | HERC6 | 37.2 | 122.6 | 0.62 | no |
| Macaque | 990 | PLSCR2 | 59.3 | 120.3 | 0.62 | no |
| Cow | 994 | PSMB8 | 11.3 | 122.7 | 0.62 | no |
| Cow | 1236 | GGT6 | 80.0 | 122.7 | 0.62 | no |
| Cow | 1053 | MOV10 | 0.0 | 122.7 | 0.62 | no |
| Macaque | 557 | IFITM3 | 87.5 | 120.4 | 0.62 | no |
| Human | 431 | HSPA6 | 45.6 | 125.8 | 0.63 | no |
| Human | 402 | PCGF5 | 19.2 | 126.0 | 0.63 | no |
| Human | 123 | NAMPT | 74.0 | 126.1 | 0.63 | no |
| Macaque | 831 | C1S | 49.6 | 121.0 | 0.64 | no |
| Macaque | 664 | PAK3 | 31.3 | 121.0 | 0.64 | no |
| Macaque | 883 | ENPP2 | 68.5 | 121.1 | 0.64 | no |
| Macaque | 652 | C22orf28 | 27.6 | 121.2 | 0.64 | no |
| Macaque | 602 | CD69 | 33.3 | 121.2 | 0.64 | no |
| Cow | 1070 | IFI44 | 63.0 | 123.7 | 0.65 | no |
| Cow | 1198 | TBX21 | 0.0 | 123.8 | 0.65 | no |
| Human | 106 | IL17RB | 92.5 | 126.8 | 0.65 | no |
| Human | 315 | ZNF385B | 88.6 | 126.9 | 0.65 | no |
| Human | 1913 | BTN3A2 | 14.6 | 126.9 | 0.65 | no |
| Macaque | 861 | MB21D1 | 39.8 | 121.6 | 0.65 | no |

| | | | | | | |
|---|---|---|---|---|---|---|
| Macaque | 646 | TYMP | 35.9 | 121.7 | 0.66 | no |
| Cow | 1221 | FLVCR2 | 70.2 | 124.2 | 0.66 | no |
| Cow | 1200 | TGM2 | 0.7 | 124.4 | 0.66 | no |
| Human | 440 | AZI2 | 88.8 | 127.3 | 0.66 | no |
| Human | 371 | SLC16A1 | 76.0 | 127.5 | 0.67 | no |
| Macaque | 911 | NCOA7 | 42.6 | 122.0 | 0.67 | no |
| Macaque | 790 | LGMN | 28.3 | 122.1 | 0.67 | no |
| Macaque | 669 | DCP1A | 30.1 | 122.2 | 0.67 | no |
| Human | 227 | USP18 | 82.7 | 127.7 | 0.67 | no |
| Cow | 1162 | CASP4 | 59.2 | 124.8 | 0.68 | no |
| Human | 510 | HAPLN3 | 70.7 | 127.9 | 0.68 | no |
| Cow | 1302 | PARP8 | 8.7 | 124.9 | 0.68 | no |
| Macaque | 975 | KCTD14 | 38.6 | 122.5 | 0.68 | no |
| Cow | 1072 | WARS | 17.7 | 125.1 | 0.68 | no |
| Cow | 1284 | LGI2 | 66.2 | 125.2 | 0.68 | no |
| Human | 514 | HDX | 59.4 | 128.2 | 0.69 | no |
| Macaque | 779 | GJA4 | 79.9 | 122.9 | 0.69 | no |
| Human | 75 | C19orf66 | 51.1 | 128.7 | 0.70 | no |
| Human | 1943 | CD7 | 80.3 | 128.7 | 0.70 | no |
| Macaque | 556 | IFITM3 | 81.8 | 123.0 | 0.70 | no |
| Macaque | 901 | FOXN2 | 62.9 | 123.1 | 0.70 | no |
| Macaque | 712 | RAB27A | 13.2 | 123.1 | 0.70 | no |
| Macaque | 832 | C1S | 48.5 | 123.1 | 0.70 | no |
| Macaque | 634 | SMAD3 | 84.3 | 123.3 | 0.71 | no |
| Macaque | 827 | FAM46A | 54.4 | 123.5 | 0.71 | no |
| Cow | 1212 | GPR63 | 1.3 | 126.3 | 0.72 | no |
| Human | 132 | LAP3 | 68.0 | 129.6 | 0.72 | no |
| Human | 421 | PARP9 | 57.0 | 129.6 | 0.72 | no |
| Human | 1915 | IFI6 | 10.8 | 129.7 | 0.72 | no |
| Human | 208 | SECTM1 | 92.7 | 129.7 | 0.72 | no |
| Macaque | 694 | MOV10 | 0.1 | 123.9 | 0.72 | no |
| Human | 506 | AGBL2 | 44.6 | 130.0 | 0.73 | no |
| Cow | 1027 | EPSTI1 | 26.4 | 126.9 | 0.73 | no |
| Human | 378 | ADAMDEC1 | 78.6 | 130.3 | 0.74 | no |
| Cow | 1139 | USP25 | 1.4 | 127.2 | 0.74 | no |
| Macaque | 603 | C4orf33 | 58.9 | 124.4 | 0.74 | no |
| Macaque | 751 | TRIM5 | 34.2 | 124.5 | 0.74 | no |
| Human | 1937 | RICTOR | 1.8 | 130.7 | 0.75 | no |
| Macaque | 844 | PTMA | 4.0 | 124.7 | 0.75 | no |
| Cow | 1024 | TNFSF13B | 62.2 | 127.6 | 0.75 | no |
| Macaque | 849 | MAX | 13.9 | 124.8 | 0.75 | no |
| Human | 225 | AIM2 | 77.4 | 131.0 | 0.75 | no |
| Macaque | 749 | IFIT1 | 54.9 | 124.9 | 0.76 | no |
| Human | 429 | SIGLEC1 | 86.6 | 131.3 | 0.76 | no |
| Cow | 1180 | MCHR1 | 41.5 | 128.0 | 0.76 | no |
| Macaque | 848 | CASP1 | 10.3 | 125.4 | 0.77 | no |
| Human | 536 | AIDA | 79.2 | 131.7 | 0.77 | no |
| Human | 345 | TRIM34 | 69.9 | 132.1 | 0.78 | no |

| | | | | | | |
|---|---|---|---|---|---|---|
| Cow | 1031 | IFI44L | 13.1 | 128.7 | 0.78 | no |
| Macaque | 854 | ADAMDEC1 | 83.1 | 125.8 | 0.78 | no |
| Macaque | 838 | IL6ST | 50.9 | 126.0 | 0.79 | no |
| Human | 482 | DLL1 | 73.6 | 132.5 | 0.79 | no |
| Macaque | 894 | C17orf60 | 76.7 | 126.2 | 0.79 | no |
| Macaque | 757 | ARNTL | 37.9 | 126.2 | 0.80 | no |
| Human | 420 | IL4I1 | 84.2 | 132.8 | 0.80 | no |
| Macaque | 686 | CFB | 35.0 | 126.4 | 0.80 | no |
| Human | 92 | MKX | 42.6 | 133.1 | 0.80 | no |
| Human | 434 | RBM43 | 54.5 | 133.2 | 0.81 | no |
| Human | 37 | RPL22 | 80.2 | 133.3 | 0.81 | no |
| Human | 456 | KIF5C | 16.9 | 133.6 | 0.82 | no |
| Human | 186 | NOS2A | 0.6 | 133.7 | 0.82 | no |
| Human | 265 | MASTL | 57.1 | 133.9 | 0.82 | no |
| Macaque | 655 | LAP3 | 55.2 | 127.2 | 0.83 | no |
| Human | 1927 | NCOA7 | 69.1 | 134.1 | 0.83 | no |
| Human | 177 | PML | 60.1 | 134.2 | 0.83 | no |
| Macaque | 808 | ZNF385B | 83.1 | 127.4 | 0.83 | no |
| Human | 415 | RXFP1 | 58.6 | 134.4 | 0.84 | no |
| Macaque | 784 | SERPINB9 | 20.2 | 127.5 | 0.84 | no |
| Cow | 1166 | FBXO16 | 21.4 | 130.8 | 0.84 | no |
| Human | 396 | ADAR | 80.9 | 134.6 | 0.84 | no |
| Human | 457 | FAM107A | 17.5 | 134.6 | 0.84 | no |
| Macaque | 748 | IFIT1 | 46.3 | 127.8 | 0.84 | no |
| Human | 541 | TLDC2 | 85.6 | 134.7 | 0.84 | no |
| Macaque | 731 | AIM2 | 15.5 | 128.9 | 0.88 | no |
| Macaque | 651 | ALDH1A1 | 29.1 | 129.2 | 0.89 | no |
| Human | 240 | TRIM38 | 87.2 | 136.7 | 0.89 | no |
| Macaque | 647 | IFNGR1 | 32.6 | 129.8 | 0.90 | no |
| Macaque | 985 | AXL | 59.6 | 129.9 | 0.91 | no |
| Macaque | 604 | FAM125B | 62.1 | 130.0 | 0.91 | no |
| Macaque | 980 | PTAR1 | 46.9 | 130.3 | 0.92 | no |
| Cow | 1013 | IFITM3 | 75.3 | 133.8 | 0.92 | no |
| Human | 1939 | FMR1 | 0.2 | 138.0 | 0.92 | no |
| Cow | 1239 | SH2D3C | 43.7 | 134.1 | 0.93 | no |
| Cow | 1066 | APOL3 | 17.4 | 134.2 | 0.93 | no |
| Cow | 1045 | IFI16 | 53.1 | 134.3 | 0.93 | no |
| Macaque | 671 | BTN3A3 | 30.4 | 131.0 | 0.94 | no |
| Cow | 1250 | TLR4 | 80.0 | 134.6 | 0.94 | no |
| Macaque | 567 | CHMP5 | 69.0 | 131.0 | 0.94 | no |
| Human | 289 | PPM1K | 88.9 | 138.9 | 0.94 | no |
| Human | 502 | P2RX7 | 87.5 | 138.9 | 0.94 | no |
| Human | 355 | DDX60 | 7.2 | 139.0 | 0.95 | no |
| Human | 437 | TNK2 | 32.4 | 139.0 | 0.95 | no |
| Human | 400 | ZC3HAV1 | 55.7 | 139.0 | 0.95 | no |
| Macaque | 762 | MX2 | 23.8 | 131.3 | 0.95 | no |
| Human | 523 | CPEB3 | 34.0 | 139.1 | 0.95 | no |
| Human | 478 | ANKRD45 | 70.5 | 139.3 | 0.95 | no |

| | | | | | | |
|---|---|---|---|---|---|---|
| Human | 480 | APOL4 | 17.4 | 139.3 | 0.96 | no |
| Cow | 1060 | CD74 | 53.1 | 135.4 | 0.96 | no |
| Human | 463 | FYB | 42.7 | 139.6 | 0.96 | no |
| Human | 476 | BRIP1 | 44.3 | 139.7 | 0.96 | no |
| Macaque | 921 | NPBWR1 | 67.6 | 132.1 | 0.97 | no |
| Macaque | 742 | CNP | 36.2 | 132.1 | 0.98 | no |
| Macaque | 759 | PABPC4 | 24.9 | 132.1 | 0.98 | no |
| Cow | 1071 | TRIM38 | 71.8 | 135.9 | 0.98 | no |
| Human | 252 | ELF1 | 96.1 | 140.3 | 0.98 | no |
| Human | 276 | MYD88 | 95.6 | 140.3 | 0.98 | no |
| Cow | 1197 | TACSTD2 | 73.0 | 136.0 | 0.98 | no |
| Macaque | 555 | IFITM3 | 84.8 | 132.4 | 0.98 | no |
| Human | 159 | AMPH | 66.1 | 140.7 | 0.99 | no |
| Human | 58 | MS4A4A | 85.9 | 140.7 | 0.99 | no |
| Macaque | 670 | BTN3A3 | 25.8 | 132.9 | 1.00 | no |
| Human | 148 | PXK | 91.7 | 141.3 | 1.00 | no |
| Human | 243 | TRIM21 | 58.0 | 141.3 | 1.00 | no |
| Macaque | 740 | TMEM49 | 5.2 | 133.5 | 1.02 | no |
| Human | 253 | ARNTL | 75.1 | 142.4 | 1.03 | no |
| Macaque | 881 | ACSL5 | 19.4 | 134.0 | 1.03 | no |
| Human | 472 | TMEM171 | 91.8 | 142.6 | 1.04 | no |
| Macaque | 871 | ZC3HAV1 | 61.4 | 134.2 | 1.04 | no |
| Macaque | 648 | NAMPT | 34.2 | 134.2 | 1.04 | no |
| Macaque | 609 | TMEM51 | 63.8 | 134.4 | 1.04 | no |
| Human | 430 | XRN | 4.2 | 143.0 | 1.04 | no |
| Cow | 1286 | CGNL1 | 2.0 | 138.5 | 1.05 | no |
| Cow | 1141 | PCDH17 | 48.9 | 138.7 | 1.05 | no |
| Macaque | 853 | IFIT2 | 12.4 | 134.8 | 1.06 | no |
| Macaque | 945 | ERAP2 | 29.5 | 135.0 | 1.06 | no |
| Cow | 1115 | OGFR | 2.5 | 139.0 | 1.06 | no |
| Cow | 1116 | ADAR | 0.6 | 139.7 | 1.08 | no |
| Macaque | 691 | TLR3 | 63.8 | 135.7 | 1.08 | no |
| Macaque | 574 | CXCL11 | 1.6 | 135.7 | 1.08 | no |
| Macaque | 880 | ACSL5 | 69.9 | 135.7 | 1.08 | no |
| Macaque | 750 | TRIM5 | 38.5 | 135.7 | 1.08 | no |
| Macaque | 983 | XRN | 4.1 | 135.9 | 1.09 | no |
| Macaque | 785 | PSCD1 | 31.3 | 136.0 | 1.09 | no |
| Cow | 1036 | IFIT5 | 0.0 | 140.2 | 1.09 | no |
| Macaque | 938 | PLA2G4A | 38.2 | 136.1 | 1.10 | no |
| Human | 237 | MSR1 | 74.1 | 145.2 | 1.10 | no |
| Macaque | 775 | BLVRA | 20.3 | 136.2 | 1.10 | no |
| Human | 1919 | Mx1 | 2.3 | 145.4 | 1.10 | no |
| Human | 141 | HPSE | 89.9 | 145.6 | 1.11 | no |
| Macaque | 756 | ELF1 | 92.8 | 136.8 | 1.12 | no |
| Human | 498 | TLR2 | 75.5 | 145.9 | 1.12 | no |
| Macaque | 758 | PABPC4 | 19.8 | 136.9 | 1.12 | no |
| Macaque | 745 | TRIM38 | 60.4 | 136.9 | 1.12 | no |
| Macaque | 730 | AIM2 | 53.7 | 137.5 | 1.14 | no |

| | | | | | | |
|---|---|---|---|---|---|---|
| Cow | 1177 | MYADM | 0.2 | 142.5 | 1.16 | no |
| Macaque | 792 | OPTN | 16.7 | 138.3 | 1.16 | no |
| Macaque | 755 | TRIM5 | 35.6 | 138.3 | 1.16 | no |
| Human | 444 | USP25 | 40.2 | 148.3 | 1.17 | no |
| Human | 17 | CHMP5 | 68.0 | 148.7 | 1.18 | no |
| Human | 490 | SLFN13 | 33.1 | 148.8 | 1.19 | no |
| Macaque | 929 | APOBEC3F | 4.8 | 139.1 | 1.19 | no |
| Human | 497 | PIK3R3 | 48.1 | 148.9 | 1.19 | no |
| Human | 91 | HSH2D | 90.2 | 149.8 | 1.21 | no |
| Human | 212 | IMPA2 | 74.2 | 150.2 | 1.22 | no |
| Human | 435 | RGL-1 | 78.5 | 150.2 | 1.22 | no |
| Cow | 1178 | LOC512672 | 10.8 | 145.6 | 1.24 | no |
| Human | 320 | NOD2 | 78.5 | 151.1 | 1.24 | no |
| Human | 259 | ZNF107 | 0.3 | 151.2 | 1.24 | no |
| Human | 487 | SLAMF8 | 85.4 | 151.8 | 1.26 | no |
| Human | 532 | ZCCHC2 | 25.4 | 151.8 | 1.26 | no |
| Macaque | 977 | NBN | 29.7 | 141.5 | 1.26 | no |
| Human | 109 | PDK1 | 85.4 | 152.1 | 1.27 | no |
| Cow | 1188 | PREX2 | 48.8 | 146.7 | 1.27 | no |
| Human | 450 | PCDH17 | 62.9 | 152.5 | 1.28 | no |
| Cow | 1161 | CA9 | 12.6 | 146.9 | 1.28 | no |
| Human | 1941 | PARP14 | 3.5 | 153.2 | 1.29 | no |
| Human | 255 | PABPC4 | 42.3 | 154.0 | 1.31 | no |
| Human | 454 | CIITA | 0.4 | 154.3 | 1.32 | no |
| Macaque | 659 | SLC1A1 | 47.2 | 143.5 | 1.32 | no |
| Human | 39 | C4orf32 | 80.8 | 154.4 | 1.32 | no |
| Macaque | 809 | ZNF385B | 76.2 | 143.7 | 1.33 | no |
| Human | 535 | STOML1 | 85.0 | 154.8 | 1.33 | no |
| Cow | 1248 | PLEKHN1 | 59.1 | 149.0 | 1.33 | no |
| Macaque | 713 | CD74 | 67.8 | 144.0 | 1.33 | no |
| Macaque | 936 | MLH3 | 0.5 | 144.0 | 1.34 | no |
| Macaque | 595 | TMEM140 | 77.3 | 144.2 | 1.34 | no |
| Macaque | 816 | IFI30 | 76.5 | 144.6 | 1.35 | no |
| Cow | 1128 | SP100 | 50.7 | 150.9 | 1.39 | no |
| Cow | 1167 | FBXO33 | 13.0 | 151.3 | 1.40 | no |
| Macaque | 942 | GJD3 | 33.1 | 146.2 | 1.40 | no |
| Macaque | 852 | JUNB | 55.0 | 146.3 | 1.41 | no |
| Cow | 1093 | SLC25A28 | 29.0 | 152.3 | 1.42 | no |
| Human | 249 | RNF19B | 72.9 | 158.7 | 1.43 | no |
| Human | 18 | PMAIP1 | 76.2 | 159.0 | 1.43 | no |
| Macaque | 810 | UNC93B1 | 62.1 | 147.4 | 1.44 | no |
| Macaque | 771 | CD274 | 6.7 | 147.7 | 1.45 | no |
| Human | 178 | TLR3 | 85.6 | 159.7 | 1.45 | no |
| Cow | 1190 | RAB8B | 47.0 | 153.6 | 1.46 | no |
| Macaque | 927 | APOBEC3F | 3.7 | 148.1 | 1.46 | no |
| Human | 474 | SLC26A4 | 22.6 | 160.6 | 1.47 | no |
| Cow | 1073 | TRIM21 | 0.1 | 154.3 | 1.48 | no |
| Human | 1936 | SHISA5 | 69.4 | 161.5 | 1.50 | no |

| | | | | | | |
|---|---|---|---|---|---|---|
| Human | 317 | UNC93B1 | 72.2 | 161.6 | 1.50 | no |
| Human | 1916 | IFIT1 | 0.0 | 162.0 | 1.51 | no |
| Cow | 1050 | PML | 39.3 | 156.5 | 1.54 | no |
| Cow | 1175 | LAYN | 26.0 | 156.6 | 1.54 | no |
| Human | 414 | APOBEC3F | 15.2 | 163.5 | 1.54 | no |
| Macaque | 974 | GPR37 | 10.6 | 151.2 | 1.55 | no |
| Macaque | 662 | HPSE | 54.1 | 151.8 | 1.57 | no |
| Human | 516 | PSORS1C1 | 8.7 | 164.8 | 1.57 | no |
| Cow | 1087 | CMTR1 | 0.2 | 158.0 | 1.58 | no |
| Macaque | 768 | TDRD7 | 1.4 | 152.8 | 1.60 | no |
| Cow | 1227 | STARD8 | 16.3 | 159.2 | 1.61 | no |
| Macaque | 984 | XRN | 5.6 | 153.5 | 1.62 | no |
| Macaque | 829 | TRIM34 | 45.0 | 153.6 | 1.63 | no |
| Macaque | 869 | RBM25 | 6.7 | 154.7 | 1.66 | no |
| Human | 1917 | IFITM2 | 74.9 | 168.3 | 1.66 | no |
| Human | 436 | TGFB1 | 79.8 | 168.5 | 1.67 | no |
| Macaque | 989 | ABL2 | 28.2 | 154.9 | 1.67 | no |
| Cow | 1100 | TAGAP | 54.3 | 161.6 | 1.68 | no |
| Human | 268 | TDRD7 | 40.0 | 169.3 | 1.68 | no |
| Cow | 1043 | PARP12 | 1.1 | 162.4 | 1.70 | no |
| Cow | 1206 | MARCKSL1 | 57.2 | 163.0 | 1.72 | no |
| Human | 142 | PAK3 | 60.0 | 170.6 | 1.72 | no |
| Macaque | 666 | PXK | 59.7 | 157.0 | 1.73 | no |
| Cow | 1086 | TAP1 | 0.0 | 164.2 | 1.75 | no |
| Cow | 1193 | SP140L | 6.7 | 164.5 | 1.75 | no |
| Human | 282 | SLC25A28 | 55.5 | 172.2 | 1.76 | no |
| Macaque | 979 | PTAR1 | 4.0 | 158.1 | 1.76 | no |
| Human | 256 | DDX3X | 0.5 | 172.8 | 1.77 | no |
| Human | 50 | TMEM140 | 80.3 | 173.1 | 1.78 | no |
| Macaque | 594 | BST2 | 0.1 | 159.0 | 1.79 | no |
| Human | 460 | RET | 33.4 | 174.6 | 1.81 | no |
| Macaque | 665 | PXK | 35.6 | 160.0 | 1.82 | no |
| Macaque | 896 | LGALS3BP | 42.5 | 160.1 | 1.82 | no |
| Human | 164 | PARP12 | 40.9 | 175.9 | 1.84 | no |
| Human | 424 | SP100 | 77.1 | 176.5 | 1.86 | no |
| Human | 336 | C9orf91 | 91.4 | 177.6 | 1.89 | no |
| Human | 527 | RHBDF2 | 76.6 | 177.6 | 1.89 | no |
| Human | 184 | TLR7 | 63.0 | 180.0 | 1.94 | no |
| Human | 302 | ERLIN1 | 78.8 | 180.5 | 1.96 | no |
| Macaque | 811 | TAGAP | 50.8 | 166.6 | 2.02 | no |
| Human | 374 | MARCK | 41.9 | 184.0 | 2.04 | no |
| Human | 483 | PRRG4 | 89.6 | 184.4 | 2.05 | no |
| Human | 419 | B3GALNT1 | 76.2 | 185.4 | 2.08 | no |
| Human | 496 | FPR2 | 83.1 | 185.6 | 2.08 | no |
| Human | 349 | TBX3 | 75.6 | 187.3 | 2.12 | no |
| Human | 318 | TAGAP | 61.6 | 188.8 | 2.16 | no |
| Human | 151 | BTN3A3 | 66.3 | 189.6 | 2.18 | no |
| Macaque | 593 | BST2 | 0.1 | 171.9 | 2.18 | no |

| | | | | | | |
|---|---|---|---|---|---|---|
| Human | 223 | APOL3 | 50.2 | 190.4 | 2.20 | no |
| Macaque | 972 | APOBEC3B | 0.3 | 172.8 | 2.21 | no |
| Human | 316 | MCOLN | 78.8 | 196.1 | 2.34 | no |
| Human | 70 | TMEM51 | 91.3 | 196.6 | 2.35 | no |
| Macaque | 760 | DDX3X | 0.5 | 177.4 | 2.35 | no |
| Human | 111 | TRIM14 | 73.4 | 200.4 | 2.44 | no |
| Macaque | 813 | TAGAP | 59.3 | 180.5 | 2.44 | no |
| Human | 441 | MORC3 | 51.4 | 202.3 | 2.49 | no |
| Cow | 1047 | PLEKHA4 | 0.0 | 192.5 | 2.52 | no |
| Cow | 1105 | TRIM34;TRIM | 0.0 | 192.7 | 2.52 | no |
| Cow | 1228 | STARD8 | 14.4 | 193.4 | 2.54 | no |
| Macaque | 704 | CDKN1A | 49.9 | 187.9 | 2.67 | no |
| Macaque | 622 | SLFN12L | 17.8 | 189.7 | 2.72 | no |
| Human | 544 | VCPIP1 | 0.0 | 215.7 | 2.81 | no |
| Macaque | 978 | NEXN | 0.0 | 196.2 | 2.92 | no |
| Human | 196 | CDKN1A | 69.7 | 221.7 | 2.96 | no |
| Human | 416 | GJD3 | 71.5 | 222.0 | 2.97 | no |
| Human | 1932 | ZC3HAV1 | 0.2 | 227.1 | 3.09 | no |
| Macaque | 695 | TLR7 | 24.6 | 207.0 | 3.25 | no |
| Human | 241 | WARS | 0.0 | 239.9 | 3.40 | no |
| Human | 84 | SLFN12L | 0.1 | 276.9 | 4.30 | no |
| Cow | 1002 | EIF2AK2 | 0.0 | 259.4 | 4.34 | no |
| Cow | 1137 | NUB1 | 0.0 | 288.7 | 5.14 | no |
| Cow | 1149 | SLFN11 | 0.0 | 288.7 | 5.14 | no |
| Cow | 1203 | TRANK1 | 0.0 | 288.7 | 5.14 | no |
| Macaque | 562 | EIF2AK2 | 0.0 | 276.5 | 5.36 | no |
| Human | 356 | DDX60 | 0.0 | 324.0 | 5.45 | no |
| Cow | 1211 | CDADC1 | 0.0 NA | NA | | no |
| Cow | 1148 | TMEM106A | 0.0 NA | NA | | no |
| Cow | 1276 | ZNF324 | 0.0 NA | NA | | no |
| Cow | 1183 | NLRC5 | 0.0 NA | NA | | no |
| Cow | 1163 | CASP8 | 0.0 NA | NA | | no |
| Human | 469 | ARHGAP27 | 0.0 NA | NA | | no |
